# Fluorescence lifetime analysis of smFRET with contribution of PIFE on donor and acceptor

**DOI:** 10.1101/2023.04.03.535482

**Authors:** Sina Jazani, Taekjip Ha

## Abstract

Single-molecule fluorescence resonance energy transfer (FRET) is a powerful technique based on dipole-dipole interaction between donor and acceptor fluorophores to observe inter- and intra-molecular dynamics in realtime with sensitivity to macro-molecular distances (∼ 2.5-10 nm). That said, some fluorophores have an inherent characteristic known as protein induced fluorescence enhancement (PIFE). PIFE is a photo-physical feature of dyes undergoing cis-trans transitions and occurs for protein-dye interactions closer than 3 nm. Here, the challenge is uncoupling the PIFE effect in the FRET data. Ignoring the PIFE effect in the analysis of the FRET data may lead to misinterpretation of the system under investigation. As a solution to this problem, we develop a computational framework based on Bayesian statistics to analyze the fluorescence lifetime signals of the donor and acceptor channels which allows us to uncouple the PIFE effects from the FRET. Our framework can extract any changes in the FRET efficiency simultaneously with any changes in the fluorescence lifetimes of the donor and acceptor due to the PIFE effect. In addition, our framework can provide other parameters, such as the donor and acceptor excitation rates, background photon rates, and detectors’ cross-talk ratios. Our framework extracts all these parameters by analyzing a single photon arrival time trace with only a few thousand photons.

---

Fluorescence resonance energy transfer (FRET) is a technique that allows us to capture the dynamic behaviors of molecules at the length scale of a few nanometers. FRET is used to measure whether two molecules are in close proximity, or determine the distance between two specific locations on macromolecules or in molecular complexes.^1–3^ FRET is a physical process by which the excited state energy of one fluorophore, the “donor”, can be transferred nonradiatively to a neighboring fluorophore, the “acceptor”. This occurs when the two molecules are close enough, usually separated by less than 10 nm. The development of single-molecule fluorescence spectroscopy^4–6^ in combination with FRET has enabled us to directly visualize conformational changes of a single molecule or complex. Also, mechanisms underlying reactions, e.g., protein-nucleic acid and protein-protein interactions, can now be observed in real time.^4,7–19^ Single-molecule FRET (smFRET) has been used to answer fundamental questions about replication, recombination, transcription, translation, RNA folding and catalysis, non-canonical DNA dynamics, protein folding and conformational changes, various motor proteins, membrane fusion proteins, ion channels, signal transduction, to name just a few with this list growing rapidly.^18,20–24^

FRET signals must be interpreted carefully in order to avoid the effect of other photo-physical events on the results.^22^ One of the well-known photo-physical events is protein induced fluorescence enhancement (PIFE) which can be observed in the case of using fluorophores such as Cy3, Cy5, and DY547.^25,26^ PIFE is due to the presence of a deactivation pathway in the first excited state of the fluorophore that leads to a radiationless transition, the cis-trans isomerization of an exocyclic double bond.^27–29^ PIFE is dye-protein interaction dependent which leads to a change in the intensity of the fluorophore upon conjugation to protein.^25–27^ PIFE can be designed in such a way to capture fast dynamics over short distances below 3 nm where thFRET is less sensitive.^25,26,30^

Both FRET and PIFE not only lead to changes in the intensity of the fluorophore but also lead to changes in the fluorescence lifetime. Several studies showed a high correlation between the lifetime and intensity.^25,31^ There are several studies trying to formulate the underlying process of the FRET in the presence of the PIFE.^27,32–34^. The summary of these studies can be seen in Fig. 1(b) where we have extra transitions in the case of PIFE for both donor and acceptor. Here, the excited donor in the trans state may take different paths to escape its excited state such as radiative decay to its ground state with the rate 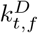, nonradiative decay to its ground state with the rate 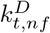, energy transfer to the excited state of the acceptor with the energy transfer (ET) rate *k*_*F*_, or nonradiative decay to the ground state of the cis configuration with the rate 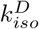. This means that the observable escape rate of the donor from its excited state is 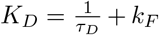 where 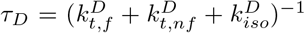 is the lifetime of the donor independent of the FRET. Here, the *k*_*F*_ changes based on the distance between the donor and acceptor (Fig. 1(a)). In this study, we are considering discrete changes in the *k*_*F*_ through time, and we are calling these discrete levels of ET rate, the FRET state. Also, *τ*_*D*_ de-pends on the isomerization rate 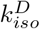 which changes based on the distance between the dye and protein (Fig. 1(c)). So, by assuming constant 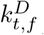 and 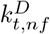 during the ex-periment, we are able to consider the PIFE effect as the main source of any changes in the lifetime of the donor. In this study, we are considering discrete changes in donor lifetime through time, and we are calling these discrete levels of the donor lifetime, the donor-PIFE state.

**FIG. 1.**
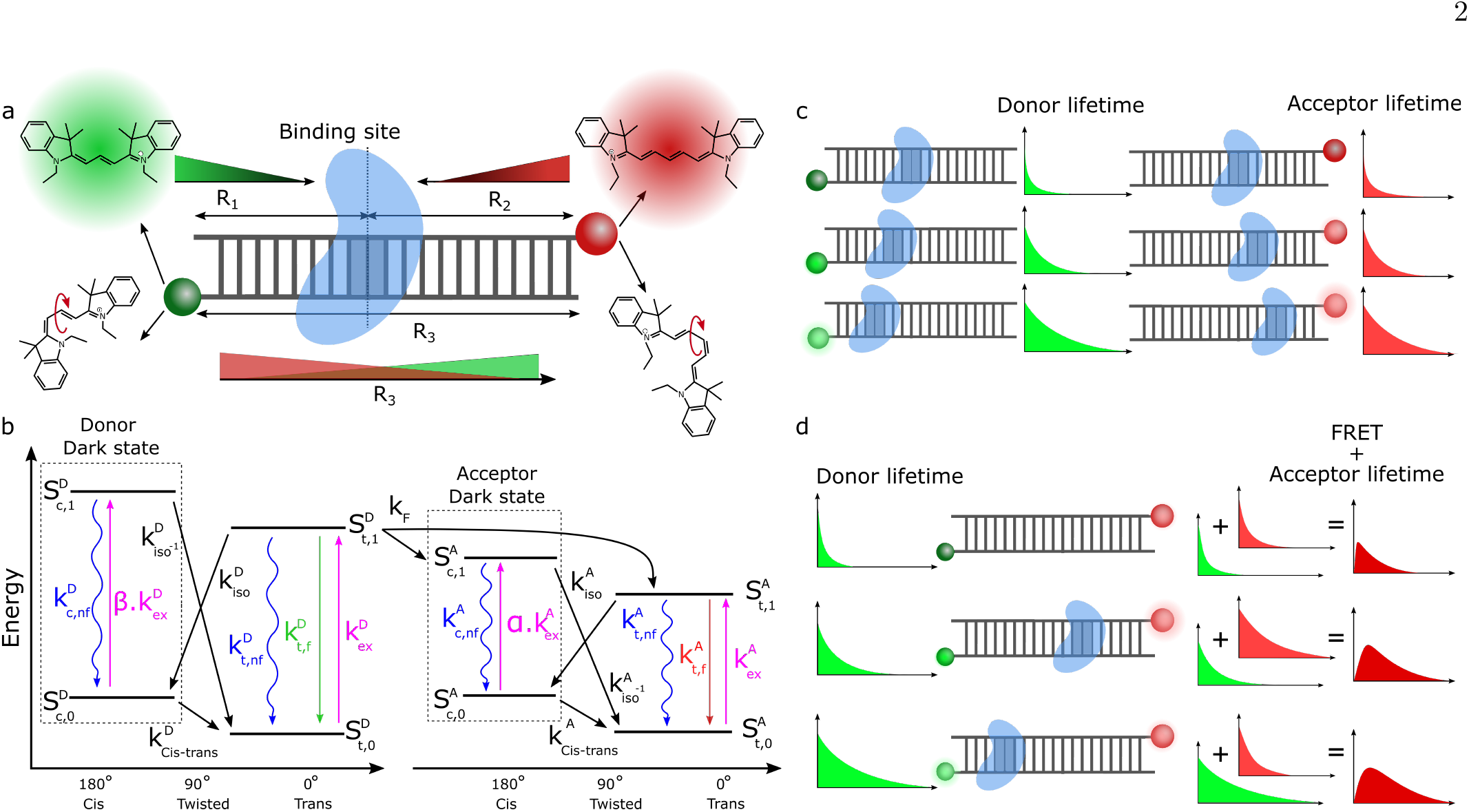
Schematic of the photophysics of Cy3 as the donor and Cy5 as the acceptor depending on the FRET and PIFE. **(a)** A cartoon representation of the cis-trans transitions of Cy3 and Cy5 depend on the distance of the protein from the Cy3, *R*_1_ and the distance of the protein from Cy5, *R*_2_. At the same time, the energy transfer rate (ET rate) depends on the distance between Cy3 and Cy5, *R*_3_. **(b)** The Jablonsky diagram of the cis-trans transition of Cy3 and Cy5, and FRET. In this case, we assume that there is no fluorescence photon emission from cis states. Here, there are many possible transitions, but in our model, we only consider the transition which leads to photon emission. This simplification reduces our calculations. **(c)** A cartoon illustration of the PIFE effects on Cy3 and Cy5. As the distance between the protein and dyes decreases (*R*_1_, *R*_2_), the fluorophore is going to stay more in the trans state and consequently, the fluorescence intensity and fluorescence lifetimes will increase. Here, we show fluorescence lifetimes with the photon arrival time distributions. **(d)** A cartoon visualization of the combination of FRET and PIFE. Here, the distribution of photons collected in the donor channel is going to be affected by the PIFE and FRET. Also, the distribution of the photons collected in the acceptor channel is a result of the convolution of two processes: first, donor excitation and resonance energy transfer; and second, emission of the acceptor.

In the case of the acceptor, there is no energy transfer from the excited state of the acceptor to the excited state of the donor. So, the excited acceptor in the trans state may take different paths to escape its excited state such as radiative decay to its ground state with the rate 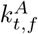, nonradiative decay to its ground state with the rate 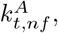, or nonradiative decay to the ground state of the cis configuration with the rate 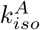. As a result, the observable escape rate of the acceptor from its excited state is 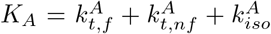. This means any changes in the lifetime of the acceptor 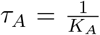 are caused by the PIFE effect. Here, we are considering discrete changes in acceptor lifetime through time, and we are calling these discrete levels of the acceptor lifetime, the acceptor-PIFE state.

In the event of direct excitation of the acceptor, the distribution of the photon arrival times collected in the acceptor channel follows an exponential decay with a lifetime of *τ*_*A*_ (Fig. 1(c)) while in the case of FRET + PIFE (Fig. 1(d)), the change in donor lifetime is due to a competition between FERT and the isomerization of the dye to the cis state. In this scenario, the distribution of photon arrival times collected from the donor channel follows an exponential decay with a lifetime of 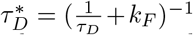 and the distribution of the photon arrival times collected in the acceptor channel follows the convolution of two exponential decay distributions with lifetimes of 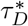 and *τ*_*A*_.

In the case that PIFE presents simultaneously with FRET, any changes in the observable donor lifetime 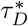 may be due to changes in the rates *k*_*F*_, 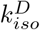, or both. This coupling causes a problem in measuring the absolute values of the FRET rates and donor lifetimes. As a solution to this problem, we developed a computational framework based on Bayesian statistics^35–39^ to uncouple FRET and PIFE. To do so, we focus on the experimental setups involving pulsed illuminations and single photon detectors (Fig. 2) which output the photon arrival times with respect to the immediate previous excitation pulse, called (micro time)^35,40^ and the collection of photon arrival times (macro time) starting from the beginning of the experiment.^41^ The macro time traces can be transformed to the intensity time traces while the micro time traces can be used to measure the fluorescence lifetimes using the time correlated single photon counting (TCSPC) histogram^31^, phasor-based approaches^42,43^ or direct analysis of the individual photons^35,36^ in order to capture the lifetime(s)^31,44^.

**FIG. 2.**
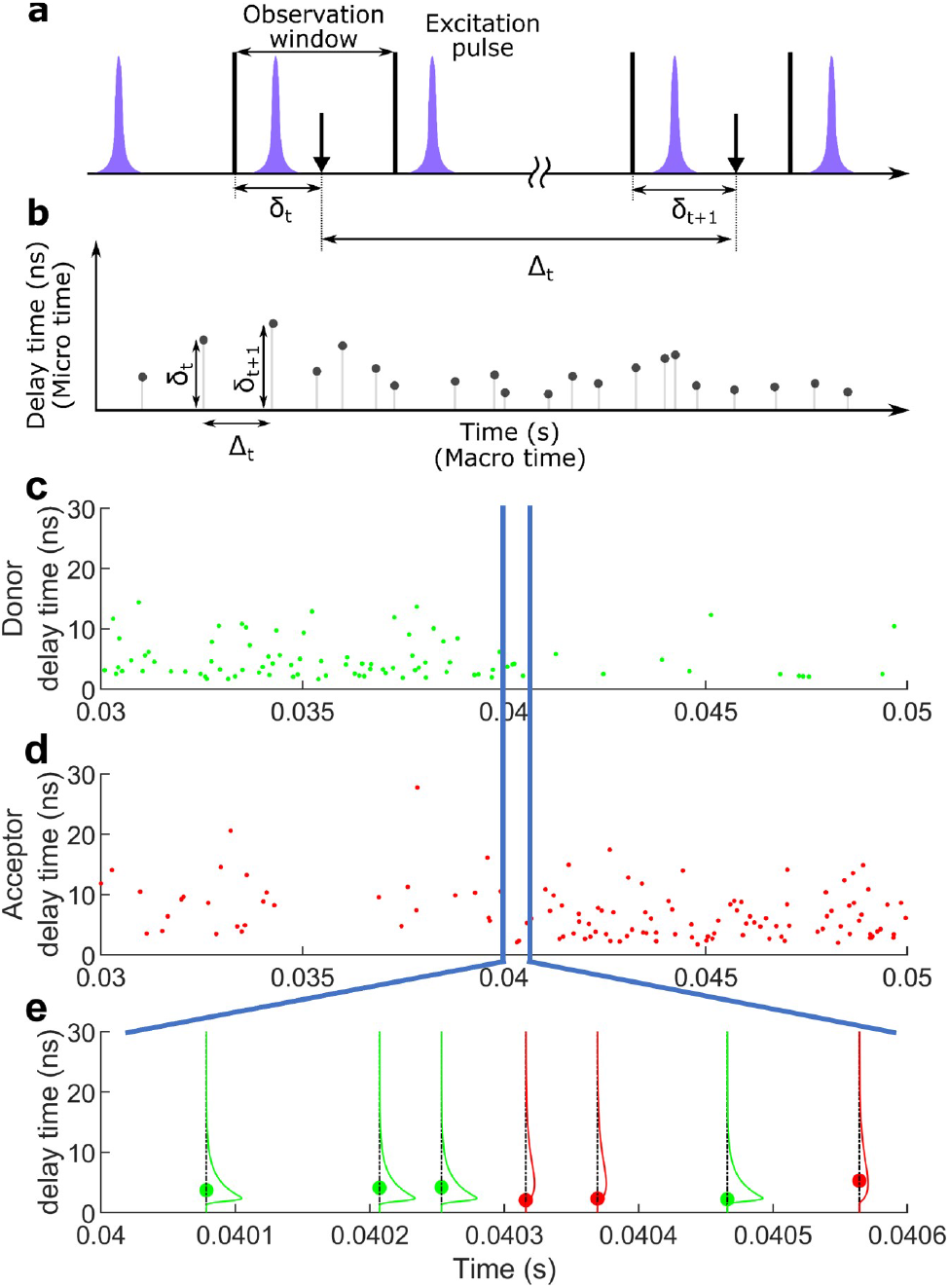
Schematic of the photon collection with a single photon detector and distributions of photon delay times collected in the donor and acceptor channels. **(a)** A cartoon representation of an array of excitation pulses and the photon collection times. **(b)** The concept of single photon arrival time traces in two time-dimensions of micro and macro times. **(c-d)** An example of single photon time traces collected by the donor and acceptor channels. **(e)** Magnified portion of time traces in (c & d) with an illustration of the detected photons and their distributions.

The core of our approach is based on the classification of the detected photons with different emissions. These sources are: (i) donor excitation and emission; (ii) donor excitation and acceptor emission due to the FRET; (iii) direct acceptor excitation and emission; (iv) donor background; and (v) acceptor background. In the cases of detected photons with background sources, we assume a uniform photon collection rate for the background in donor and acceptor channels. In the cases of detected photons with donor or acceptor excitation sources, our classification approach can be explained by separating excitation and emission. In the case of excitation, we assume donor and acceptor excitation rates may vary based on their PIFE states.^32^ This is due to the fact that the dye in the cis state is dark, and the amount of time that dye stays in the cis state or the trans state affects the total intensity of the dye. In the case of emission, as we discussed above, each scenario requires a different model with fluorescence lifetimes depending on the FRET and PIFE states. Our analysis relies on the fluorescent lifetimes of individual detectable photons. As a result, since there is no radiative transition from the excited cis state, we safely ignored all transitions within or from the cis states in our analysis.

Here, we take advantage of the information carried by individual detected photos with single-photon interarrival temporal resolution by directly analyzing both macro and micro time traces. This allows us to capture discrete changes in *k*_*F*_ or *τ*_*D*_ and *τ*_*A*_ due to the PIFE effect with high temporal resolution. In this study, we applied the Hidden Markov model (HMM)^45–58^ to capture any discrete changes in FRET efficiency, donor-PIFE and acceptor-PIFE with a resolution of single-photon arriving times. This means we have three hidden state trajectory traces with Markovian properties, one for FRET, one for donor-PIFE, and one for acceptor-PIFE.

## I. RESULTS

### Overview

The goal of this study is to analyze a multi-channel single-photon arrival time trace to capture instantaneous changes in ET rate *k*_*F*_, changes in donor lifetime *τ*_*D*_ caused by the PIFE effect (donor-PIFE), and changes in the acceptor lifetime *τ*_*A*_ due to the PIFE effect (acceptor-PIFE). We utilized Bayesian statistics^37–39^ in order to estimate the parameters of interest. According to Bayes’ theorem, we have the relationship between the likelihood and the posterior as

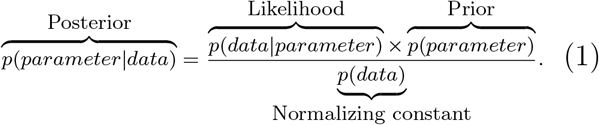

Using the Bayesian model allows us to access the posterior probability distribution of each parameter of interest. Unlike point estimation approaches such as the maximum likelihood estimation (MLE)^59^ that find parameters related to the maximum likelihood probability distribution, our outputs are the probability distributions. The widths of these distributions represent the errors of our parameter estimations. Also, as a comparison to point estimation approaches we include the maximum a posteriori

(MAP), the most probable values, plus the 95% confidence interval as the error of our estimations. The MAP estimate should be equal to MLE with the assumption of a negligible prior effect on the posterior. ^37–39^

To demonstrate the introduced approach, we consider scenarios with multiple FRET states *m*_*F*_ = 1, …, *M*_*F*_, donor-PIFE states *m*_*D*_ = 1, …, *M*_*D*_ and acceptor-PIFE states *m*_*A*_ = 1, …, *M*_*A*_. Our method outputs the ET rates 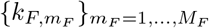, donor lifetimes corresponding to donor-PIFE states 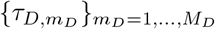, and ac-ceptor lifetimes corresponding to acceptor-PIFE states 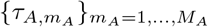. Here, we assume that all transitions for the FRET and PIFE are discrete and as a result, we can estimate the FRET state transition rate matrix 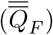, donor-PIFE state transition rate matrix 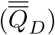, and acceptor-PIFE state transition rate matrix 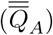. The transition rate matrices can be written in the form of

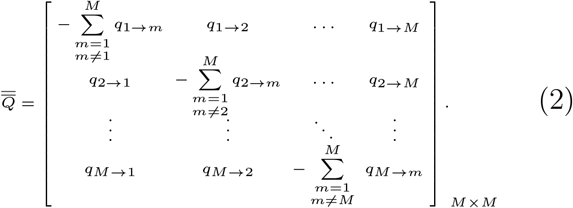

### Validation with simulated data

To validate our method, we generated synthetic time traces based on donor-only pulse excitation with the donor and acceptor detection channels recording single-photon micro and macro time traces. The synthetic time traces are generated based on two different sets of parameter configurations: (i) transitions between two molecular conformations (two FRET states) without any PIFE, Fig. 3; (ii) transitions between multiple molecular conformations (multiple FRET states) while the donor and acceptor lifetimes change due to PIFE, Fig. 4. In all figures, we show the results in the form of MAP estimate and the 95% confidence interval as the uncertainty introduced by the finiteness of the data and the inherent noise. To evaluate the precision of our method, we compared the results with the ground truth values. A summary of results for all parameters is shown in Supplementary Tables S1 and S2.

**FIG. 3.**
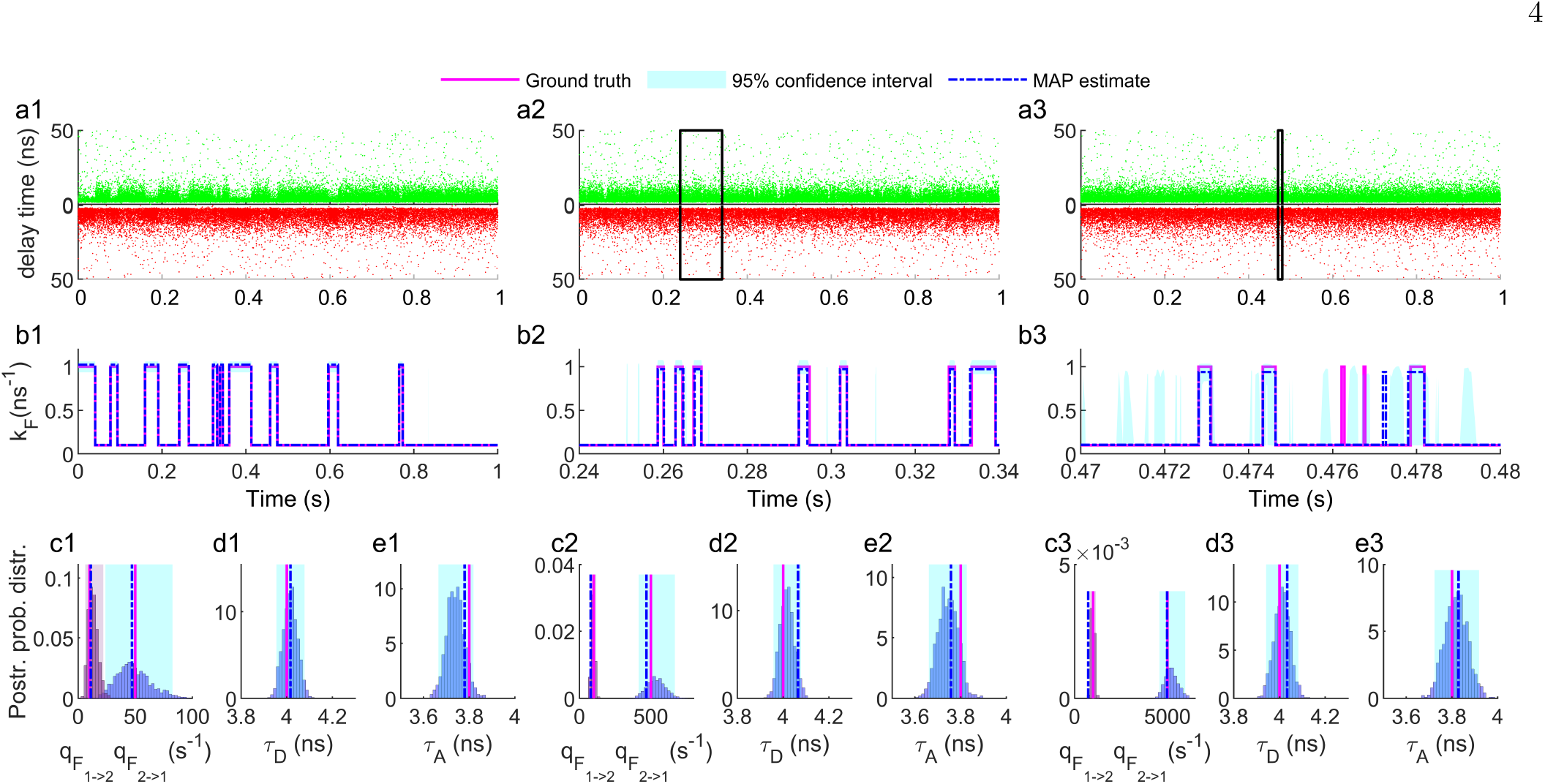
Evaluation of the method based on the synthetic time traces with different FRET state transition rates. **(a1-3)** Synthetic single-photon time traces generated with different FRET state transition rates between two FRET states [*qF*_1→2_, *qF*_2→1_ ] of [10, 50], [100, 500] and [1000, 5000] s^−1^. The ET rates (*k*_*F*_), are 0.1 and 1 ns^−1^ for lower and higher FRET states. Each trace contains 5×10^4^ photons with donor and acceptor lifetimes of 4 and 3.8 ns, respectively. The background photon emission rates are 500 photons s^−1^ for each donor and acceptor channels and cross-talks between channels are 10%. **(b1-3)** Estimations of ET rate trajectories by analyzing traces shown in (a1-3). For visual purposes, panels (b2) and (b3) are showing the results of our analysis for the portions of the time traces shown with black boxes in (a2) and (a3), while we analyzed the whole time traces. **(c1-3)** The results of our analysis for the transition rates between FRET states, are shown in the form of distributions with highlighted 95% confidence intervals. **(d1-3)** The results of our analysis for the donor fluorescence lifetimes. **(e1-3)** The results of our analysis for the acceptor fluorescence lifetimes.

**FIG. 4.**
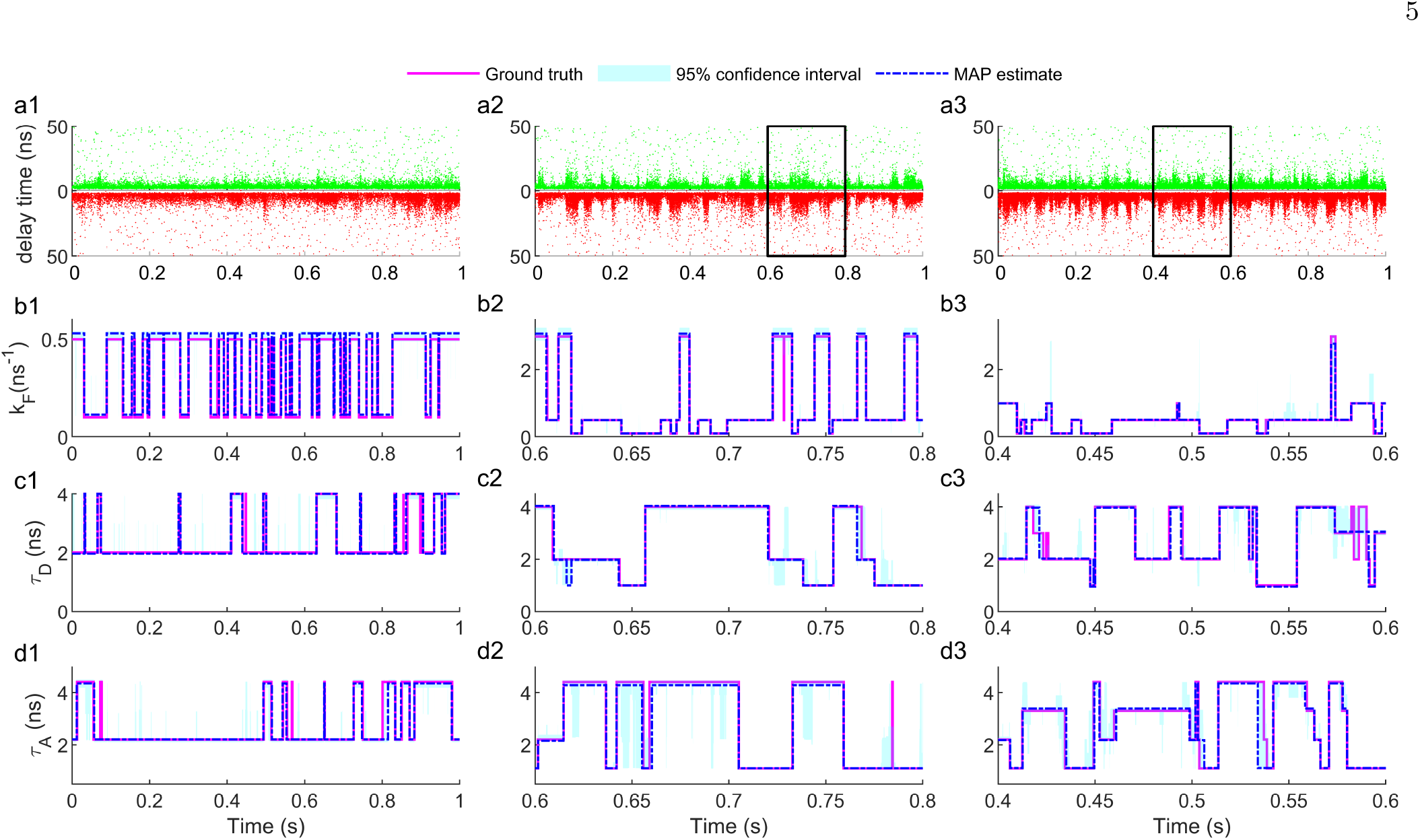
Evaluation of the method based on synthetic time traces with multiple FRET states in presence of PIFE effects on donor and acceptor dyes. **(a1-3)** Synthetic single-photon time traces generated with FRET transitions between two, three and four FRET states including donor-PIFE with two, three and four different states (donor lifetimes), and acceptor-PIFE with two, three and four states (acceptor lifetimes). These time traces contain the total number of photons of 2×10^4^, 5×10^4^ and 10×10^4^ with the same background photon rates of 500 photons s^−1^ for each channel, and 10% cross-talks between channels. **(b1-3)** Estimations of FRET state trajectories by analyzing traces shown in (a1-3). For visual purposes, panels (b2) and (b3) are showing the results of our analysis for the portions of the time traces shown with black boxes in (a2) and (a3), while we analyzed the whole time traces. **(c1-3)** Estimations of donor-PIFE state trajectories representing changes in donor lifetime by analyzing traces shown in (a1-3). The colors are assigned the same as the plots in (b) panels. **(d1-3)** Estimations of acceptor-PIFE state trajectories representing changes in acceptor lifetime by analyzing traces shown in (a1-3). The colors are assigned the same as the plots in (b) panels. Also, other parameters such as state transition rates between the states are reported in the Supplementary Table S1.

To test if our method can capture fast state transitions, we generated simulated data with various transition rates for a system with two discrete FRET states. In Fig. 3(a1-3), we illustrate three synthetic time traces as the collection of single-photon arrival times from both donor and acceptor channels. Here, the y-axis represents the arrival time of photons after excitation pulse (micro time) and the x-axis represents the photon detection starting from the beginning of the experiment (macro time). The total number of photons for Fig. 3(a1-3) are 5×10^4^ photons while the transition rates between FRET states [*qF*_1→2_, *qF*_2→1_], are [10, 50], [100, 500], [1000, 5000] s^−1^. So, the average number of photons per dwell in each FRET state is [5000, 1000], [500, 100] and [50, 10]. In Fig. 3(b1-3), we show the output of our model in the form of the FRET state trajectory of the molecules at any given time in comparison with the ground truth.

In Fig. 3(b1-3), the estimations of the state trajectories over time for time traces with higher state transition rates are worse than those with lower rates. This increase in error was expected since fewer photons per dwell are collected for the time traces with higher state transition rates. Despite the increasing error, it is clearly possible to capture FRET transitions with only tens of photons per dwell by relying on the single-photon lifetime analysis. Also, as we show in Fig. 3(c1-3), Fig. 3(d1-3) and Fig. 3(e1-3), our model is able to estimate the FRET state transitions rates [*qF*_1→2_, *qF*2→1 ], donor and acceptor fluorescence lifetimes (*τ*_*D*_, *τ*_*A*_) in the absence of energy transfer. As we mentioned before, these lifetimes are independent from FRET effect on the apparent lifetimes 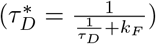.

Next, we evaluate our model on systems with multiple FRET states and multiple PIFE states for both donor and acceptor (Fig. 4). We consider donor and acceptor lifetimes changing through time (due to PIFE) for a bio-molecule undergoing conformational changes between FRET states. We also studied different numbers of FRET, donor-PIFE and acceptor-PIFE states to evaluate the capability of our method on systems with higher complexities (higher number of states). The results of our method are estimations of resonance energy transfer rates (changes due to FRET), donor lifetimes (changes due to the donor-PIFE), and acceptor lifetimes (changes due to the acceptor-PIFE) at any given time that we detected a photon (Fig. 4).

By increasing the number of states, the error of our es-timations rises due to the higher complexity of the data. For instance, in the case that we have two FRET states, two donor-PIFE states, and two acceptor-PIFE states, the number of different lifetimes for the donor is 2×2, 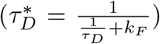, while the acceptor has two lifetimes. It means that there is a total of 6 lifetimes to estimate for this data set (Fig. 4(a1)). So, increasing the number of FRET, donor-PIFE and acceptor-PIFE states to 3 states for each, leads to 12 lifetimes (Fig. 4(a2)), and 4 states for each, leads to 20 lifetimes (Fig. 4(a3)). Besides the complexity due to the number of lifetimes, by increasing the number of states (FRET or PIFE), the size of the transition rate matrices increase which means more number parameters to estimate. For example, for the time trace in Fig. 4(a1) we have three transition rate matrices each with the dimension of 2×2, and in total, there are 6 parameters to calculate while for Fig. 4(a2) their dimensions increase to 3×3 and there are 18 parameters to calculate, and in Fig. 4(a3) this increases to 4×4 and 36 parameters. In order to increase the accuracy and precision of our estimations, we analyzed time traces with a higher number of photons for systems with higher complexity (higher number of FRET and PIFE states).

### Validation with experimental data

We were able to validate our method using experimental data by showing the capability of our method to estimate the ET rates, FRET state transition rates, donor and acceptor fluorescence lifetimes and compare them with the outputs of previously established methods^60^ Here, we analyzed experimental single molecule fluorescence time traces based on three different dye-labeled proteins, *α*3D, gpW, and the WW domain. In these experiments, *α*3D and gpW both were labeled with Alexa 488 and Alexa 594, and WW domain was labeled with Alexa 488 and Atto 647N as the donor and acceptor.^60^ These dyes do not have any reported cis-trans transitions, therefore no PIFE effect^25,26^ making them a perfect system to validate our approach, especially on the FRET state trajectories with different transition rates. Also, as previously reported, our biological system is well defined^60^, and we expect to see only two molecular conformations for these proteins. Here, we have access to the reported, transition rates between FRET states, FRET efficiencies, and donor and acceptor lifetimes, and we use these values as the ground truth to validate our method.

Here, we illustrate representative time traces for *α*3D, gpW, and WW domain, each containing ∼ 2 ×10^4^ photons collected from donor and acceptor channels, combined (Fig. 5(a1-3)). The results of our analysis are in form of ET rates (*K*_*F*_), transition rates between FRET states, and donor and acceptor lifetimes (Fig. 5(d1-3) and Fig. 5(e1-3)).

**FIG. 5.**
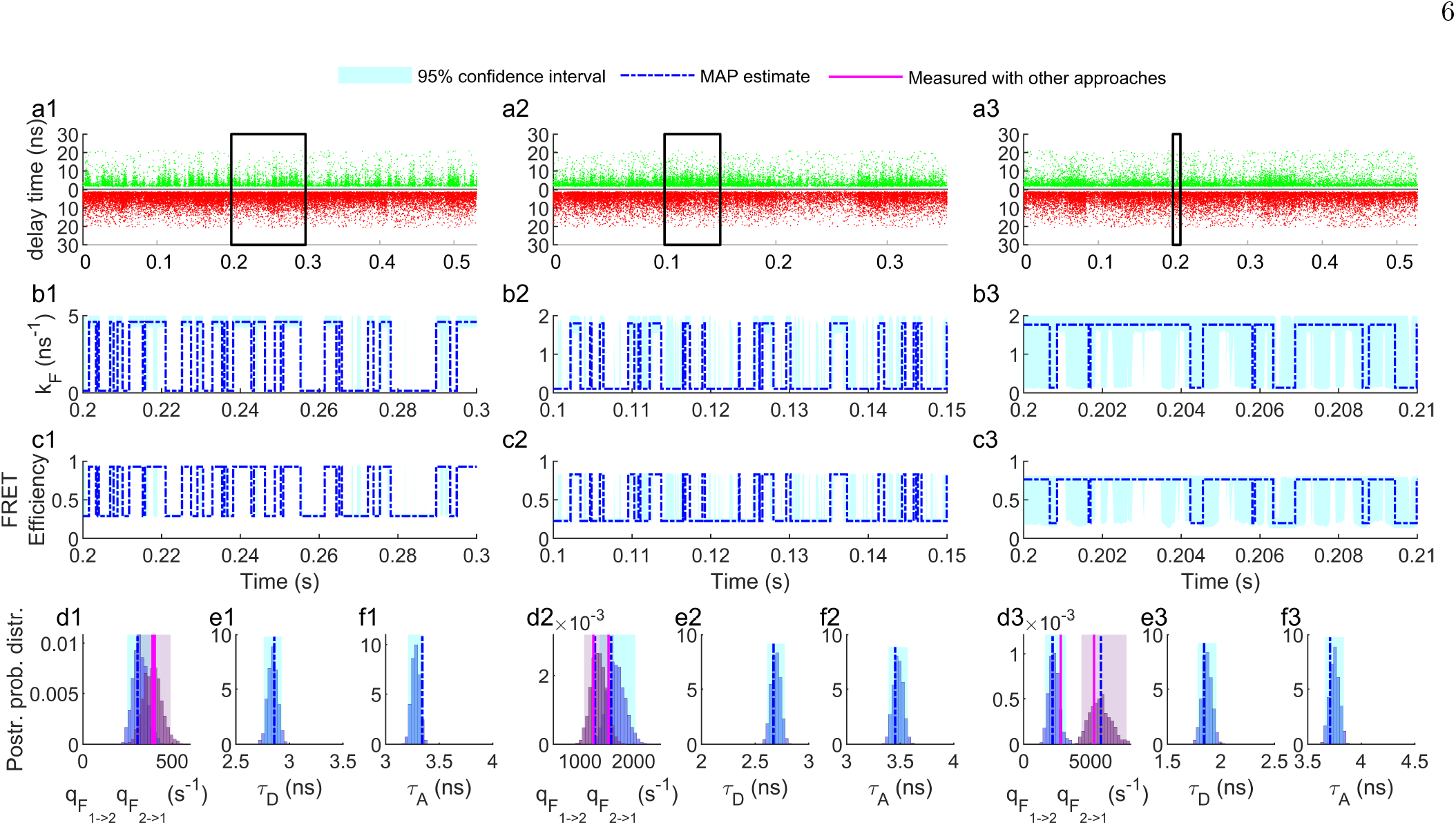
Evaluation of our method with experimental time traces with three bio-molecules of *α*3D, gpW, and WW domain with different transition rates between FRET states. **(a1-3)** Experimental single-photon arrival time traces of *α*3D, gpW, and WW domain collected with donor and acceptor channels. Each time trace contains multiple individual collected data sets visualized as one time trace with approximately 2×10^4^ photons in total. **(b1-3)** Estimations of FRET state trajectories (rate changing through time) by analyzing traces shown in (a1-3). **(c1-3)** Estimations of FRET state trajectories (FRET efficiency changing through time) by analyzing traces shown in (a1-3). **(c1-3)** The results of our analysis for the transition rates between FRET states are shown in the form of distributions with highlighted 95% confidence intervals and the most probable output of our analysis (MAP) versus the measured values by other approaches. **(d1-3)** The results of our analysis for the lifetimes of the donor. **(e1-3)** The results of our analysis for the lifetimes of the acceptor.

These results clearly show the capability of our approach to estimating discrete changes of ET rate *k*_*F*_ for very fast dynamics like the WW domain (*∼* 5000 s^−1^), although the error increased for such fast dynamics due to the small number of photons per dwell [15, 6]. The immediate solution to reduce this error is to increase the laser power to increase the excitation rate of the donor and consequently collect more photons per unit of time. Collecting additional photons means more photons per dwell and consequently reduces the error.

Although we analyze only a few thousand photons, our results are comparable to the reported values in the literature (> 10^5^).^31,60^ Also, we evaluate the accuracy and precision of our estimations as the outputs of our method based on the time traces with different numbers of photons. and it can be found in Supplementary Fig. S1 and Fig. S2.

As an additional experimental validation, we suggest applying our method for labeled molecules with donor and acceptor dyes that show the PIFE effect. In this case, we expect to have donor-PIFE, acceptor-PIFE and FRET at the same time. This experiment may be challenging since access to the ground truth is limited. So, instead, the previously published estimations of the FRET efficiency, and fluorescence lifetimes of donor and acceptor, can be used to evaluate the output of our method. As an example of such experiments, we can recommend a PcrA Helicase at 5^*′*^ partial duplex junction while the Cy3 and Cy5 are located at the end tail and partial duplex junction, respectively.^61^ This system will allow us to observe the PIFE effect while the ET rate is changing due to the translocation of the ssDNA from 3^*′*^→ 5^*′*^.^61^

## II. CONCLUSION

In this research, we present a Bayesian statistical approach for analyzing multi-channel fluorescence lifetime microscopy data. Our approach provides a way to uncouple the FRET and PIFE effects by analyzing time traces with a relatively low photon count. It simultaneously measures FRET efficiency, donor lifetime, acceptor lifetime, and changes in these parameters over time. This approach allows us to reduce the laser intensity and consequently minimize the bleaching rate and photo-damage to the sample.^62,63^ Additionally, it is less sensitive to donor or acceptor blinking effects since it relies on the individual detected photons instead of changes in intensity.

In this study, we expand on other studies (they have shown that fluorescence signals can be analyzed using only the single photon arrival time information.^31,44^) by mathematically reformating their approach to simultaneously analyze the donor and acceptor signals. Therefore, by honing in on variations in single photon arrival times, our method is capable of detecting the PIFE effect, which manifests in changes in fluorescence lifetimes, as well as measuring FRET. The PIFE data obtained through our approach can be used to observe fast molecular dynamics happening below *∼*2.5 nm (sensitivity limit of the FRET)^25,26,30^ and detect conformational changes of the targeted bi-molecule or binding-unbinding, instability of a protein at the target site, etc.^30,64^

Current state-of-the-art approaches rely on analyzing the intensity time traces or a marginal effect of the single-photon time traces. However, these methods have limitations since they require a large number of photons and cannot accurately track rapid changes in donor and acceptor lifetimes in the presence of FRET.^25,26,65^ Our approach, on the other hand, uses single photons, enabling us to simultaneously estimate FRET and PIFE state trajectories with exceptional time resolution, as low as single photon arrival time of *∼*10μs. Moreover, the information contained within each set of single photon arrival times is unique and dependent on the system’s underlying configuration.

We have implemented a Bayesian approach to our model, providing a robust estimation of error. Unlike other metrics such as residuals (i.e., chi-square), our method considers the content and length of the single photon arrival time traces, allowing for more precise measurements and an accurate assessment of the width of the posterior probability distribution.^66,67^

Our approach, which relies solely on measuring the fluorescence lifetimes of the donor and acceptor, has a broad range of applications. It is particularly useful in cases where molecular conformations undergo continuous transitions rather than discrete ones, such as in DNA translocations,^61^ or small deviations before or after conformational changes, such as delay time in protein relaxation^68^ or drift corrections.^69^

To address such scenarios, we propose a solution where we combine our model using a Hidden Markov Model (HMM) with a continuous model, such as the Gaussian process approach.^70,71^ By doing so, we can completely or partially separate the discrete dynamics from the continuous ones, providing a comprehensive and precise understanding of the system’s behavior. To address the issue of super-fast dynamics that occur below the data acquisition time (single-photon arrival times), we propose extending our model by applying the Hidden Markov Jump Process (HMJP) to capture these rapid transitions.^72^

Finally, since we are using Bayesian approaches, the scalability challenge is an important consideration due to the high computational cost. To improve computational efficiency, we can analyze portions of time traces in a parallel manner, which will increase computational time linearly with respect to the size of the time traces and available CPU cores on the computer. Alternatively, we can use Bayesian-neural networks to speed up the model, making it applicable for high throughput analysis.^73–76^ These approaches offer solutions for improving scalability and reducing computational time, allowing for the analysis of large datasets with high efficiency.

## III. METHOD

### Model overview

Here, as the observation trace, we consider a multichannel fluorescence lifetime time traces which is the collection of micro times and single photon arrival times after the excitation pulses, 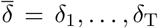. As the result of analyzing such traces we estimate the parameters include: (i) ET rates related to each FRET state of the molecule (e.g. fold and unfold), 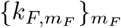; (ii) FRET state trajectory of a single molecule, 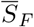; (iii) FRET state transition rate matrix, 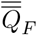; (iv) fluorescence lifetimes of the donor at each state of the molecule (in the case of PIFE effect), 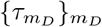; (v) PIFE state trajectory of the donor, 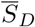; (vi) donor-PIFE state transition rate matrix, 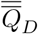; (vii) fluorescence lifetimes of the donor at each state of the molecule (in the case of PIFE effect), 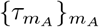; (viii) PIFE state trajectory of the acceptor, 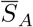; (ix) acceptor-PIFE state transition rate matrix, 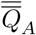; (x) detector cross-talks, *η*_*A*_ and *η*_*D*_; and, (xi) background photon rates, *μ*_back,*A*_ and *μ*_back,*D*_. Also, there are auxiliary variables such as excitation rates of donor 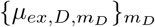and the acceptor 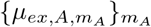 which more information about them can be found in the Supplementary Materials Sec. S2 and S3. In this chapter, we explain each one of the mentioned parameters in detail and the computational and methodology in detail can be found in Supplementary Materials Sec. S4. The summary of the notation, abbreviations, and mathematical definitions can be found in Supplementary Tables S3 and S4.

### Model description

In our model, the likelihood contains two parts. First, the detection channel, 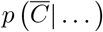 and the second part of the likelihood “the micro time”, 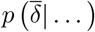 In or-der to include the case of acceptor excitation, we defines some weights over the source of the photon t, 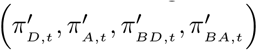, which might comes from the excited donor, excited acceptor, donor background or the acceptor background, respectively. These weights are functions of background photon rates of acceptor and donor channel *μ*_back,*A*_, *μ*_back,*D*_, acceptor and donor excitation rates 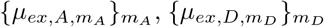, and the PIFE states of the acceptor and donor dyes *S*_*A,t*_, *S*_*D,t*_. Also, in order to take into account the cross talk between the channels, we consider the cross-talk ratios for each channel *η*_*A*_ and *η*_*D*_. More information can be found in Supplementary Materials Sec. S2 and S3.

As the result, the first part of the likelihood, the detection channel can be written as

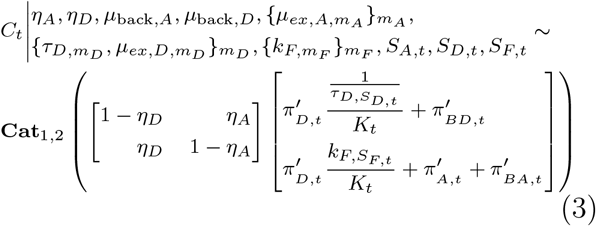

where *C*_*t*_ = 1 represents the detection of the photon in the donor channel and *C*_*t*_ = 2 is equivalent to the detection of the photon in the acceptor channel. The 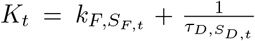. Then, the second part of the likelihood can be written as

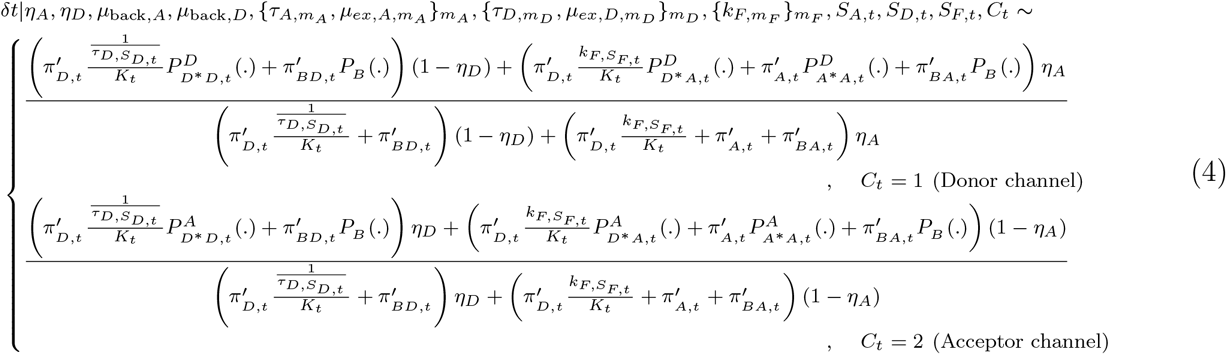

where, the photon arrival time distributions are: 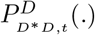donor excitation and emission collected in donor channel; 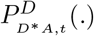 donor excitation and accep-tor emission due to FRET collected in donor channel; 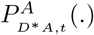 donor excitation and acceptor emission due to FRET collected in acceptor channel; 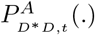 donor excitation and emission collected in acceptor channel; 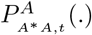 acceptor excitation and emission collected in acceptor channel; 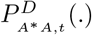 acceptor excitation and emis-sion collected in donor channel; and *P*_*B*_ (.) is the uniform background photon arrival time during the pulses periods. A detailed description of these distributions can be found in the Supplementary Materials Sec. S1.

In order to consider the errors caused by the instrument in the distributions of measured fluorescence lifetimes, we incorporate the distribution for the instrument response function (IRF) into our model. This distribution can be well fitted by a Gaussian or as we explained in the Supplementary materials Sec. S2, it can easily be generalized to any functional form such as Gamma like what it has been used in the previous publications.^60^

As a dynamical model for the state trajectories, we consider a Markovian property and accordingly,^45–58^ de-fine a rate matrix 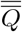. Since, in our model we consider the photon arrival times as the timestamp, for each time that we detect a photon, macro time t at time *T*_*t*_, we can calculate the transition probability matrix

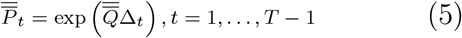

where, Δ_*t*_ is the inter-arrival time between detected photons *t* and *t* + 1. As the result of the Eq. (5), the adaptation of states through time are

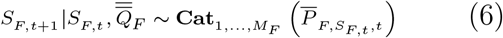

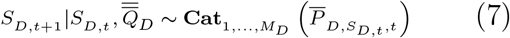

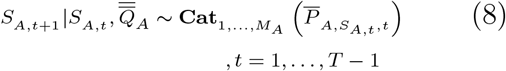

where, *S*_*F,t*_, *S*_*D,t*_, *S*_*A,t*_ are the FRET state, donor-PIFE state, and acceptor-PIFE state at the time of detected photon *t*. The 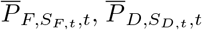, and 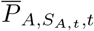 are equal to the *S*_*F,t*_, *S*_*D,t*_ and *S*_*A,t*_ rows of the corresponding transition probability matrices in transition between detected photon *t* and *t* + 1. Additional detail on calculating the transition matrices and updating the state trajectories can be found in the Supplementary Materials Sec. S4.

### Model inference

In this study we follow the Bayesian statistics^37,38,77^ to estimate the variables of interest (*η*_*A*_, *η*_*D*_, *μ*_back,*A*_, *μ*_back,*D*_, 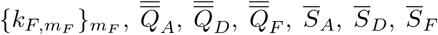. Since we are using the Bayesian approach, we need to define the priors and our choices for these priors are described in detail in the Supplementary Materials Sec. S3.

Once the choices for the priors of above variables are made, we form a joint posterior probability ℙ(*η*_*A*_, *η*_*D*_, *μ*_back,*A*_, *μ*_back,*D*_, 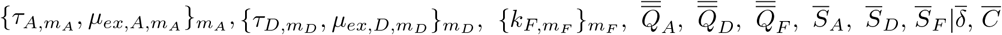 encompassing all unknown variables which we may wish to determine.

Due to the complexity of our posterior with respect to our variables, we cannot construct an analytic form for our posterior. For this reason, we develop a computational scheme exploiting Markov chain Monte Carlo^77,78^ that can be used to generate pseudo-random samples from this posterior. Our MCMC exploits a Gibbs sampling scheme^77–79^. Accordingly, posterior samples are generated by updating each one of the variables involved sequentially by sampling conditioned on all other variables and measurements 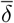 and 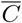. Conceptually, the steps involved in the generation of each posterior sample are:

- Sampling the FRET state trajectory through time 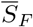
- Sampling the donor-PIFE state trajectory through time 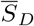
- Sampling the acceptor-PIFE state trajectory through time 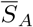
- Sampling the ET rates 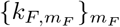
- Sampling the donor lifetimes 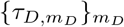
- Sampling the acceptor lifetimes 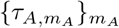
- Sampling the FRET state transition rate matrix 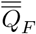
- Sampling the donor-PIFE state transition rate matrix 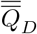
- Sampling the acceptor-PIFE state transition rate matrix 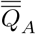
- Sampling the donor and acceptor excitation rates

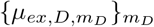 and 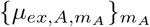

- Sampling the donor and acceptor background photon rates *μ* _back,*D*_ and *μ* _back,*A*_
- Sampling the cross-talk ratios for donor and acceptor channels, *η*_*D*_ and *η*_*A*_

## DATA AVAILABILITY

The data generated and analyzed in this study are freely available from the corresponding author upon request.

## ACKNOWLEDGMENTS

SJ acknowledges W. Taylor Cottle and Alberto Marin-Gonzalez for their help in writing this manuscript.

## AUTHOR CONTRIBUTIONS

SJ developed analysis software, computational tools and analyzed data; SJ, TH conceived research; TH oversaw all aspects of the projects.

## CODE AVAILABILITY

The source code and the installable app (written in Matlab) versions of the methods developed herewith are freely available to download at [https://github.com/sinajazani/FL_smFRET_smPIFE].

## Refrences

1 B. W. Van Der Meer, G. Coker, and S.-Y. S. Chen, Resonance energy transfer: theory and data VCH publishers, 1994).

2 P. Wu and L. Brand, “Resonance energy transfer: methods and applications,” Analytical biochemistry 218, 1–13 (1994).

3 R. M. Clegg, “[18] fluorescence resonance energy transfer and nucleic acids,” Methods in enzymology 211, 353–388 (1992).

4 W. E. Moerner and M. Orrit, “Illuminating single molecules in condensed matter,” Science 283, 1670–1676 (1999).

5 S. Nie and R. N. Zare, “Optical detection of single molecules,” Annual review of biophysics and biomolecular structure 26, 567– 596 (1997).

6 X. S. Xie and J. K. Trautman, “Optical studies of single molecules at room temperature,” Annual review of physical chemistry 49, 441–480 (1998).

7 R. M. Clegg, “Fluorescence resonance energy transfer,” Current opinion in biotechnology 6, 103–110 (1995).

8 L. Stryer and R. P. Haugland, “Energy transfer: a spectroscopic ruler.” Proceedings of the National Academy of Sciences of the United States of America 58, 719 (1967).

9 S. Weiss, “Measuring conformational dynamics of biomolecules by single molecule fluorescence spectroscopy,” Nature structural biology 7, 724–729 (2000).

10 X. Zhuang, “Single-molecule rna science,” Annu. Rev. Biophys. Biomol. Struct. 34, 399–414 (2005).

11 T. Paul, S. C. Bera, and P. P. Mishra, “Direct observation of breathing dynamics at the mismatch induced dna bubble with nanometre accuracy: a smfret study,” Nanoscale 9, 5835–5842 (2017).

12 T. Ha, T. Enderle, D. Ogletree, D. S. Chemla, P. R. Selvin, and S. Weiss, “Probing the interaction between two single molecules: fluorescence resonance energy transfer between a single donor and a single acceptor,” Proceedings of the National Academy of Sciences 93, 6264–6268 (1996).

13 G. Schütz, W. Trabesinger, and T. Schmidt, “Direct observation of ligand colocalization on individual receptor molecules,” Biophysical journal 74, 2223–2226 (1998).

14 J. C. Sanders and E. D. Holmstrom, “Integrating single-molecule fret and biomolecular simulations to study diverse interactions between nucleic acids and proteins,” Essays in biochemistry 65, 37 (2021).

15 E. Šimková and D. Staněk, “Probing nucleic acid interactions and pre-mrna splicing by förster resonance energy transfer (fret) microscopy,” International journal of molecular sciences 13, 14929– 14945 (2012).

16 R. Lamichhane, A. Solem, W. Black, and D. Rueda, “Singlemolecule fret of protein–nucleic acid and protein–protein complexes: surface passivation and immobilization,” Methods 52, 192–200 (2010).

17 K. Truong and M. Ikura, “The use of fret imaging microscopy to detect protein–protein interactions and protein conformational changes in vivo,” Current opinion in structural biology 11, 573– 578 (2001).

18 R. Roy, S. Hohng, and T. Ha, “A practical guide to singlemolecule fret,” Nature methods 5, 507–516 (2008).

19 A. Jain, R. Liu, B. Ramani, E. Arauz, Y. Ishitsuka, K. Ragunathan, J. Park, J. Chen, Y. K. Xiang, and T. Ha, “Probing cellular protein complexes using single-molecule pull-down,” Nature 473, 484–488 (2011).

20 B. Schuler and W. A. Eaton, “Protein folding studied by singlemolecule fret,” Current opinion in structural biology 18, 16–26 (2008).

21 H. Mazal and G. Haran, “Single-molecule fret methods to study the dynamics of proteins at work,” Current opinion in biomedical engineering 12, 8–17 (2019).

22 B. Hellenkamp, S. Schmid, O. Doroshenko, O. Opanasyuk, R. Kühnemuth, S. Rezaei Adariani, B. Ambrose, M. Aznauryan, A. Barth, V. Birkedal, et al., “Precision and accuracy of single-molecule fret measurements—a multi-laboratory benchmark study,” Nature methods 15, 669–676 (2018).

23 M. Blanco and N. G. Walter, “Analysis of complex single-molecule fret time trajectories,” in Methods in enzymology, Vol. 472 (Elsevier, 2010) pp. 153–178.

24 D. Singh, Y. Wang, J. Mallon, O. Yang, J. Fei, A. Poddar, D. Ceylan, S. Bailey, and T. Ha, “Mechanisms of improved specificity of engineered cas9s revealed by single-molecule fret analysis,” Nature structural & molecular biology 25, 347–354 (2018).

25 H. Hwang, H. Kim, and S. Myong, “Protein induced fluorescence enhancement as a single molecule assay with short distance sensitivity,” Proceedings of the National Academy of Sciences 108, 7414–7418 (2011).

26 S. Myong, S. Cui, P. V. Cornish, A. Kirchhofer, M. U. Gack, J. U. Jung, K.-P. Hopfner, and T. Ha, “Cytosolic viral sensor rig-i is a 5’-triphosphate–dependent translocase on double-stranded rna,” Science 323, 1070–1074 (2009).

27 M. E. Sanborn, B. K. Connolly, K. Gurunathan, and M. Levitus, “Fluorescence properties and photophysics of the sulfoindocyanine cy3 linked covalently to dna,” The Journal of Physical Chemistry B 111, 11064–11074 (2007).

28 M. Levitus and S. Ranjit, “Cyanine dyes in biophysical research: the photophysics of polymethine fluorescent dyes in biomolecular environments,” Quarterly reviews of biophysics 44, 123–151 (2011).

29 K. Jia, Y. Wan, A. Xia, S. Li, F. Gong, and G. Yang, “Characterization of photoinduced isomerization and intersystem crossing of the cyanine dye cy3,” The Journal of Physical Chemistry A 111, 1593–1597 (2007).

30 H. Hwang and S. Myong, “Protein induced fluorescence enhancement (pife) for probing protein–nucleic acid interactions,” Chemical Society Reviews 43, 1221–1229 (2014).

31 M. Sorokina, H.-R. Koh, S. S. Patel, and T. Ha, “Fluorescent lifetime trajectories of a single fluorophore reveal reaction intermediates during transcription initiation,” Journal of the American Chemical Society 131, 9630–9631 (2009).

32 E. Lerner, E. Ploetz, J. Hohlbein, T. Cordes, and S. Weiss, “A quantitative theoretical framework for protein-induced fluorescence enhancement–förster-type resonance energy transfer (pife-fret),” The Journal of Physical Chemistry B 120, 6401–6410 (2016).

33 F. Rashid, V.-S. Raducanu, M. S. Zaher, M. Tehseen, S. Habuchi, and S. M. Hamdan, “Initial state of dna-dye complex sets the stage for protein induced fluorescence modulation,” Nature communications 10, 1–14 (2019).

34 B. Nguyen, M. A. Ciuba, A. G. Kozlov, M. Levitus, and T. M. Lohman, “Protein environment and dna orientation affect protein-induced cy3 fluorescence enhancement,” Biophysical journal 117, 66–73 (2019).

35 M. Tavakoli, S. Jazani, I. Sgouralis, W. Heo, K. Ishii, T. Tahara, and S. Pressé, “Direct photon-by-photon analysis of time-resolved pulsed excitation data using bayesian nonparametrics,” Cell Reports Physical Science 1, 100234 (2020).

36 M. Fazel, S. Jazani, L. Scipioni, A. Vallmitjana, E. Gratton, M. A. Digman, and S. Pressé, “High resolution fluorescence lifetime maps from minimal photon counts,” ACS Photonics 9, 1015–1025 (2022).

37 A. Lee, K. Tsekouras, C. Calderon, C. Bustamante, and S. Pressé, “Unraveling the thousand word picture: an introduction to super-resolution data analysis,” Chemical reviews 117, 7276–7330 (2017).

38 M. Tavakoli, J. N. Taylor, C.-B. Li, T. Komatsuzaki, and S. Pressé, “Single molecule data analysis: An introduction,” arXiv preprint arXiv:1606.00403 (2016).

39 K. E. Hines, “A primer on bayesian inference for biophysical systems,” Biophysical journal 108, 2103–2113 (2015).

40 M. Fazel, S. Jazani, L. Scipioni, A. Vallmitjana, E. Gratton, M. A. Digman, and S. Pressé, “High resolution fluorescence life-time maps from minimal photon counts,” ACS Nano (2022).

41 N. K. Lee, A. N. Kapanidis, Y. Wang, X. Michalet, J. Mukhopadhyay, R. H. Ebright, and S. Weiss, “Accurate fret measurements within single diffusing biomolecules using alternating-laser excitation,” Biophysical journal 88, 2939–2953 (2005).

42 S. Ranjit, L. Malacrida, D. M. Jameson, and E. Gratton, “Fit-free analysis of fluorescence lifetime imaging data using the phasor approach,” Nature protocols 13, 1979–2004 (2018).

43 M. A. Digman, V. R. Caiolfa, M. Zamai, and E. Gratton, “The phasor approach to fluorescence lifetime imaging analysis,” Biophysical journal 94, L14–L16 (2008).

44 Y. He, M. Lu, and H. P. Lu, “Single-molecule photon stamping fret spectroscopy study of enzymatic conformational dynamics,” Physical Chemistry Chemical Physics 15, 770–775 (2013).

45 S. A. McKinney, C. Joo, and T. Ha, “Analysis of single-molecule fret trajectories using hidden markov modeling,” Biophysical journal 91, 1941–1951 (2006).

46 J. E. Bronson, J. Fei, J. M. Hofman, R. L. Gonzalez Jr, and C. H. Wiggins, “Learning rates and states from biophysical time series: a bayesian approach to model selection and single-molecule fret data,” Biophysical journal 97, 3196–3205 (2009).

47 D. Kelly, M. Dillingham, A. Hudson, and K. Wiesner, “A new method for inferring hidden markov models from noisy time sequences,” PloS one 7, e29703 (2012).

48 B. G. Keller, A. Kobitski, A. Jäschke, G. U. Nienhaus, and F. Noé, “Complex rna folding kinetics revealed by single-molecule fret and hidden markov models,” Journal of the American Chemical Society 136, 4534–4543 (2014).

49 K. Okamoto and Y. Sako, “Variational bayes analysis of a photon-based hidden markov model for single-molecule fret trajectories,” Biophysical journal 103, 1315–1324 (2012).

50 J. F. Beausang, C. Zurla, C. Manzo, D. Dunlap, L. Finzi, and P. C. Nelson, “Dna looping kinetics analyzed using diffusive hidden markov model,” Biophysical journal 92, L64–L66 (2007).

51 M. Andrec, R. M. Levy, and D. S. Talaga, “Direct determination of kinetic rates from single-molecule photon arrival trajectories using hidden markov models,” The Journal of Physical Chemistry A 107, 7454–7464 (2003).

52 S. Uphoff, K. Gryte, G. Evans, and A. N. Kapanidis, “Improved temporal resolution and linked hidden markov modeling for switchable single-molecule fret,” ChemPhysChem 12, 571– 579 (2011).

53 F. Noé, H. Wu, J.-H. Prinz, and N. Plattner, “Projected and hidden markov models for calculating kinetics and metastable states of complex molecules,” The Journal of chemical physics 139, 11B609 1 (2013).

54 M. Pirchi, R. Tsukanov, R. Khamis, T. E. Tomov, Y. Berger, D. C. Khara, H. Volkov, G. Haran, and E. Nir, “Photon-by-photon hidden markov model analysis for microsecond single-molecule fret kinetics,” The Journal of Physical Chemistry B 120, 13065–13075 (2016).

55 F. E. Müllner, S. Syed, P. R. Selvin, and F. J. Sigworth, “Improved hidden markov models for molecular motors, part 1: basic theory,” Biophysical journal 99, 3684–3695 (2010).

56 J.-W. Meent, J. Bronson, F. Wood, R. Gonzalez Jr, and C. Wiggins, “Hierarchically-coupled hidden markov models for learning kinetic rates from single-molecule data,” in International Conference on Machine Learning (PMLR, 2013) pp. 361–369.

57 J. Stigler and M. Rief, “Hidden markov analysis of trajectories in single-molecule experiments and the effects of missed events,” ChemPhysChem 13, 1079–1086 (2012).

58 D. S. Talaga, “Markov processes in single molecule fluorescence,” Current opinion in colloid & interface science 12, 285–296 (2007).

59 H. Akaike, “Information theory and an extension of the maximum likelihood principle,” in Selected papers of hirotugu akaike (Springer, 1998) pp. 199–213.

60 H. S. Chung, J. M. Louis, and I. V. Gopich, “Analysis of fluorescence lifetime and energy transfer efficiency in single-molecule photon trajectories of fast-folding proteins,” The Journal of Physical Chemistry B 120, 680–699 (2016).

61 J. Park, S. Myong, A. Niedziela-Majka, K. S. Lee, J. Yu, T. M. Lohman, and T. Ha, “Pcra helicase dismantles reca filaments by reeling in dna in uniform steps,” Cell 142, 544–555 (2010).

62 R. Dixit and R. Cyr, “Cell damage and reactive oxygen species production induced by fluorescence microscopy: effect on mitosis and guidelines for non-invasive fluorescence microscopy,” The Plant Journal 36, 280–290 (2003).

63 V. Magidson and A. Khodjakov, “Circumventing photodamage in live-cell microscopy,” Methods in Cell Biology 114, 545–560 (2013).

64 J. J. McCann, U. B. Choi, L. Zheng, K. Weninger, and M. E. Bowen, “Optimizing methods to recover absolute fret efficiency from immobilized single molecules,” Biophysical Journal 99, 961– 970 (2010).

65 Y. Gidi, M. Götte, and G. Cosa, “Conformational changes spanning angstroms to nanometers via a combined protein-induced fluorescence enhancement–förster resonance energy transfer method,” The Journal of Physical Chemistry B 121, 2039–2048 (2017).

66 S. Baker and R. D. Cousins, “Clarification of the use of chi-square and likelihood functions in fits to histograms,” Nuclear Instruments and Methods in Physics Research 221, 437–442 (1984).

67 G. Cowan, Statistical data analysis (Oxford university press, 1998).

68 I. Sgouralis, M. Whitmore, L. Lapidus, M. J. Comstock, and S. Pressé, “Single molecule force spectroscopy at high data acquisition: A bayesian nonparametric analysis,” The Journal of chemical physics 148, 123320 (2018).

69 I. Sgouralis and S. Pressé, “Icon: an adaptation of infinite hmms for time traces with drift,” Biophysical journal 112, 2117–2126 (2017).

70 E. Snelson and Z. Ghahramani, “Local and global sparse gaussian process approximations,” in Artificial Intelligence and Statistics (PMLR, 2007) pp. 524–531.

71 J. S. Bryan IV, I. Sgouralis, and S. Pressé, “Inferring effective forces for langevin dynamics using gaussian processes,” The Journal of Chemical Physics 152, 124106 (2020).

72 Z. Kilic, I. Sgouralis, and S. Pressé, “Generalizing hmms to continuous time for fast kinetics: Hidden markov jump processes,” Biophysical Journal 120, 409–423 (2021).

73 L. V. Jospin, W. Buntine, F. Boussaid, H. Laga, and M. Bennamoun, “Hands-on bayesian neural networks–a tutorial for deep learning users,” arXiv preprint arXiv:2007.06823 (2020).

74 J. Lampinen and A. Vehtari, “Bayesian approach for neural networks—review and case studies,” Neural Networks 14, 257–274 (2001).

75 D. Titterington, “Bayesian methods for neural networks and related models,” Statistical Science, 128–139 (2004).

76 H. Wang and D.-Y. Yeung, “A survey on bayesian deep learning,” ACM Computing Surveys (CSUR) 53, 1–37 (2020).

77 A. Gelman, J. Hwang, and A. Vehtari, “Understanding predictive information criteria for bayesian models,” Statistics and computing 24, 997–1016 (2014).

78 C. P. Robert, G. Casella, and G. Casella, Introducing monte carlo methods with r, Vol. 18 (Springer, 2010).

79 U. Von Toussaint, “Bayesian inference in physics,” Reviews of Modern Physics 83, 943 (2011).

