## Supplementary materials for "Fluorescence lifetime analysis of smFRET with contribution of PIFE on donor and acceptor"

Sina Jazani
*Department of Biophysics and Biophysical Chemistry,*
*Johns Hopkins University School of Medicine, Baltimore, MD, USA*

Taekjip Ha
*Department of Biophysics and Biophysical Chemistry,*
*Johns Hopkins University School of Medicine, Baltimore, MD, USA*
*Department of Biophysics, Johns Hopkins University, Baltimore, MD, USA*
*Department of Biomedical Engineering, Johns Hopkins University, Baltimore, MD, USA and*
*Howard Hughes Medical Institute, Baltimore, MD, USA*

Here we talk through supplementary materials and details of our method. We arrange the supplementary materials to different sections to discuss: (i) Complementary results. (ii) Methodology and Mathematical details of our method, including the simulation approach, choices of priors, and the inference method. (iv) Summary of notation and other conventions used throughout this study as well as detailed parameter choices for the simulations and analyses.

### Contents

S1. Additional results 3

S2. Summary of our results in the form of point estimates 5

S3. Detailed methods description 7
S3.1. Description of the data simulation 7
S3.1.1. Description of the state trajectories 7
S3.1.2. Description of the weights on the detected photons 7
S3.1.3. Description of the probability on detection channel 8
S3.1.4. Description of the probability on photon arrival times 9

S4. Detailed description of the inference framework 12
S4.1. Description of prior probability distributions 12
S4.1.1. Prior on acceptor and donor channels background photon emission rates $\mu_{back,A}$ and $\mu_{back,D}$ 13
S4.1.2. Priors on the acceptor and donor channel cross-talk ratios $\eta_A$, $\eta_D$ 14
S4.1.3. Prior on acceptor and donor excitation rates $\{\mu_{ex,A,m_A}\}_{m_A}$ and $\{\mu_{ex,D,m_D}\}_{m_D}$ 14
S4.1.4. Priors on the acceptor and donor lifetimes $\{\tau_{A,m_A}\}_{m_A}$ and $\{\tau_{D,m_D}\}_{m_D}$ and ET rates $\{k_{F,m_F}\}_{m_F}$ 14
S4.1.5. Prior on the initial states acceptor $S_{A,1}$, donor $S_{D,1}$, and FRET $S_{F,1}$ 14
S4.1.6. Prior on the wights of initial states for acceptor-PIFE $\overline{\zeta}_A$, donor-PIFE $\overline{\zeta}_D$, and FRET $\overline{\zeta}_F$ 14
S4.1.7. Prior on the elements of state transition rate matrices $\overline{\overline{Q}}_A$, $\overline{\overline{Q}}_D$, $\overline{\overline{Q}}_F$ 15
S4.2. Summary of model equations 16

S5. Description of the computational scheme 19
S5.1. Overview of the sampling updates 20
S5.2. Update the state trajectory of acceptor lifetimes $\overline{S}_A$ 21
S5.3. Update the state trajectory of donor lifetimes $\overline{S}_D$ 21
S5.4. Update the state trajectory of energy transfer rates $\overline{S}_F$ 22
S5.5. Update the acceptor-PIFE state transition matrix $\overline{\overline{Q}}_A$ 23
S5.6. Update the transition matrix of donor lifetime trajectory $\overline{\overline{Q}}_D$ 23
S5.7. Update the transition matrix of FRET trajectory $\overline{\overline{Q}}_F$ 24
S5.8. Update the weight on the initial state of acceptor PIFE $\overline{\zeta}_A$ 25
S5.9. Update the weight on initial state of donor PIFE $\overline{\zeta}_D$ 25
S5.10. Update the weight on initial state of FRET $\overline{\zeta}_F$ 25
S5.11. Jointly update the acceptor lifetimes of all states $\{\tau_{A,m_A}\}_{m_A}$, donor lifetimes of all states $\{\tau_{D,m_D}\}_{m_D}$, and resonance energy transfer rates of all states $\{k_{F,m_F}\}_{m_F}$ 25
S5.12. Jointly update the cross-talk ratios of acceptor and donor channels, $\eta_A$ and $\eta_D$ 28
S5.13. Jointly update the acceptor excitation rates $\{\mu_{ex,A,m_A}\}_{m_A}$, donor excitation rates $\{\mu_{ex,D,m_D}\}_{m_D}$, acceptor channel background photon emission rate $\mu_{\text{back},A}$, and donor channel background photon emission rate $\mu_{\text{back},D}$ 29
S5.14. Maximum of posteriori (MAP) estimate 31

S6. Summary of notation, abbreviations, parameters and other options 32

### S1. Additional results

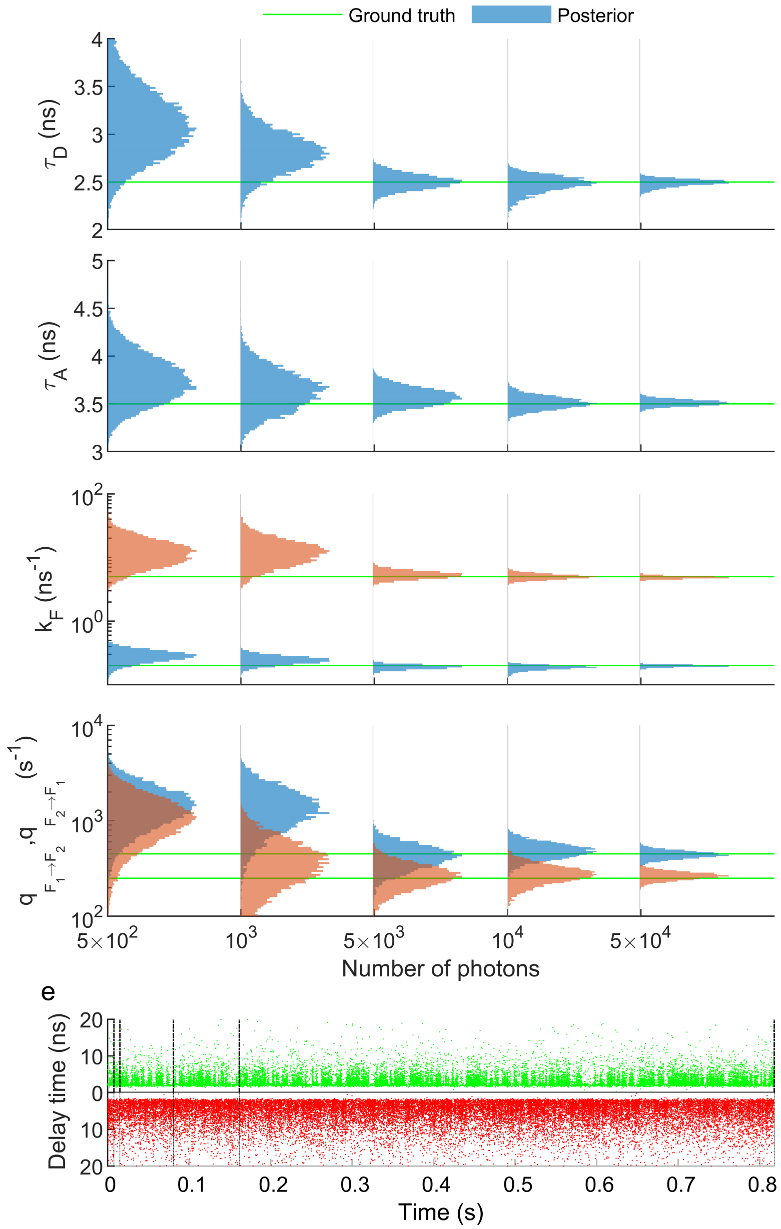


FIG. S1. **Analysis of synthetic time trace of a two FRET states system with different numbers of photons. (a-d)** Posterior probability distributions of the donor lifetime, acceptor lifetime, energy transfer rate and the FRET state transition rates with respect to different numbers of photons. Here, we are using $5\times10^2$, $10^3$, $5\times10^3$, $10^4$ and $5\times10^4$ photons as the input signals. The ground truths are shown with solid green color lines. **(e)** The single-photon arrival time traces are collected with donor and acceptor channels. In order to be as close as possible to the experimental data sets, each time trace contains multiple individual collected data sets visualized as one time trace with approximately $5\times10^4$ photons in total.

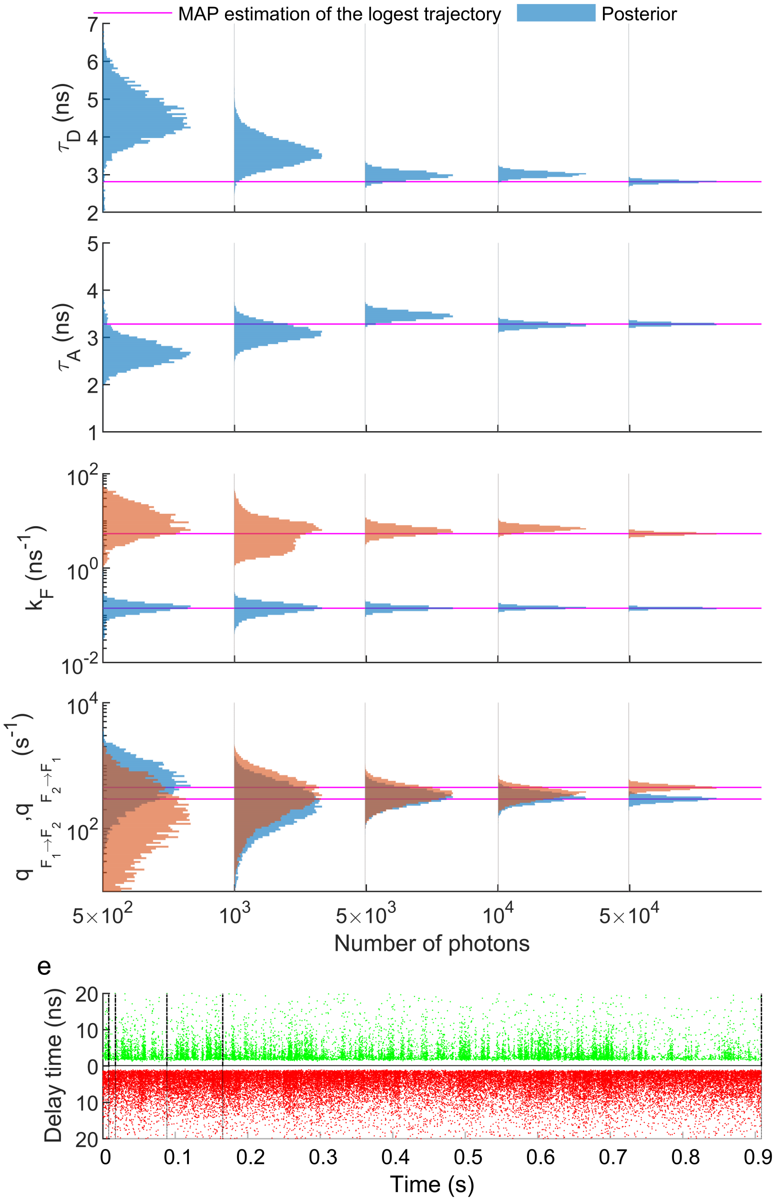


FIG. S2. **Analysis of $\alpha$3D experimental data with different numbers of photons. (a-d)** Posterior probability distributions of the donor lifetime, acceptor lifetime, energy transfer rate and the FRET state transition rates with respect to different numbers of photons. Here, we are using $5{\times}10^2$, $10^3$, $5 \times 10^3$, $10^4$ and $5 \times 10^4$ photons as the input signals. The most probable values are shown with solid magenta color lines. **(e)** Experimental single-photon arrival time traces of $\alpha$3D collected with donor and acceptor channels. Each time trace contains multiple individual collected data sets visualized as one time trace with approximately $5{\times}10^4$ photons in total.

### S2. Summary of our results in the form of point estimates

TABLE S1. Listed characteristic values (point estimates) of the posterior probability distributions of resonance energy transfer rates $\{k_{F,m_F}\}_{m_F}$, donor lifetimes $\{\tau_{D,m_D}\}_{m_D}$, acceptor lifetime $\{\tau_{A,m_A}\}_{m_A}$, donor channel cross-talk $\eta_D$, and acceptor channel cross-talk $\eta_A$. Mean and std refer to mean value and standard deviation of the posterior (i.e. square root of variance). Values are listed according to the figures.

| | $\{k_{F,m_F}\}_{m_F}$ ($s^{-1}$) | | $\{\tau_{D,m_D}\}_{m_D}$ (ns) | | $\{\tau_{A,m_A}\}_{m_A}$ (ns) | | $\eta_D$ | | $\eta_A$ | |
|---|---|---|---|---|---|---|---|---|---|---|
| | mean | std | mean | std | mean | std | mean | std | mean | std |
| Fig. 3(a1) | 0.097<br>0.919 | 0.004<br>0.060 | 3.930 | 0.063 | 3.634 | 0.076 | 0.073 | 0.016 | 0.096 | 0.012 |
| Fig. 3(a2) | 0.093<br>0.890 | 0.004<br>0.079 | 3.908 | 0.062 | 3.604 | 0.083 | 0.066 | 0.027 | 0.094 | 0.020 |
| Fig. 3(a3) | 0.094<br>0.776 | 005<br>0.097 | 3.904 | 0.070 | 3.663 | 0.099 | 0.045 | 0.053 | 0.091 | 0.025 |
| Fig. 4(a1) | 0.482<br>0.096 | 0.026<br>0.009 | 1.921<br>3.768 | 0.044<br>0.148 | 2.088<br>4.170 | 0.062<br>0.159 | 0.081 | 0.018 | 0.116 | 0.024 |
| Fig. 4(a2) | 0.480<br>0.097<br>2.866 | 0.015<br>0.005<br>0.189 | 3.840<br>0.965<br>1.813 | 0.082<br>0.036<br>0.089 | 4.201<br>1.060<br>2.097 | 0.097<br>0.025<br>0.125 | 0.086 | 0.014 | 0.098 | 0.005 |
| Fig. 4(a3) | 0.477<br>2.481<br>0.926<br>0.096 | 0.014<br>0.189<br>0.045<br>0.005 | 2.902<br>0.909<br>1.961<br>3.895 | 0.099<br>0.032<br>0.042<br>0.089 | 2.148<br>1.039<br>3.245<br>4.252 | 0.058<br>0.027<br>0.072<br>0.079 | 0.087 | 0.007 | 0.096 | 0.007 |
| Fig. 5(a1) | 0.135<br>4.284 | 0.012<br>0.777 | 2.704 | 0.089 | 3.173 | 0.069 | 0.24 | 0.042 | 0.051 | 0.010 |
| Fig. 5(a2) | 0.092<br>1.470 | 0.014<br>0.250 | 2.540 | 0.083 | 3.333 | 0.086 | 0.28 | 0.021 | 0.090 | 0.022 |
| Fig. 5(a3) | 0.057<br>1.420 | 0.055<br>0.385 | 1.705 | 0.088 | 3.622 | 0.083 | 0.311 | 0.012 | 0.160 | 0.010 |
| Fig. S1($5\times10^2$) | 12.81<br>0.281 | 6.393<br>0.061 | 3.112 | 0.362 | 3.754 | 0.245 | 0.178 | 0.071 | 0.070 | 0.020 |
| Fig. S1($\times10^3$) | 12.80<br>0.244 | 5.990<br>0.038 | 2.808 | 0.209 | 3.634 | 0.207 | 0.133 | 0.065 | 0.074 | 0.019 |
| Fig. S1($5\times10^3$) | 5.36<br>0.190 | 0.721<br>0.015 | 2.500 | 0.078 | 3.572 | 0.093 | 0.254 | 0.022 | 0.042 | 0.007 |
| Fig. S1($\times10^4$) | 4.997<br>0.186 | 0.488<br>0.015 | 2.474 | 0.088 | 3.510 | 0.066 | 0.262 | 0.013 | 0.049 | 0.004 |
| Fig. S1($5\times10^4$) | 4.876<br>0.196 | 0.219<br>0.007 | 2.489 | 0.039 | 3.51 | 0.032 | 0.238 | 0.019 | 0.051 | 0.003 |
| Fig. S2($5\times10^2$) | 11.64<br>0.203 | 8.313<br>0.436 | 4.549 | 0.682 | 2.689 | 0.271 | 0.176 | 0.111 | 0.070 | 0.039 |
| Fig. S2($\times10^3$) | 6.296<br>0.141 | 4.562<br>0.037 | 3.606 | 0.359 | 3.095 | 0.182 | 0.214 | 0.125 | 0.045 | 0.019 |
| Fig. S2($5\times10^3$) | 6.273<br>0.142 | 1.344<br>0.017 | 2.982 | 0.199 | 3.459 | 0.084 | 0.345 | 0.032 | 0.051 | 0.006 |
| Fig. S2($\times10^4$) | 6.966<br>0.146 | 0.943<br>0.011 | 3.008 | 0.073 | 3.259 | 0.053 | 0.309 | 0.023 | 0.046 | 0.004 |
| Fig. S2($5\times10^4$) | 5.37<br>0.142 | 0.376<br>0.006 | 2.814 | 0.037 | 3.282 | 0.029 | 0.328 | 0.007 | 0.047 | 0.002 |

TABLE S2. Listed characteristic values (point estimates) of the posterior probability distributions of FRET transition rate matrix $Q_F$, donor PIFE transition rate matrix $Q_D$, and acceptor PIFE transition rate matrix $Q_A$. Mean and std refer to mean value and standard deviation of the posterior (i.e. square root of variance). Values are listed according to the figures. Since the diagonal values of the transition matrix are equal to the negative sum of the row values, we are not show them.

| | $Q_F$ (s$^{-1}$) | | $Q_D$ (s$^{-1}$) | | $Q_A$ (s$^{-1}$) | |
|---|---|---|---|---|---|---|
| | mean | std | mean | std | mean | std |
| Fig. 3(a1) | $\begin{bmatrix} - & 49.82 \\ 12.90 & - \end{bmatrix}$ | $\begin{bmatrix} - & 28.56 \\ 8.159 & - \end{bmatrix}$ | - | - | - | - |
| Fig. 3(a2) | $\begin{bmatrix} - & 94.19 \\ 531.8 & - \end{bmatrix}$ | $\begin{bmatrix} - & 21.35 \\ 129.3 & - \end{bmatrix}$ | - | - | - | - |
| Fig. 3(a3) | $\begin{bmatrix} - & 5230 \\ 990.1 & - \end{bmatrix}$ | $\begin{bmatrix} - & 717.3 \\ 155.7 & - \end{bmatrix}$ | - | - | - | - |
| Fig. 4(a1) | $\begin{bmatrix} - & 76.27 \\ 44.96 & - \end{bmatrix}$ | $\begin{bmatrix} - & 27.19 \\ 19.25 & - \end{bmatrix}$ | $\begin{bmatrix} - & 60.09 \\ 19.19 & - \end{bmatrix}$ | $\begin{bmatrix} - & 36.03 \\ 10.48 & - \end{bmatrix}$ | $\begin{bmatrix} - & 44.55 \\ 14.22 & - \end{bmatrix}$ | $\begin{bmatrix} - & 30.82 \\ 9.96 & - \end{bmatrix}$ |
| Fig. 4(a2) | $\begin{bmatrix} - & 73.93 & 40.10 \\ 30.85 & - & 69.68 \\ 61.13 & 27.08 & - \end{bmatrix}$ | $\begin{bmatrix} - & 29.04 & 22.20 \\ 18.49 & - & 27.90 \\ 23.56 & 19.02 & - \end{bmatrix}$ | $\begin{bmatrix} - & 53.68 & 10.56 \\ 4.570 & - & 125.44 \\ 43.04 & 25.47 & - \end{bmatrix}$ | $\begin{bmatrix} - & 23.98 & 19.22 \\ 7.840 & - & 68.59 \\ 18.32 & 29.90 & - \end{bmatrix}$ | $\begin{bmatrix} - & 44.31 & 89.38 \\ 57.92 & - & 59.67 \\ 28.83 & 8.490 & - \end{bmatrix}$ | $\begin{bmatrix} - & 18.02 & 72.07 \\ 27.67 & - & 60.27 \\ 20.38 & 10.52 & - \end{bmatrix}$ |
| Fig. 4(a3) | $\begin{bmatrix} - & 16.86 & 57.94 & 111.8 \\ 24.64 & - & 26.13 & 6.584 \\ 38.91 & 10.36 & - & 37.52 \\ 43.47 & 97.61 & 68.81 & - \end{bmatrix}$ | $\begin{bmatrix} - & 21.31 & 44.52 & 40.19 \\ 14.93 & - & 29.89 & 13.13 \\ 20.26 & 14.31 & - & 22.38 \\ 23.31 & 49.38 & 44.14 & - \end{bmatrix}$ | $\begin{bmatrix} - & 53.11 & 11.60 & 28.28 \\ 19.19 & - & 46.82 & 13.00 \\ 65.82 & 23.55 & - & 64.46 \\ 36.77 & 65.48 & 36.46 & - \end{bmatrix}$ | $\begin{bmatrix} - & 40.61 & 18.27 & 24.56 \\ 20.89 & - & 25.84 & 13.51 \\ 54.40 & 29.47 & - & 44.67 \\ 36.88 & 54.52 & 26.61 & - \end{bmatrix}$ | $\begin{bmatrix} - & 14.88 & 37.58 & 71.51 \\ 79.25 & - & 57.09 & 21.73 \\ 37.17 & 31.66 & - & 33.37 \\ 13.60 & 58.92 & 21.81 & - \end{bmatrix}$ | $\begin{bmatrix} - & 16.73 & 46.69 & 32.15 \\ 46.38 & - & 37.93 & 28.06 \\ 39.45 & 27.32 & - & 25.83 \\ 13.96 & 29.78 & 24.48 & - \end{bmatrix}$ |
| Fig. 5(a1) | $\begin{bmatrix} - & 320.4 \\ 390.8 & - \end{bmatrix}$ | $\begin{bmatrix} - & 76.36 \\ 96.11 & - \end{bmatrix}$ | - | - | - | - |
| Fig. 5(a2) | $\begin{bmatrix} - & 1685 \\ 1353 & - \end{bmatrix}$ | $\begin{bmatrix} - & 335.3 \\ 282.6 & - \end{bmatrix}$ | - | - | - | - |
| Fig. 5(a3) | $\begin{bmatrix} - & 2290 \\ 5877 & - \end{bmatrix}$ | $\begin{bmatrix} - & 889.1 \\ 1719 & - \end{bmatrix}$ | - | - | - | - |
| Fig. S1(5×$10^2$) | $\begin{bmatrix} - & 1489 \\ 1101 & - \end{bmatrix}$ | $\begin{bmatrix} - & 745.1 \\ 613.2 & - \end{bmatrix}$ | - | - | - | - |
| Fig. S1(×$10^3$) | $\begin{bmatrix} - & 1392 \\ 384.6 & - \end{bmatrix}$ | $\begin{bmatrix} - & 697.5 \\ 222.7 & - \end{bmatrix}$ | - | - | - | - |
| Fig. S1(5×$10^3$) | $\begin{bmatrix} - & 427.6 \\ 268.1 & - \end{bmatrix}$ | $\begin{bmatrix} - & 123.7 \\ 78.51 & - \end{bmatrix}$ | - | - | - | - |
| Fig. S1(×$10^4$) | $\begin{bmatrix} - & 486.3 \\ 276.9 & - \end{bmatrix}$ | $\begin{bmatrix} - & 92.65 \\ 61.61 & - \end{bmatrix}$ | - | - | - | - |
| Fig. S1(5×$10^4$) | $\begin{bmatrix} - & 440.9 \\ 265.5 & - \end{bmatrix}$ | $\begin{bmatrix} - & 41.16 \\ 25.72 & - \end{bmatrix}$ | - | - | - | - |
| Fig. S2(5×$10^2$) | $\begin{bmatrix} - & 648.5 \\ 225.3 & - \end{bmatrix}$ | $\begin{bmatrix} - & 462.9 \\ 267.3 & - \end{bmatrix}$ | - | - | - | - |
| Fig. S2(×$10^3$) | $\begin{bmatrix} - & 274.9 \\ 375.7 & - \end{bmatrix}$ | $\begin{bmatrix} - & 198.7 \\ 270.9 & - \end{bmatrix}$ | - | - | - | - |
| Fig. S2(5×$10^3$) | $\begin{bmatrix} - & 324.7 \\ 360.4 & - \end{bmatrix}$ | $\begin{bmatrix} - & 95.70 \\ 119.1 & - \end{bmatrix}$ | - | - | - | - |
| Fig. S2(×$10^4$) | $\begin{bmatrix} - & 317.2 \\ 362.5 & - \end{bmatrix}$ | $\begin{bmatrix} - & 61.37 \\ 77.32 & - \end{bmatrix}$ | - | - | - | - |
| Fig. S2(5×$10^4$) | $\begin{bmatrix} - & 294.4 \\ 450.4 & - \end{bmatrix}$ | $\begin{bmatrix} - & 28.46 \\ 41.84 & - \end{bmatrix}$ | - | - | - | - |

### S3. Detailed methods description

#### S3.1. Description of the data simulation

##### S3.1.1. Description of the state trajectories

Since in our model we consider the Markovian dynamics for state transitions, we can directly sample the state of the molecule at any given excitation pulse $p$. For the first states, we can directly sample them from the uniform distributions

$$S_{A,1} \sim \mathbf{Cat}_{1,\ldots,M_A}\left(\left[\frac{1}{M_A},\ldots,\frac{1}{M_A}\right]\right) \tag{S1}$$

$$S_{D,1} \sim \mathbf{Cat}_{1,\ldots,M_D}\left(\left[\frac{1}{M_D},\ldots,\frac{1}{M_D}\right]\right) \tag{S2}$$

$$S_{F,1} \sim \mathbf{Cat}_{1,\ldots,M_F}\left(\left[\frac{1}{M_F},\ldots,\frac{1}{M_F}\right]\right) \tag{S3}$$

where, $M_A$, $M_D$ and $M_F$ are the number of acceptor PIFE, donor PIFE and FRET states. Then for sampling the next state we need to calculate the transition probability matrix $\left(\overline{\overline{P}}_A, \overline{\overline{P}}_D, \overline{\overline{P}}_F\right)$, based on the transition rate matrix $\left(\overline{\overline{Q}}_A, \overline{\overline{Q}}_D, \overline{\overline{Q}}_F\right)$

$$\overline{\overline{P}}_A = \exp\left(\overline{\overline{Q}}_A \Delta_p\right) \tag{S4}$$

$$\overline{\overline{P}}_D = \exp\left(\overline{\overline{Q}}_D \Delta_p\right) \tag{S5}$$

$$\overline{\overline{P}}_F = \exp\left(\overline{\overline{Q}}_F \Delta_p\right) \tag{S6}$$

where, $\Delta_p$ is equal to the inter-pulse time. Next, we can sample states conditional on the previous states and marching forward

$$S_{A,p+1}|S_{A,p} \sim \mathbf{Cat}_{1,\ldots,M_A}\left(\overline{P}_{A,S_{A,p}}\right) \tag{S7}$$

$$S_{D,p+1}|S_{D,p} \sim \mathbf{Cat}_{1,\ldots,M_D}\left(\overline{P}_{D,S_{D,p}}\right) \tag{S8}$$

$$S_{F,p+1}|S_{F,p} \sim \mathbf{Cat}_{1,\ldots,M_F}\left(\overline{P}_{F,S_{F,p}}\right) \tag{S9}$$

where, $\overline{P}_{A,S_{A,p}}$ is equal to $S_{A,p}$ row of the transition probability matrix $\overline{\overline{P}}_A$, and the same for the donor PIFE and FRET transitions.

##### S3.1.2. Description of the weights on the detected photons

To generate the simulated data, we consider a complete case when the source of the detected photon can be an excited donor or acceptor, or coming from the background. In this case, we consider the possibility of detecting at most one photon per pulse.

$$W_p \sim \mathbf{Bernoulli}\left([\pi_{w,p}, 1-\pi_{w,p}]\right) \tag{S10}$$

where

$$\pi_{w,p} = \pi_{D,p} + \pi_{A,p} + \pi_{BD} + \pi_{BA}. \tag{S11}$$

Here, $\pi_{D,p}$ is the probability of exciting the donor, $\pi_{A,p}$ is the probability of exciting the acceptor with the donor excitation laser pulse $p$, $\pi_{BD}$ is the probability of collecting a background photon through the donor detector, and $\pi_{BA}$ is the probability of collecting a background photon through the acceptor detector. Here, we assume the probability of two events happen at the same time is negligible and with this assumption these probabilities can be written in the form of

$$\pi_{D,p} = 1 - \exp\left(-\mu_{ex,D,S_{D,p}}\right) \tag{S12}$$
$$\pi_{A,p} = 1 - \exp\left(-\mu_{ex,A,S_{A,p}}\right) \tag{S13}$$
$$\pi_{BD} = 1 - \exp\left(-\Delta_p \mu_{back,D}\right) \tag{S14}$$
$$\pi_{BA} = 1 - \exp\left(-\Delta_p \mu_{back,A}\right) \tag{S15}$$

where, $\{\mu_{ex,D,m_D}\}_{m_D}$ is the donor excitation rate per pulse where $m_D = 1, \dots, M_D$ represent multiple donor excitation rates depending on the state of the molecule due to PIFE effect. Also, we consider the PIFE effect on the acceeptor by setting $\{\mu_{ex,A,m_A}\}_{m_A}$ as the acceptor excitation rates per pulse. In the case of background photon emission rates, we consider different values for the acceptor and donor channels as $\mu_{\text{back},A}$ and $\mu_{\text{back},D}$, respectively.

$$\pi'_{D,p} = \pi_{D,p}\left(1 - \pi_{A,p}\right)\left(1 - \pi_{BD}\right)\left(1 - \pi_{BA}\right) \tag{S16}$$
$$\pi'_{A,p} = \pi_{A,p}\left(1 - \pi_{D,p}\right)\left(1 - \pi_{BD}\right)\left(1 - \pi_{BA}\right) \tag{S17}$$
$$\pi'_{BD,p} = \pi_{BD}\left(1 - \pi_{D,p}\right)\left(1 - \pi_{A,p}\right)\left(1 - \pi_{BA}\right) \tag{S18}$$
$$\pi'_{BA,p} = \pi_{BA}\left(1 - \pi_{D,p}\right)\left(1 - \pi_{A,p}\right)\left(1 - \pi_{BD}\right) \tag{S19}$$

##### S3.1.3. Description of the probability on detection channel

After sampling the photon detection, we only keep the pules that we detect a photon ($W_p = 1$) and accordingly we only keep the part of state trajectories in which we detect photons. We show the index of such photons with $t$. So ,next we need to sample the source of that photon, which can be donor, acceptor, donor background or acceptor background, the tag on the photon at pulse $p$ is equal to

$$\gamma_t | \overline{\pi'}_t \sim \mathbf{Cat}_{D,A,BD,BA}\left(\left[\pi'_{D,t}, \pi'_{A,t}, \pi'_{BD,t}, \pi'_{BA,t}\right]\right) \tag{S20}$$

where, $\overline{\pi'}_t = \left(\pi'_{D,t}, \pi'_{A,t}, \pi'_{BD,t}, \pi'_{BA,t}\right)$ are the normalized weights at detected photon $t$.

Next, we need to consider the case that the donor is excited and there are several pathways that energy can be transferred. In this case, we can have the donor inverse lifetime through time $k_{D,t}$, and resonance energy transfer rate $k_{F,t}$, or transfer the energy to the acceptor.

$$q_t | \gamma_t, k_{D,t}, k_{F,t} \sim \mathbb{I}\left(\gamma_t = D\right) \mathbf{Cat}_{D,A}\left(\left[\pi_1, \pi_2\right]\right) + A\mathbb{I}\left(\gamma_t = A\right) + D\mathbb{I}\left(\gamma_t = BD\right) + A\mathbb{I}\left(\gamma_t = BA\right) \tag{S21}$$

where

$$\pi_1 = \frac{\frac{1}{\tau_{D,S_{D,t}}}}{\frac{1}{\tau_{D,S_{D,t}}} + k_{F,S_{F,t}}} \tag{S22}$$

$$\pi_2 = \frac{k_{F,S_{F,t}}}{\frac{1}{\tau_{D,S_{D,t}}} + k_{F,S_{F,t}}} \tag{S23}$$

and by marginalization over $\gamma_t$ we have

$$q_t | \overline{\pi'}_t, \tau_{D,S_{D,t}}, k_{F,S_{F,t}} \sim \mathbf{Cat}_{D,A}\left(\left[\pi''_1, \pi''_2\right]\right) \tag{S24}$$

where

$$\pi''_t = \pi'_{D,t}\left(\frac{\frac{1}{\tau_{D,S_{D,t}}}}{K_t}\right) + \pi'_{BD,t} \tag{S25}$$

$$\pi''_t = \pi'_{D,t}\left(\frac{k_{F,S_{F,t}}}{K_t}\right) + \pi'_{A,t} + \pi'_{BA,t} \tag{S26}$$

where, $K_t = \frac{1}{\tau_{D,S_{D,t}}} + k_{F,S_{F,t}}$. Also, we need to incorporate the uniform background detection possibility into account. As the result we have

At top of the photon labels we have the detection efficiency which correspond to our capability of separating/filtering photons based on their wavelengths. In this case the photon detection is

$$c_t|q_t, \eta_A, \eta_D, \overline{\pi'}_t, \tau_{D,S_{D,t}}, k_{F,S_{F,t}} \sim \begin{cases} \mathbf{Cat}_{D,A}\left([1-\eta_D, \eta_D]\right) & q_t = D \\ \mathbf{Cat}_{D,A}\left([\eta_A, 1-\eta_A]\right) & q_t = A \end{cases} \tag{S27}$$

where, $\eta_D$ is the probability of cross-talk of donor photons into the acceptor channel and $\eta_A$ is the probability of cross-talk of acceptor photons into the detector channel.

Here, we can simplify the formulation by marginalizing over $q_t$ and as the result we have

$$c_t|\eta_A, \eta_D, \overline{\pi'}_t, \tau_{D,S_{D,t}}, k_{F,S_{F,t}} \sim \mathbf{Cat}_{D,A}\left([\pi''_1, \pi''_2]\begin{bmatrix} 1-\eta_D & \eta_D \\ \eta_A & 1-\eta_A \end{bmatrix}\right) \tag{S28}$$

where, $\overline{\pi'}_p$ is a function of acceptor and donor background photon emission rates $\mu_{\text{back},A}$, $\mu_{\text{back},D}$, and acceptor and donor excitation rate per pulse corresponding to the donor and acceptor PIFE state $\mu_{ex,A,S_{A,t}}$, $\mu_{ex,D,S_{D,t}}$.

##### S3.1.4. Description of the probability on photon arrival times

After sampling the source and detection channel of detected photons, at the next step we need to construct the probability of photon arrival times from the photons based on their source and channel we detect them. In the case that we detected a photon from the donor channel, the distribution of delay times after the excitation time $t_{ex}$ has a form of

$$\delta_t - t_{ex}|\tau_{D,S_{D,t}}, k_{F,S_{F,t}} \sim \mathbf{Exp}\left(\frac{1}{\frac{1}{\tau_{D,S_{D,t}}} + k_{F,S_{F,t}}}\right) \tag{S29}$$

where $\frac{1}{\frac{1}{\tau_{D,S_{D,t}}}+k_{F,S_{F,t}}}$ is equal to the lifetime of the donor conditional on its PIFE state $S_{D,t}$ and the resonance energy transition rate $k_{F,S_{F,t}}$ corresponding to its FRET state $S_{F,t}$. Next, we have to consider the effect of IRF, which in this case we approximate it with a Gaussian distribution

$$t_{ex} \sim \mathbf{Normal}\left(\mu_{IRF}, \sigma^2_{IRF}\right) \tag{S30}$$

where, $\mu_{IRF}$ is the mean value of the fitted IRF (equivalent to the offset of the instrumental photon counter), and $\sigma_{IRF}$ is the standard deviation of the fitted IRF which is the result of an error caused by the excitation pulse and the instrumental photon counter. Here, by having the convolution over the excitation time $t_{ex}$ we have

$$p\left(\delta_t\right) = \int_0^{\delta_t} \mathbf{Normal}\left(t_{ex}; \mu_{IRF}, \sigma^2_{IRF}\right) \mathbf{Exp}\left(\delta_t - t_{ex}; \frac{1}{\frac{1}{\tau_{D,S_{D,t}}} + k_{F,S_{F,t}}}\right) dt_{ex} \tag{S31}$$

which we call this probability $P^D_{D^*D}(.)$

$$P^D_{D^*D}(.) = \frac{K_t}{2} e^{K_t\left(\mu_{\text{IRF},D} - \delta_t + \frac{\sigma^2_{\text{IRF},D}}{2}K_t\right)} \left[erf\left(\frac{\mu_{\text{IRF},D} + K_t\sigma^2_{\text{IRF},D}}{\sigma_{\text{IRF},D}\sqrt{2}}\right) - erf\left(\frac{\mu_{\text{IRF},D} - \delta_t + K_t\sigma^2_{\text{IRF},D}}{\sigma_{\text{IRF},D}\sqrt{2}}\right)\right]. \tag{S32}$$

Here, based on the approximation on the ingratiation limit and instead of integral from zero, we can consider starting from $-\infty$, and as a result we can simplify above equation to

$$P^D_{D^*D}(.) = \frac{K_t}{2} \exp\left(K_t\left(\mu_{\text{IRF},D} - \delta_t + \frac{\sigma^2_{\text{IRF},D}}{2}K_t\right)\right) erfc\left(\frac{\mu_{\text{IRF},D} - \delta_t + K_t\sigma^2_{\text{IRF},D}}{\sigma_{\text{IRF},D}\sqrt{2}}\right). \tag{S33}$$

The same analogy can be applied to calculate the probability distribution of the photon with the source of the donor excitation emission resonance energy transfer (FRET), and acceptor emission and detection. For thiss case the equation Eq. (S29) will change to

$$t_F - t_{ex} \sim \mathbf{Exp}\left(\frac{1}{K_t}\right) \tag{S34}$$

$$\delta_t - t_F \sim \mathbf{Exp}\left(\tau_{A,S_{A,t}}\right) \tag{S35}$$

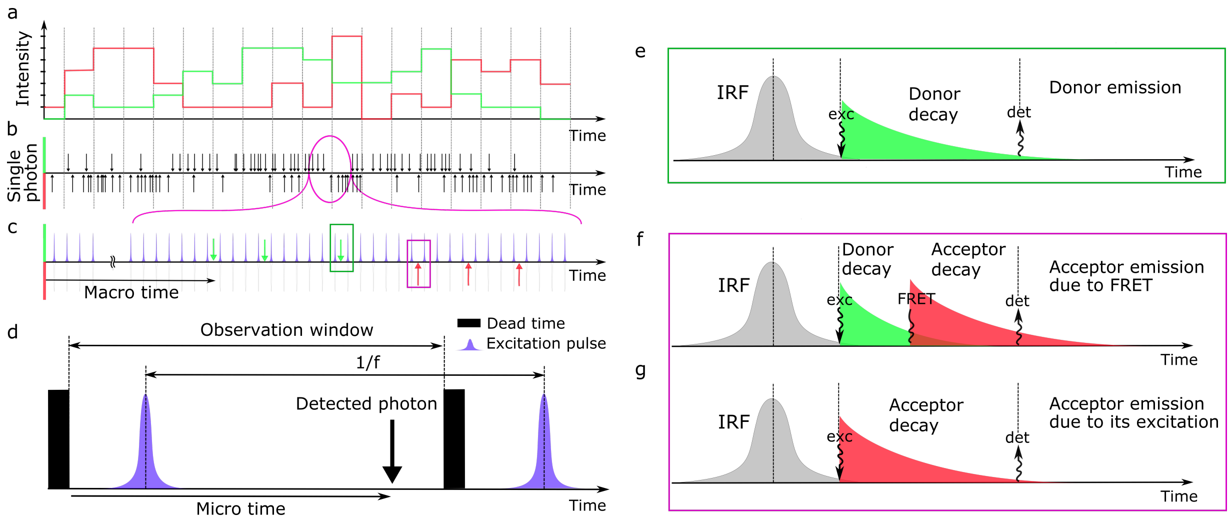


FIG. S3. **A cartoon representation of the photon delay times distribution.** **(a)** The Intensity signal of the donor and acceptor channels as the result of collections of photons peer bin size. **(b)** The single photon arrival time traces of donor and acceptor channels corresponding to intensity traces in (a). **(c)** The magnified portion of the single photon arrival time traces in (b) illustrates the positions of the photons with respect to pulse excitations. **(d)** Illustration of the camera observation window, dead-time and excitation pulse versus the photon detection time. **(e)** Illustration of IRF and donor decay time. in this case, the source of the photon is from the excited donor and the emission from the donor. **(e)** Illustration of IRF, FRET, and acceptor decay time. in this case, the source of the photon is the excited donor and emission is from the acceptor due to FRET. **(g)** Illustration of IRF and acceptor decay time. in this case, the source of the photon is the excited acceptor and emission is from the acceptor as well.

where, $t_F$ is the time that the energy transfer to the acceptor, and by marginalization over the $t_F$ we have

$$\delta_t - t_{ex} \sim f_{D^*,A}(.) \tag{S36}$$

where

$$f_{D^*A}(.) = \int_{t_{ex}}^{\delta_t} \textbf{Exp}\left(\frac{1}{K_t}\right) \textbf{Exp}\left(\tau_{A,S_{A,t}}\right) dt_F = \frac{\exp\left(\frac{t_{ex}-\delta_t}{\tau_{A,S_{A,t}}}\right) - \exp\left(K_t\left(t_{ex}-\delta_t\right)\right)}{\tau_{A,S_{A,t}} - \frac{1}{K_t}}. \tag{S37}$$

Next, by marginalizing over excitation distribution we have

$$\begin{aligned} P^A_{D^*A}(.) &= \int_0^{\delta_t} \textbf{Normal}\left(t_{ex}; \mu_{\text{IRF},A}, \sigma^2_{\text{IRF},A}\right) f_{D^*A}(.) dt_{ex} = \int_0^{\delta_t} \frac{e^{-\frac{(t_{ex}-\mu-\text{IRF},A)^2}{2\sigma^2_{\text{IRF},A}}}}{\sqrt{2\pi\sigma^2_{\text{IRF},A}}} \frac{e^{\frac{t_{ex}-\delta_t}{\tau_{A,S_{A,t}}}} - e^{K_t(t_{ex}-\delta_t)}}{\tau_{A,S_{A,t}} - \frac{1}{K_t}} dt_{ex} \\ &\approx \int_{-\infty}^{\delta_t} \frac{e^{-\frac{(t_{ex}-\mu-\text{IRF},A)^2}{2\sigma^2_{\text{IRF},A}}}}{\sqrt{2\pi\sigma^2_{\text{IRF},A}}} \frac{e^{\frac{t_{ex}-\delta_t}{\tau_{A,S_{A,t}}}} - e^{K_t(t_{ex}-\delta_t)}}{\tau_{A,S_{A,t}} - \frac{1}{K_t}} dt_{ex} \\ &= \frac{0.5}{\tau_{A,S_{A,t}} - \frac{1}{K_t}} \left[ \begin{array}{l} \exp\left(\frac{\sigma^2_{\text{IRF},A} + 2\tau_{A,S_{A,t}}\left(\mu_{\text{IRF},A}-\delta_t\right)}{2\left(\tau_{A,S_{A,t}}\right)^2}\right) erfc\left(\frac{\sigma^2_{\text{IRF},A} + \tau_{A,S_{A,t}}\left(\mu_{\text{IRF},A}-\delta_t\right)}{\sigma_{\text{IRF},A}\tau_{A,S_{A,t}}\sqrt{2}}\right) \\ -\exp\left(K_t\left(\mu_{\text{IRF},A} - \delta_t + \frac{\sigma^2_{\text{IRF},A}}{2}K_t\right)\right) erfc\left(\frac{\sigma^2_{\text{IRF},A}K_t + \left(\mu_{\text{IRF},A}-\delta_t\right)}{\sigma_{\text{IRF},A}\sqrt{2}}\right) \end{array} \right]. \end{aligned} \tag{S38}$$

Also, for the case of acceptor excitation and emission, if detection happen in acceptor channel we have

$$\begin{aligned} P^A_{A^*A}(.) &= \int_0^{\delta_t} \textbf{Normal}\left(t_{ex}; \mu_{\text{IRF},A}, \sigma^2_{\text{IRF},A}\right) \textbf{EXP}\left(\delta_t - t_{ex}; \tau^0_A\right) dt_{ex} \\ &= \frac{1}{2\tau_{A,S_{A,t}}} \exp\left(\frac{1}{\tau_{A,S_{A,t}}}\left(\mu_{\text{IRF},A} - \delta_t + \frac{\sigma^2_{\text{IRF},A}}{2\tau_{A,S_{A,t}}}\right)\right) erfc\left(\frac{\mu_{\text{IRF},A} - \delta_t + \frac{\sigma^2_{\text{IRF},A}}{\tau_{A,S_{A,t}}}}{\sigma_{\text{IRF},A}\sqrt{2}}\right). \end{aligned} \tag{S39}$$

Now, we need to consider the scenarios that photons collect in other detectors. In these cases, the mean and standard deviation of the IRFs should change accordingly. For example, in the case of acceptor excitation and emission if the detection happen in donor channel, we have

$$P^{D}_{A^*A}(.) = \frac{1}{2\tau_{A,S_{A,t}}} \exp\left(\frac{1}{\tau_{A,S_{A,t}}}\left(\mu_{\text{IRF},D} - \delta_t + \frac{\sigma^2_{\text{IRF},D}}{2\tau_{A,S_{A,t}}}\right)\right) erfc\left(\frac{\mu_{\text{IRF},D} - \delta_t + \frac{\sigma^2_{\text{IRF},D}}{\tau_{A,S_{A,t}}}}{\sigma_{\text{IRF},D}\sqrt{2}}\right). \tag{S40}$$

Finally, we need to encounter the effect of the cross-talks between the detectors. In this case, donor or acceptor photons have a chance to be detected by the other detector. These events happen due to the distribution of the emission wavelength of dyes and imperfection of the filters. As the result, we have

$$\delta_t|\eta_A, \eta_D, \mu_{\text{back},A}, \mu_{\text{back},D}, \{\tau_{A,m_A}, \mu_{ex,A,m_A}\}_{m_A}, \{\tau_{D,m_D}, \mu_{ex,D,m_D}\}_{m_D}, \{k_{F,m_F}\}_{m_F}, S_{A,t}, S_{D,t}, S_{F,t}, C_t, \sim \tag{S41}$$

$$\begin{cases} \left(\pi'_D \frac{\frac{1}{\tau_{D,S_{D,t}}}}{K_t} P^{D}_{D^*D,t}(.) + \pi'_{BD}P_B(.)\right)(1-\eta_D) + \left(\pi'_D \frac{k_{F,S_{F,t}}}{K_t} P^{D}_{D^*A,t}(.) + \pi'_A P^{D}_{A^*A,t}(.) + \pi'_{BA}P_B(.)\right)\eta_A \\ \qquad c_t = 1 \text{ (Donor channel)} \\ \left(\pi'_D \frac{\frac{1}{\tau_{D,S_{D,t}}}}{K_t} P^{A}_{D^*D,t}(.) + \pi'_{BD}P_B(.)\right)\eta_D + \left(\pi'_D \frac{k_{F,S_{F,t}}}{K_t} P^{A}_{D^*A,t}(.) + \pi'_A P^{A}_{A^*A,t}(.) + \pi'_{BA}P_B(.)\right)(1-\eta_A) \\ \qquad c_t = 2 \text{ (Acceptor channel).} \end{cases} \tag{S42}$$

where, $\eta_A$ is the detection cross-talk probability of acceptor photons, $\eta_D$ is the detection cross-talk probability of donor photons, and $\overline{\pi}' = (\pi'_D, \pi'_A, \pi'_{BD}, \pi'_{BA})$ is the set of weights on the photons mentioned in Eqs. (S16)-(S19). Also, this equation needs to be normalized and as a result is equal to:

$$\delta_t|\eta_A, \eta_D, \mu_{\text{back},A}, \mu_{\text{back},D}, \{\tau_{A,m_A}, \mu_{ex,A,m_A}\}_{m_A}, \{\tau_{D,m_D}, \mu_{ex,D,m_D}\}_{m_D}, \{k_{F,m_F}\}_{m_F}, S_{A,t}, S_{D,t}, S_{F,t}, C_t \sim \tag{S43}$$

$$\begin{cases} \dfrac{\left(\pi'_D \frac{\frac{1}{\tau_{D,S_{D,t}}}}{K_t} P^{D}_{D^*D,t}(.) + \pi'_{BD}P_B(.)\right)(1-\eta_D) + \left(\pi'_D \frac{k_{F,S_{F,t}}}{K_t} P^{D}_{D^*A,t}(.) + \pi'_A P^{D}_{A^*A,t}(.) + \pi'_{BA}P_B(.)\right)\eta_A}{\left(\pi'_D \frac{\frac{1}{\tau_{D,S_{D,t}}}}{K_t} + \pi'_{BD}\right)(1-\eta_D) + \left(\pi'_D \frac{k_{F,S_{F,t}}}{K_t} + \pi'_A + \pi'_{BA}\right)\eta_A} \\ \qquad c_t = 1 \text{ (Donor channel)} \\ \dfrac{\left(\pi'_D \frac{\frac{1}{\tau_{D,S_{D,t}}}}{K_t} P^{A}_{D^*D,t}(.) + \pi'_{BD}P_B(.)\right)\eta_D + \left(\pi'_D \frac{k_{F,S_{F,t}}}{K_t} P^{A}_{D^*A,t}(.) + \pi'_A P^{A}_{A^*A,t}(.) + \pi'_{BA}P_B(.)\right)(1-\eta_A)}{\left(\pi'_D \frac{\frac{1}{\tau_{D,S_{D,t}}}}{K_t} + \pi'_{BD}\right)\eta_D + \left(\pi'_D \frac{k_{F,S_{F,t}}}{K_t} + \pi'_A + \pi'_{BA}\right)(1-\eta_A)} \\ \qquad c_t = 2 \text{ (Acceptor channel).} \end{cases} \tag{S44}$$

where, we consider the case that

$$\int_0^\infty P^{D}_{D^*D,t}(.)d\delta_t \approx 1 \tag{S45}$$

$$\int_0^\infty P^{A}_{D^*D,t}(.)d\delta_t \approx 1 \tag{S46}$$

$$\int_0^\infty P^{D}_{D^*A,t}(.)d\delta_t \approx 1 \tag{S47}$$

$$\int_0^\infty P^{A}_{D^*A,t}(.)d\delta_t \approx 1 \tag{S48}$$

$$\int_0^\infty P^{D}_{A^*A,t}(.)d\delta_t \approx 1 \tag{S49}$$

$$\int_0^\infty P^{A}_{A^*A,t}(.)d\delta_t \approx 1 \tag{S50}$$

$$\int_0^\infty P_B(.)d\delta_t \approx 1. \tag{S51}$$

These assumptions allow us to have simpler version of likelihood as

$$
\begin{aligned}
&p\left(\delta_t|\eta_A,\eta_D,\mu_{\text{back},A},\mu_{\text{back},D},\{\tau_{A,m_A},\mu_{ex,A,m_A}\}_{m_A},\{\tau_{D,m_D},\mu_{ex,D,m_D}\}_{m_D},\{k_{F,m_F}\}_{m_F},S_{A,t},S_{D,t},S_{F,t},C_t\right)\\
&\quad\times p\left(C_t|\eta_A,\eta_D,\mu_{\text{back},A},\mu_{\text{back},D},\{\mu_{ex,A,m_A}\}_{m_A},\{\tau_{D,m_D},\mu_{ex,D,m_D}\}_{m_D},\{k_{F,m_F}\}_{m_F},S_{A,t},S_{D,t},S_{F,t}\right)\sim\\
&\begin{cases}
\left(\pi'_D\dfrac{\frac{1}{\tau_{D,S_{D,t}}}}{K_t}P^D_{D^*D,t}(.)+\pi'_{BD}P_B(.)\right)(1-\eta_D)+\left(\pi'_D\dfrac{k_{F,S_{F,t}}}{K_t}P^D_{D^*A,t}(.)+\pi'_AP^D_{A^*A,t}(.)+\pi'_{BA}P_B(.)\right)\eta_A\\
\qquad\qquad C_t=1\text{ (Donor channel)}\\
\left(\pi'_D\dfrac{\frac{1}{\tau_{D,S_{D,t}}}}{K_t}P^A_{D^*D,t}(.)+\pi'_{BD}P_B(.)\right)\eta_D+\left(\pi'_D\dfrac{k_{F,S_{F,t}}}{K_t}P^A_{D^*A,t}(.)+\pi'_AP^A_{A^*A,t}(.)+\pi'_{BA}P_B(.)\right)(1-\eta_A)\\
\qquad\qquad C_t=2\text{ (Acceptor channel).}
\end{cases}
\end{aligned}
\tag{S52}
$$

Also, due to consideration of differences in donor and acceptor channel IRFs, we have

$$P^D_{D^*D,t}(.)=\frac{K_t}{2}\exp\left(K_t\left(\mu_{D,t}+\frac{\sigma^2_{\text{IRF},D}}{2}K_t\right)\right)erfc\left(\frac{\mu_{D,t}+\sigma^2_{\text{IRF},D}K_t}{\sigma_{\text{IRF},D}\sqrt{2}}\right)\tag{S53}$$

$$P^D_{D^*A,t}(.)=\frac{0.5}{\tau_{A,S_{A,t}}-\frac{1}{K_t}}\left[\begin{aligned}&\exp\left(\frac{\sigma^2_{\text{IRF},D}+2\tau_{A,S_{A,t}}\mu_{D,t}}{2\left(\tau_{A,S_{A,t}}\right)^2}\right)erfc\left(\frac{\mu_{D,t}+\frac{\sigma^2_{\text{IRF},D}}{\tau_{A,S_{A,t}}}}{\sigma_{\text{IRF},D}\sqrt{2}}\right)\\&-\exp\left(K_t\left(\mu_{D,t}+\frac{\sigma^2_{\text{IRF},D}}{2}K_t\right)\right)erfc\left(\frac{\mu_{D,t}+\sigma^2_{\text{IRF},D}K_t}{\sigma_{\text{IRF},D}\sqrt{2}}\right)\end{aligned}\right]\tag{S54}$$

$$P^A_{D^*A,t}(.)=\frac{0.5}{\tau_{A,S_{A,t}}-\frac{1}{K_t}}\left[\begin{aligned}&\exp\left(\frac{\sigma^2_{\text{IRF},A}+2\tau_{A,S_{A,t}}\mu_{A,t}}{2\left(\tau_{A,S_{A,t}}\right)^2}\right)erfc\left(\frac{\mu_{A,t}+\frac{\sigma^2_{\text{IRF},A}}{\tau_{A,S_{A,t}}}}{\sigma_{\text{IRF},A}\sqrt{2}}\right)\\&-\exp\left(K_t\left(\mu_{A,t}+\frac{\sigma^2_{\text{IRF},A}}{2}K_t\right)\right)erfc\left(\frac{\mu_{A,t}+\sigma^2_{\text{IRF},A}K_t}{\sigma_{\text{IRF},A}\sqrt{2}}\right)\end{aligned}\right]\tag{S55}$$

$$P^A_{D^*D,t}(.)=\frac{K_t}{2}\exp\left(K_t\left(\mu_{A,t}+\frac{\sigma^2_{\text{IRF},A}}{2}K_t\right)\right)erfc\left(\frac{\mu_{A,t}+\sigma^2_{\text{IRF},A}K_T}{\sigma_{\text{IRF},A}\sqrt{2}}\right)\tag{S56}$$

$$P^A_{A^*A,t}(.)=\frac{1}{2\tau_{A,S_{A,t}}}\exp\left(\frac{\mu_{A,t}+\frac{\sigma^2_{\text{IRF},A}}{2\tau_{A,S_{A,t}}}}{\tau_{A,S_{A,t}}}\right)erfc\left(\frac{\mu_{A,t}+\frac{\sigma^2_{\text{IRF},A}}{\tau_{A,S_{A,t}}}}{\sigma_{\text{IRF},A}\sqrt{2}}\right)\tag{S57}$$

$$P^D_{A^*A,t}(.)=\frac{1}{2\tau_{A,S_{A,t}}}\exp\left(\frac{\mu_{D,t}+\frac{\sigma^2_{\text{IRF},D}}{2\tau_{A,S_{A,t}}}}{\tau_{A,S_{A,t}}}\right)erfc\left(\frac{\mu_{D,t}+\frac{\sigma^2_{\text{IRF},D}}{\tau_{A,S_{A,t}}}}{\sigma_{\text{IRF},D}\sqrt{2}}\right)\tag{S58}$$

$$P_B(.)=\mathbf{Uniform}\left([0,T_p]\right).\tag{S59}$$

where, $\mu_{D,t}=\mu_{\text{IRF},D}-\delta_t$, $\mu_{A,t}=\mu_{\text{IRF},A}-\delta_t$, and $K_t=k_{F,S_{F,t}}+\frac{1}{\tau_{D,S_{D,t}}}$.
Detailed parameter choices for all simulations performed are listed in Table S6.

### S4. Detailed description of the inference framework

#### S4.1. Description of prior probability distributions

The model parameters in our framework that require priors are: acceptor and donor background photon emission rates $\mu_{\text{back},A}$, $\mu_{\text{back},D}$; acceptor and donor channel cross-talk ratios $\eta_A$, $\eta_D$; acceptor and donor excitation rates $\{\mu_{ex,A,m_A}\}_{m_A}$, $\{\mu_{ex,D,m_D}\}_{m_D}$; acceptor and donor lifetimes $\{\tau_{A,m_A}\}_{m_A}$, $\{\tau_{D,m_D}\}_{m_D}$ and ET rates $\{k_{F,m_F}\}_{m_F}$; hyper

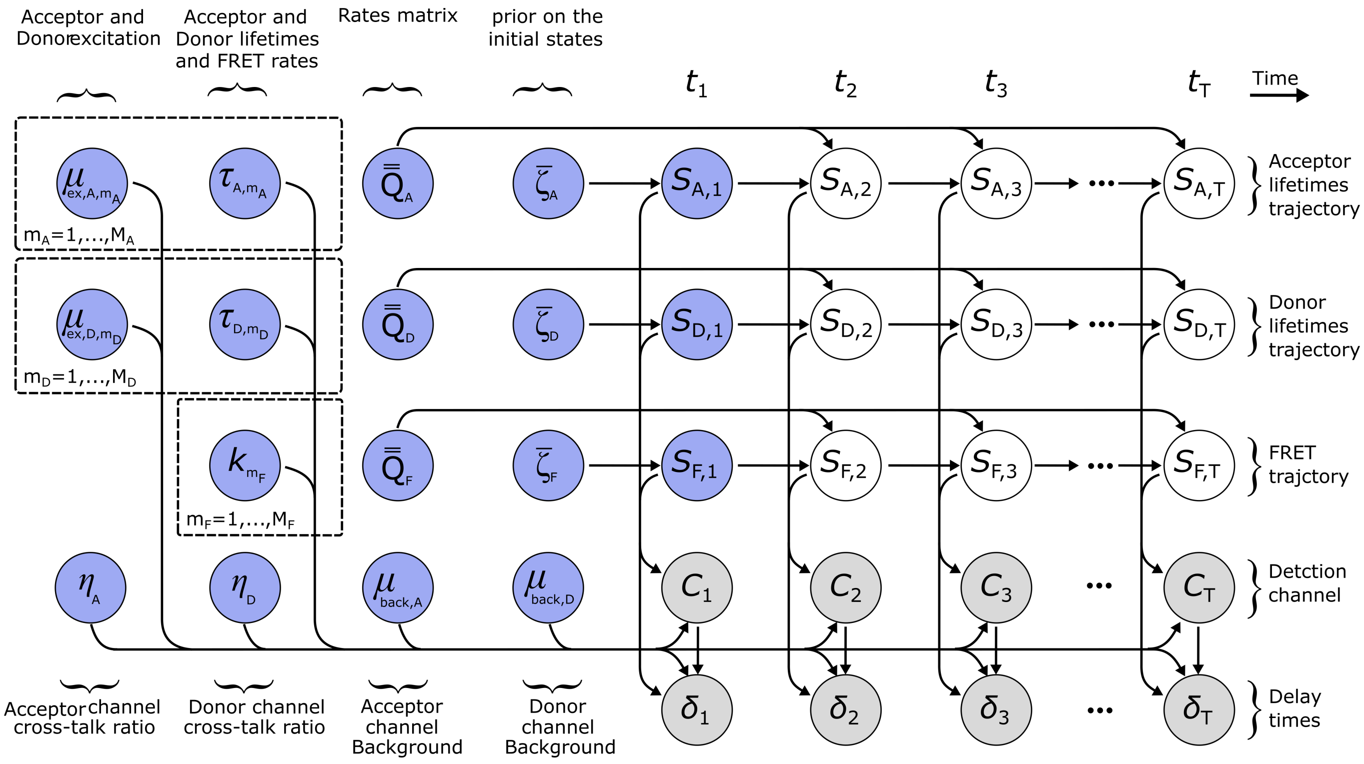


FIG. S4. **Graphical summary of the framework.** The model molecule evolves over the course of the experiment which is marked by t=1,2,...,T. Here, $(S_{A,t}, S_{D,t}, S_{F,t})$ denote the acceptor-PIFE, donor-PIFE and FRET state trajectories at time $t$; During the experiment, $\delta_t$, is the delay time of the detected photon in channel $C_t$. $\{\tau_{A,m_A}\}_{m_A}$ are the acceptor lifetimes for each acceptor-PIFE state $m_A$, $\{\tau_{D,m_D}\}_{m_D}$ are the donor lifetimes for each donor-PIFE state $m_D$, and $\{k_{F,m_F}\}_{m_F}$ are the ET rates of each FRET states $m_F$. The $\overline{\overline{Q}}_A$, $\overline{\overline{Q}}_D$ and $\overline{\overline{Q}}_F$ are transition rates matrix for acceptor and donor lifetimes and FRET trajectories. $\overline{\zeta}_A$, $\overline{\zeta}_D$ and $\overline{\zeta}_F$ are the weights on the initial states of acceptor-PIFE, donor-PIFE and FRET, respectively. To incorporate the cross-talks in detectors, we have $\eta_A$ and $\eta_D$ for acceptor and donor channels. The background photon emission rates in channels are shown with $\mu_{\text{back},A}$ and $\mu_{\text{back},D}$. The $\{\mu_{ex,A,m_A}\}_{m_A}$ and $\{\mu_{ex,D,m_D}\}_{m_D}$ are the acceptor and donor excitation rates corresponding to their states.

prior weights over the acceptor states $\overline{\beta}_{A,hyper}$, donor states $\overline{\beta}_{D,hyper}$, and FRET states $\overline{\beta}_{F,hyper}$; initial states of acceptor $S_{A,1}$, donor $S_{D,1}$, and FRET $S_{F,1}$; acceptor escape rates $\{\lambda_{A,m_A}\}_{m_A}$ and their ratio of escape rates $\{\overline{\pi}_{A,m_A}\}_{m_A}$; donor escape rates $\{\lambda_{D,m_D}\}_{m_D}$ and their ratio of escape rates $\{\overline{\pi}_{D,m_D}\}_{m_D}$; and FRET escape rates $\{\lambda_{F,m_F}\}_{m_F}$ and their ratio of escape rates $\{\overline{\pi}_{F,m_F}\}_{m_F}$. Our choices are described below.

###### S4.1.1. Prior on acceptor and donor channels background photon emission rates $\mu_{back,A}$ and $\mu_{back,D}$

Since the acceptor and donor channels background photon emission rates are positive values, we consider gamma distributions prior for them.

$$\mu_{\text{back},A} \sim \mathbf{Gamma}\left(\alpha_{\text{back},A}, \beta_{\text{back},A}\right) \tag{S60}$$

$$\mu_{\text{back},D} \sim \mathbf{Gamma}\left(\alpha_{\text{back},D}, \beta_{\text{back},D}\right) \tag{S61}$$

where, $\alpha_{\text{back},A}$ and $\beta_{\text{back},A}$ are the parameters of gamma prior on acceptor channel background, and the $\alpha_{\text{back},D}$ and $\beta_{\text{back},D}$ are parameters of the gamma prior on the donor channel background.

###### S4.1.2. Priors on the acceptor and donor channel cross-talk ratios $\eta_A$, $\eta_D$

To ensure that $\eta_A$ and $\eta_D$ sampled in our formulation attain only positive values between 0 and 1, we place a Beta priors on both

$$\begin{aligned} \eta_A &\sim \mathbf{Beta}\left(\alpha_{\eta,D}, \beta_{\eta,D}\right) \\ \eta_D &\sim \mathbf{Beta}\left(\alpha_{\eta,A}, \beta_{\eta,A}\right). \end{aligned} \tag{S62}$$

###### S4.1.3. Prior on acceptor and donor excitation rates $\{\mu_{ex,A,m_A}\}_{m_A}$ and $\{\mu_{ex,D,m_D}\}_{m_D}$

Since the acceptor and donor excitation rates are positive values, we consider gamma distributions prior for them.

$$\mu_{ex,A,m_A} \sim \mathbf{Gamma}\left(\alpha_{\mu_A}, \beta_{\mu_A}\right), \qquad m_A = 1, \ldots, M_A \tag{S63}$$

$$\mu_{ex,D,m_D} \sim \mathbf{Gamma}\left(\alpha_{\mu_D}, \beta_{\mu_D}\right), \qquad m_D = 1, \ldots, M_D \tag{S64}$$

where, $\alpha_{\mu_A}$, $\beta_{\mu_A}$, $\alpha_{\mu_D}$ and $\beta_{\mu_D}$ are parameters of the gamma prior.

###### S4.1.4. Priors on the acceptor and donor lifetimes $\{\tau_{A,m_A}\}_{m_A}$ and $\{\tau_{D,m_D}\}_{m_D}$ and ET rates $\{k_{F,m_F}\}_{m_F}$

To ensure that $\{\tau_{A,m_A}\}_{m_A}$, $\{\tau_{D,m_D}\}_{m_D}$ and $\{k_{F,m_F}\}_{m_F}$ sampled in our formulation attain only positive values, we place Gamma priors on them.

$$\tau_{A,m_A} \sim \mathbf{Gamma}\left(\alpha_A, \beta_A\right), \qquad m_A = 1, \ldots, M_A \tag{S65}$$

$$\tau_{D,m_D} \sim \mathbf{Gamma}\left(\alpha_D, \beta_D\right), \qquad m_D = 1, \ldots, M_D \tag{S66}$$

$$k_{F,m_F} \sim \mathbf{Gamma}\left(\alpha_F, \beta_F\right), \qquad m_F = 1, \ldots, M_F \tag{S67}$$

where, $\alpha_A$, $\beta_A$, $\alpha_D$,$\beta_D$, $\alpha_F$ and $\beta_F$ are parameters of the gamma prior.

###### S4.1.5. Prior on the initial states acceptor $S_{A,1}$, donor $S_{D,1}$, and FRET $S_{F,1}$

The initial states need a prior which has a dependency on the corresponding weights on each state. As the result, we have

$$S_{A,1}|\overline{\zeta}_A \sim \mathbf{Cat}_{1,\ldots,M_A}\left(\overline{\beta}_A\right) \tag{S68}$$

$$S_{D,1}|\overline{\zeta}_D \sim \mathbf{Cat}_{1,\ldots,M_D}\left(\overline{\beta}_D\right) \tag{S69}$$

$$S_{F,1}|\overline{\zeta}_F \sim \mathbf{Cat}_{1,\ldots,M_F}\left(\overline{\beta}_F\right) \tag{S70}$$

where, $\overline{\zeta}_A$, $\overline{\zeta}_D$, and $\overline{\zeta}_F$ are the weights on the first states.

###### S4.1.6. Prior on the wights of initial states for acceptor-PIFE $\overline{\zeta}_A$, donor-PIFE $\overline{\zeta}_D$, and FRET $\overline{\zeta}_F$

In order to have a discrete distribution over these weights, we consider Dirichlet distributions

$$\overline{\zeta}_A \sim \mathbf{Dir}\left(\alpha_{A,\zeta}\overline{\beta}_{A,hyper}\right) \tag{S71}$$

$$\overline{\zeta}_D \sim \mathbf{Dir}\left(\alpha_{D,\zeta}\overline{\beta}_{D,hyper}\right) \tag{S72}$$

$$\overline{\zeta}_F \sim \mathbf{Dir}\left(\alpha_{F,\zeta}\overline{\beta}_{F,hyper}\right) \tag{S73}$$

where, $\alpha_{A,\zeta}$, $\alpha_{D,\zeta}$, and $\alpha_{F,\zeta}$ are the concentration parameters, and $\overline{\beta}_{A,hyper}$, $\overline{\beta}_{D,hyper}$ and $\overline{\beta}_{F,hyper}$ are the base distributions of the priors which can be set to uniform distributions.

##### S4.1.7. Prior on the elements of state transition rate matrices $\overline{\overline{Q}}_A$, $\overline{\overline{Q}}_D$, $\overline{\overline{Q}}_F$

Since the diagonal elements of the transition matrix are the negative sum of each row, we have in total $M_A\,(M_A-1)$, $M_D\,(M_D-1)$ and $M_F\,(M_F-1)$ transition rates for state transition rate matrices $\overline{\overline{Q}}_A$, $\overline{\overline{Q}}_D$, $\overline{\overline{Q}}_F$, respectively. To ensure that these rates are positive, we consider gamma distribution

$$\lambda_{A,m_A} \sim \mathbf{Gamma}\left(\alpha_{\lambda_A}, \beta_{\lambda_A}\right) \qquad , m_A = 1, \ldots, M_A\,(M_A-1) \tag{S74}$$
$$\lambda_{D,m_D} \sim \mathbf{Gamma}\left(\alpha_{\lambda_D}, \beta_{\lambda_D}\right) \qquad , m_D = 1, \ldots, M_D\,(M_D-1) \tag{S75}$$
$$\lambda_{F,m_F} \sim \mathbf{Gamma}\left(\alpha_{\lambda_F}, \beta_{\lambda_F}\right) \qquad , m_F = 1, \ldots, M_F\,(M_F-1) \tag{S76}$$

where, $M_A$ is the total number of acceptor-PIFE states, $M_D$ is the total number of donor-PIFE states, $M_F$ is the total number of FRET states, and $\alpha_{\lambda_A}$, $\beta_{\lambda_A}$, $\alpha_{\lambda_D}$, $\beta_{\lambda_D}$, $\alpha_{\lambda_F}$ and$\beta_{\lambda_F}$ are the prior parameters.

In order to be able to directly sample the transition rates, we can discretize the transition rate matrix into two parts. first, a diagonal matrix that contains the escape rates from states; second, the ratio of escape rates on each state to other states. This can be formulated as

$$\overline{\overline{Q}}_A = \begin{bmatrix} -\sum_{j=1}^{M_A-1}\lambda_{A,j} & \lambda_{A,1} & \cdots & \lambda_{A,M_A-1} \\ \lambda_{A,M_A} & -\sum_{j=M_A}^{2*M_A-2}\lambda_{A,j} & \cdots & \lambda_{A,2*M_A-2} \\ \vdots & \vdots & \ddots & \vdots \\ \lambda_{A,(M_A-1)^2+1} & \lambda_{A,(M_A-1)^2+2} & \cdots & -\sum_{j=(M_A-1)^2+1}^{M_A*(M_A-1)}\lambda_{A,j} \end{bmatrix} = \begin{bmatrix} \Lambda_{A,1} & 0 & \cdots & 0 \\ 0 & \Lambda_{A,2} & \cdots & 0 \\ \vdots & \vdots & \ddots & \vdots \\ 0 & 0 & \cdots & \Lambda_{A,M_A} \end{bmatrix} \begin{bmatrix} -1 & \pi^A_{1,1} & \cdots & \pi^A_{1,M_A} \\ \pi^A_{2,1} & -1 & \cdots & \pi^A_{2,M_A} \\ \vdots & \vdots & \ddots & \vdots \\ \pi^A_{M_A,1} & \pi^A_{M_A,2} & \cdots & -1 \end{bmatrix}. \tag{S77}$$

So, in this case, instead of having a prior for each transition rate, we have priors on escape rate and the ratio of escape to other states

$$\Lambda_{A,m_A} \sim \mathbf{Gamma}\left(\alpha_{\lambda,A}, \beta_{\lambda,A}\right), \qquad m_A = 1, \ldots, M_A \tag{S78}$$
$$\overline{\pi}^A_{m_A} \sim \mathbf{Dir}\left(\alpha_{\pi,A}, \overline{\beta}_{\pi,A}\right), \qquad m_A = 1, \ldots, M_A \tag{S79}$$
$$\overline{\beta}_{\pi,A} = \left[\frac{1}{M_A-1}, \ldots, \frac{1}{M_A-1}\right]. \tag{S80}$$

Also, the same logic applies to the donor-PIFE and FRET state transition rate matrices where we have

$$\overline{\overline{Q}}_D = \begin{bmatrix} -\sum_{j=1}^{M_D-1}\lambda_{D,j} & \lambda_{D,1} & \cdots & \lambda_{D,M_D-1} \\ \lambda_{D,M_D} & -\sum_{j=M_D}^{2*M_D-2}\lambda_{D,j} & \cdots & \lambda_{D,2*M_D-2} \\ \vdots & \vdots & \ddots & \vdots \\ \lambda_{D,(M_D-1)^2+1} & \lambda_{D,(M_D-1)^2+2} & \cdots & -\sum_{j=(M_D-1)^2+1}^{M_D*(M_D-1)}\lambda_{D,j} \end{bmatrix} = \begin{bmatrix} \Lambda_{D,1} & 0 & \cdots & 0 \\ 0 & \Lambda_{D,2} & \cdots & 0 \\ \vdots & \vdots & \ddots & \vdots \\ 0 & 0 & \cdots & \Lambda_{D,M_D} \end{bmatrix} \begin{bmatrix} -1 & \pi^D_{1,1} & \cdots & \pi^D_{1,M_D} \\ \pi^D_{2,1} & -1 & \cdots & \pi^D_{2,M_D} \\ \vdots & \vdots & \ddots & \vdots \\ \pi^D_{M_D,1} & \pi^D_{M_D,2} & \cdots & -1 \end{bmatrix}. \tag{S81}$$

So, in this case, instead of having a prior for each transition rate, we have priors on escape rate and the ratio of escape to other states

$$\Lambda_{D,m_D} \sim \mathbf{Gamma}\left(\alpha_{\lambda,D}, \beta_{\lambda,D}\right), \qquad m_D = 1, \ldots, M_D \tag{S82}$$
$$\overline{\pi}^D_{m_D} \sim \mathbf{Dir}\left(\alpha_{\pi,D}, \overline{\beta}_{\pi,D}\right), \qquad m_D = 1, \ldots, M_D \tag{S83}$$
$$\overline{\beta}_{\pi,D} = \left[\frac{1}{M_D-1}, \ldots, \frac{1}{M_D-1}\right]. \tag{S84}$$

$$\overline{\overline{Q}}_F = \begin{bmatrix} -\sum_{j=1}^{M_F-1}\lambda_{F,j} & \lambda_{F,1} & \cdots & \lambda_{F,M_F-1} \\ \lambda_{F,M_F} & -\sum_{j=M_F}^{2*M_F-2}\lambda_{F,j} & \cdots & \lambda_{F,2*M_F-2} \\ \vdots & \vdots & \ddots & \vdots \\ \lambda_{F,(M_F-1)^2+1} & \lambda_{F,(M_F-1)^2+2} & \cdots & -\sum_{j=(M_F-1)^2+1}^{M_F*(M_F-1)}\lambda_{F,j} \end{bmatrix} = \begin{bmatrix} \Lambda_{F,1} & 0 & \dots & 0 \\ 0 & \Lambda_{F,2} & \dots & 0 \\ \vdots & \vdots & \ddots & \vdots \\ 0 & 0 & \dots & \Lambda_{F,M_F} \end{bmatrix} \begin{bmatrix} -1 & \pi^F_{1,1} & \cdots & \pi^F_{1,M_F} \\ \pi^F_{2,1} & -1 & \cdots & \pi^A_{2,M_F} \\ \vdots & \vdots & \ddots & \vdots \\ \pi^F_{M_F,1} & \pi^F_{M_F,2} & \cdots & -1 \end{bmatrix}. \tag{S85}$$

So, in this case, instead of having a prior for each transition rate, we have priors on escape rate and the ratio of escape to other states

$$\Lambda_{F,m_F} \sim \textbf{Gamma}\left(\alpha_{\lambda,F}, \beta_{\lambda,F}\right), \qquad m_F = 1, \dots, M_F \tag{S86}$$

$$\overline{\pi}^F_{m_F} \sim \textbf{Dir}\left(\alpha_{\pi,F}, \overline{\beta}_{\pi,F}\right), \qquad m_F = 1, \dots, M_F \tag{S87}$$

$$\overline{\beta}_{\pi,F} = \left[\frac{1}{M_F-1}, \dots, \frac{1}{M_F-1}\right]. \tag{S88}$$

#### S4.2. Summary of model equations

For concreteness, below we summarize the entire set of equations used in our framework, including a complete list of priors and hyperpriors

$$\mu_{ex,A,m_A} \sim \textbf{Gamma}\left(\alpha_{\mu_A}, \beta_{\mu_A}\right), \qquad m_A = 1, \dots, M_A \tag{S89}$$

$$\mu_{ex,D,m_D} \sim \textbf{Gamma}\left(\alpha_{\mu_D}, \beta_{\mu_D}\right), \qquad m_D = 1, \dots, M_D \tag{S90}$$

$$\mu_{\text{back},A} \sim \textbf{Gamma}\left(\alpha_{\text{back},A}, \beta_{\text{back},A}\right) \tag{S91}$$

$$\mu_{\text{back},D} \sim \textbf{Gamma}\left(\alpha_{\text{back},D}, \beta_{\text{back},D}\right) \tag{S92}$$

$$\overline{\zeta}_A \sim \textbf{Dir}\left(\alpha_{A,\zeta}, \overline{\beta}_{A,hyper}\right) \tag{S93}$$

$$\overline{\zeta}_D \sim \textbf{Dir}\left(\alpha_{A,\zeta}, \overline{\beta}_{D,hyper}\right) \tag{S94}$$

$$\overline{\zeta}_F \sim \textbf{Dir}\left(\alpha_{A,\zeta}, \overline{\beta}_{F,hyper}\right) \tag{S95}$$

$$\overline{\beta}_{A,hyper} = \left[\frac{1}{M_A-1}, \dots, \frac{1}{M_A-1}\right] \tag{S96}$$

$$\overline{\beta}_{D,hyper} = \left[\frac{1}{M_D-1}, \dots, \frac{1}{M_D-1}\right] \tag{S97}$$

$$\overline{\beta}_{F,hyper} = \left[\frac{1}{M_F-1}, \dots, \frac{1}{M_F-1}\right] \tag{S98}$$

$$S_{A,1}|\overline{\zeta}_A \sim \textbf{Cat}_{1,\dots,M_A}\left(\overline{\beta}_A\right) \tag{S99}$$

$$S_{D,1}|\overline{\zeta}_D \sim \textbf{Cat}_{1,\dots,M_D}\left(\overline{\beta}_D\right) \tag{S100}$$

$$S_{F,1}|\overline{\zeta}_F \sim \textbf{Cat}_{1,\dots,M_F}\left(\overline{\beta}_F\right) \tag{S101}$$

$$S_{A,t+1}|S_{A,t}, \overline{\overline{Q}}_A, \{\overline{\pi}_{A,m_A}\}_{m_A}, \Delta_t \sim \textbf{Cat}_{1,\dots,M_A}\left(\overline{\overline{P}}_{A,t}\left(S_{A,t},:\right)\right), \qquad t = 1, \dots, T-1 \tag{S102}$$

$$S_{D,t+1}|S_{D,t}, \overline{\overline{Q}}_D, \{\overline{\pi}_{D,m_D}\}_{m_D}, \Delta_t \sim \textbf{Cat}_{1,\dots,M_D}\left(\overline{\overline{P}}_{D,t}\left(S_{D,t},:\right)\right), \qquad t = 1, \dots, T-1 \tag{S103}$$

$$S_{F,t+1}|S_{F,t}, \overline{\overline{Q}}_F, \{\overline{\pi}_{F,m_F}\}_{m_F}, \Delta_t \sim \textbf{Cat}_{1,\dots,M_F}\left(\overline{\overline{P}}_{F,t}\left(S_{F,t},:\right)\right), \qquad t = 1, \dots, T-1 \tag{S104}$$

$$\overline{\overline{P}}_{A,t} = \exp\left(\overline{\overline{Q}}_A \Delta_t\right) \tag{S105}$$

$$\overline{\overline{P}}_{D,t} = \exp\left(\overline{\overline{Q}}_D \Delta_t\right) \tag{S106}$$

$$\overline{\overline{P}}_{F,t} = \exp\left(\overline{\overline{Q}}_F \Delta_t\right) \tag{S107}$$

$$\overline{\overline{Q}}_A = \begin{bmatrix} -\sum_{j=1}^{M_A-1}\lambda_{A,j} & \lambda_{A,1} & \cdots & \lambda_{A,M_A-1} \\ \lambda_{A,M_A} & -\sum_{j=M_A}^{2*M_A-2}\lambda_{A,j} & \cdots & \lambda_{A,2*M_A-2} \\ \vdots & \vdots & \ddots & \vdots \\ \lambda_{A,(M_A-1)^2+1} & \lambda_{A,(M_A-1)^2+2} & \cdots & -\sum_{j=(M_A-1)^2+1}^{M_A*(M_A-1)}\lambda_{A,j} \end{bmatrix} = \begin{bmatrix} \Lambda_{A,1} & 0 & \cdots & 0 \\ 0 & \Lambda_{A,2} & \cdots & 0 \\ \vdots & \vdots & \ddots & \vdots \\ 0 & 0 & \cdots & \Lambda_{A,M_A} \end{bmatrix} \begin{bmatrix} -1 & \pi^A_{1,1} & \cdots & \pi^A_{1,M_A} \\ \pi^A_{2,1} & -1 & \cdots & \pi^A_{2,M_A} \\ \vdots & \vdots & \ddots & \vdots \\ \pi^A_{M_A,1} & \pi^A_{M_A,2} & \cdots & -1 \end{bmatrix} \tag{S108}$$

$$\overline{\overline{Q}}_D = \begin{bmatrix} -\sum_{j=1}^{M_D-1}\lambda_{D,j} & \lambda_{D,1} & \cdots & \lambda_{D,M_D-1} \\ \lambda_{D,M_D} & -\sum_{j=M_D}^{2*M_D-2}\lambda_{D,j} & \cdots & \lambda_{D,2*M_D-2} \\ \vdots & \vdots & \ddots & \vdots \\ \lambda_{D,(M_D-1)^2+1} & \lambda_{D,(M_D-1)^2+2} & \cdots & -\sum_{j=(M_D-1)^2+1}^{M_D*(M_D-1)}\lambda_{D,j} \end{bmatrix} = \begin{bmatrix} \Lambda_{D,1} & 0 & \cdots & 0 \\ 0 & \Lambda_{D,2} & \cdots & 0 \\ \vdots & \vdots & \ddots & \vdots \\ 0 & 0 & \cdots & \Lambda_{D,M_D} \end{bmatrix} \begin{bmatrix} -1 & \pi^D_{1,1} & \cdots & \pi^D_{1,M_D} \\ \pi^D_{2,1} & -1 & \cdots & \pi^A_{2,M_D} \\ \vdots & \vdots & \ddots & \vdots \\ \pi^D_{M_D,1} & \pi^D_{M_D,2} & \cdots & -1 \end{bmatrix} \tag{S109}$$

$$\overline{\overline{Q}}_F = \begin{bmatrix} -\sum_{j=1}^{M_F-1}\lambda_{F,j} & \lambda_{F,1} & \cdots & \lambda_{F,M_F-1} \\ \lambda_{F,M_F} & -\sum_{j=M_F}^{2*M_F-2}\lambda_{F,j} & \cdots & \lambda_{F,2*M_F-2} \\ \vdots & \vdots & \ddots & \vdots \\ \lambda_{F,(M_F-1)^2+1} & \lambda_{F,(M_F-1)^2+2} & \cdots & -\sum_{j=(M_F-1)^2+1}^{M_F*(M_F-1)}\lambda_{F,j} \end{bmatrix} = \begin{bmatrix} \Lambda_{F,1} & 0 & \cdots & 0 \\ 0 & \Lambda_{F,2} & \cdots & 0 \\ \vdots & \vdots & \ddots & \vdots \\ 0 & 0 & \cdots & \Lambda_{F,M_F} \end{bmatrix} \begin{bmatrix} -1 & \pi^F_{1,1} & \cdots & \pi^F_{1,M_F} \\ \pi^F_{2,1} & -1 & \cdots & \pi^F_{2,M_F} \\ \vdots & \vdots & \ddots & \vdots \\ \pi^F_{M_F,1} & \pi^F_{M_F,2} & \cdots & -1 \end{bmatrix} \tag{S110}$$

$$\tau_{A,m_A} \sim \textbf{Gamma}\left(\alpha_A, \beta_A\right), \qquad m_A = 1, \ldots, M_A \tag{S111}$$
$$\Lambda_{A,m_A} \sim \textbf{Gamma}\left(\alpha_{\lambda,A}, \beta_{\lambda,A}\right), \qquad m_A = 1, \ldots, M_A \tag{S112}$$
$$\overline{\pi}^A_{m_A} \sim \textbf{Dir}\left(\alpha_{\pi,A}, \overline{\beta}_{\pi,A}\right), \qquad m_A = 1, \ldots, M_A \tag{S113}$$
$$\overline{\beta}_{\pi,A} = \left[\frac{1}{M_A-1}, \ldots, \frac{1}{M_A-1}\right] \tag{S114}$$

$$\tau_{D,m_D} \sim \textbf{Gamma}\left(\alpha_D, \beta_D\right), \qquad m_D = 1, \ldots, M_D \tag{S115}$$
$$\Lambda_{D,m_D} \sim \textbf{Gamma}\left(\alpha_{\lambda,D}, \beta_{\lambda,D}\right), \qquad m_D = 1, \ldots, M_D \tag{S116}$$
$$\overline{\pi}^D_{m_D} \sim \textbf{Dir}\left(\alpha_{\pi,D}, \overline{\beta}_{\pi,D}\right), \qquad m_D = 1, \ldots, M_D \tag{S117}$$
$$\overline{\beta}_{\pi,D} = \left[\frac{1}{M_D-1}, \ldots, \frac{1}{M_D-1}\right] \tag{S118}$$

$$k_{F,m_F} \sim \textbf{Gamma}\left(\alpha_F, \beta_F\right), \qquad m_F = 1, \ldots, M_F \tag{S119}$$
$$\Lambda_{F,m_F} \sim \textbf{Gamma}\left(\alpha_{\lambda,F}, \beta_{\lambda,F}\right), \qquad m_F = 1, \ldots, M_F \tag{S120}$$
$$\overline{\pi}^F_{m_F} \sim \textbf{Dir}\left(\alpha_{\pi,F}, \overline{\beta}_{\pi,F}\right), \qquad m_F = 1, \ldots, M_F \tag{S121}$$
$$\overline{\beta}_{\pi,F} = \left[\frac{1}{M_F-1}, \ldots, \frac{1}{M_F-1}\right] \tag{S122}$$

$$\eta_A \sim \textbf{Beta}\left(\alpha_{\eta,A}, \beta_{\eta,A}\right) \tag{S123}$$
$$\eta_D \sim \textbf{Beta}\left(\alpha_{\eta,D}, \beta_{\eta,D}\right) \tag{S124}$$

$$\pi'_{D,t} = \pi_{D,t}\left(1-\pi_{A,t}\right)\left(1-\pi_{BD}\right)\left(1-\pi_{BA}\right) \tag{S125}$$
$$\pi'_{A,t} = \pi_{A,t}\left(1-\pi_{D,t}\right)\left(1-\pi_{BD}\right)\left(1-\pi_{BA}\right) \tag{S126}$$
$$\pi'_{BD,t} = \pi_{BD}\left(1-\pi_{D,t}\right)\left(1-\pi_{A,t}\right)\left(1-\pi_{BA}\right) \tag{S127}$$
$$\pi'_{BA,t} = \pi_{BA}\left(1-\pi_{D,t}\right)\left(1-\pi_{A,t}\right)\left(1-\pi_{BD}\right) \tag{S128}$$

$$\pi_{D,t} = 1-\exp\left(-\delta_p \mu_{ex,D,S_{D,t}}\right) \tag{S129}$$
$$\pi_{A,t} = 1-\exp\left(-\delta_p \mu_{ex,A,S_{A,t}}\right) \tag{S130}$$
$$\pi_{BD} = 1-\exp\left(-T_p \mu_{\text{back},D}\right) \tag{S131}$$
$$\pi_{BA} = 1-\exp\left(-T_p \mu_{\text{back},A}\right) \tag{S132}$$

$$C_t|\eta_A,\eta_D,\mu_{\text{back},A},\mu_{\text{back},D},\{\mu_{ex,A,m_A}\}_{m_A},\{\tau_{D,m_D},\mu_{ex,D,m_D}\}_{m_D},\{k_{F,m_F}\}_{m_F},S_{A,t},S_{D,t},S_{F,t} \sim$$
$$\mathbf{Cat}_{1,2}\left(\left[\pi'_{D,t}\left(\frac{\frac{1}{\tau_{D,S_{D,t}}}}{K_t}\right)+\pi'_{BD,t}\ ,\ \pi'_{D,t}\left(\frac{k_{F,S_{F,t}}}{K_t}\right)+\pi'_{A,t}+\pi'_{BA,t}\right]\begin{bmatrix}1-\eta_D & \eta_D\\ \eta_A & 1-\eta_A\end{bmatrix}\right) \tag{S133}$$
$$t=1,\ldots,T \tag{S134}$$

$$\delta_t|C_t,\eta_A,\eta_D,\mu_{\text{back},A},\mu_{\text{back},D},\{\tau_{A,m_A},\mu_{ex,A,m_A}\}_{m_A},\{\tau_{D,m_D},\mu_{ex,D,m_D}\}_{m_D},\{k_{F,m_F}\}_{m_F},S_{A,t},S_{D,t},S_{F,t} \sim$$
$$\begin{cases}\dfrac{\left[\pi'_{D,t}f^{D}_{D^*D,t}+\pi'_{BD,t}P_B(.)\right](1-\eta_D)+\left[\pi'_{D,t}f^{D}_{D^*A,t}+\pi'_{A,t}f^{D}_{A^*A,t}+\pi'_{BA,t}P_B(.)\right]\eta_A}{\left(\pi'_{D,t}\frac{\frac{1}{\tau_{D,S_{D,t}}}}{K_t}+\pi'_{BD,t}\right)(1-\eta_D)+\left(\pi'_{D,t}\frac{k_{F,S_{F,t}}}{K_t}+\pi'_{A,t}+\pi'_{BA,t}\right)\eta_A} & C_t=1 \text{ (Donor channel)}\\[2ex] \dfrac{\left[\pi'_{D,t}f^{A}_{D^*D,t}+\pi'_{BD,t}P_B(.)\right]\eta_D+\left[\pi'_{D,t}f^{A}_{D^*A,t}+\pi'_{A,t}f^{A}_{A^*A,t}(.)+\pi'_{BA,t}P_B(.)\right](1-\eta_A)}{\left(\pi'_{D,t}\frac{\frac{1}{\tau_{D,S_{D,t}}}}{K_t}+\pi'_{BD,t}\right)\eta_D+\left(\pi'_{D,t}\frac{k_{F,S_{F,t}}}{K_t}+\pi'_{A}+\pi'_{BA,t}\right)(1-\eta_A)} & C_t=2 \text{ (Acceptor channel)}\end{cases} \tag{S135}$$
$$t=1,\ldots,T \tag{S136}$$

$$f^{D}_{D^{*}D,t} = \frac{1}{2\tau_{D,S_{D,t}}} \exp\left(K_t\left(\mu_{D,t} + \frac{\sigma^2_{\text{IRF},D}}{2}K_t\right)\right) erfc\left(\frac{\mu_{D,t} + \sigma^2_{\text{IRF},D}K_t}{\sigma_{\text{IRF},D}\sqrt{2}}\right) \tag{S137}$$

$$f^{D}_{D^{*}A,t} = \frac{K_{F,S_{F,t}}}{2\left(K_t\tau_{A,S_{A,t}} - 1\right)} \left[\begin{array}{l} \exp\left(\frac{\sigma^2_{\text{IRF},D} + 2\tau_{A,S_{A,t}}\mu_{D,t}}{2\left(\tau_{A,S_{A,t}}\right)^2}\right) erfc\left(\frac{\frac{\sigma^2_{\text{IRF},D}}{\tau_{A,S_{A,t}}} + \mu_{D,t}}{\sigma_{\text{IRF},D}\sqrt{2}}\right) \\ - \exp\left(K_t\left(\mu_{D,t} + \frac{\sigma^2_{\text{IRF},D}}{2}K_t\right)\right) erfc\left(\frac{\sigma^2_{\text{IRF},D}K_t + \mu_{D,t}}{\sigma_{\text{IRF},D}\sqrt{2}}\right) \end{array}\right] \tag{S138}$$

$$f^{A}_{D^{*}A,t} = \frac{K_{F,S_{F,t}}}{2\left(K_t\tau_{A,S_{A,t}} - 1\right)} \left[\begin{array}{l} \exp\left(\frac{\sigma^2_{\text{IRF},A} + 2\tau_{A,S_{A,t}}\mu_{A,t}}{2\left(\tau_{A,S_{A,t}}\right)^2}\right) erfc\left(\frac{\frac{\sigma^2_{\text{IRF},A}}{\tau_{A,S_{A,t}}} + \mu_{A,t}}{\sigma_{\text{IRF},A}\sqrt{2}}\right) \\ - \exp\left(K_t\left(\mu_{A,t} + \frac{\sigma^2_{\text{IRF},A}}{2}K_t\right)\right) erfc\left(\frac{\sigma^2_{\text{IRF},A}K_t + \mu_{A,t}}{\sigma_{\text{IRF},A}\sqrt{2}}\right) \end{array}\right] \tag{S139}$$

$$f^{A}_{D^{*}D,t} = \frac{1}{2\tau_{D,S_{D,t}}} \exp\left(K_t\left(\mu_{A,t} + \frac{\sigma^2_{\text{IRF},A}}{2}K_t\right)\right) erfc\left(\frac{\mu_{A,t} + \sigma^2_{\text{IRF},A}K_t}{\sigma_{\text{IRF},A}\sqrt{2}}\right) \tag{S140}$$

$$P^{A}_{A^{*}A,t} = \frac{1}{2\tau_{A,S_{A,t}}} \exp\left(\frac{\mu_{A,t} + \frac{\sigma^2_{\text{IRF},A}}{2\tau_{A,S_{A,t}}}}{\tau_{A,S_{A,t}}}\right) erfc\left(\frac{\mu_{A,t} + \frac{\sigma^2_{\text{IRF},A}}{\tau_{A,S_{A,t}}}}{\sigma_{\text{IRF},A}\sqrt{2}}\right) \tag{S141}$$

$$P^{D}_{A^{*}A,t}(.) = \frac{1}{2\tau_{A,S_{A,t}}} \exp\left(\frac{\mu_{D,t} + \frac{\sigma^2_{\text{IRF},D}}{2\tau_{A,S_{A,t}}}}{\tau_{A,S_{A,t}}}\right) erfc\left(\frac{\mu_{D,t} + \frac{\sigma^2_{\text{IRF},D}}{\tau_{A,S_{A,t}}}}{\sigma_{\text{IRF},D}\sqrt{2}}\right) \tag{S142}$$

$$P_B(.) = \textbf{Uniform}\left([0, T_p]\right) \tag{S143}$$

where, $\mu_{D,t} = \mu_{\text{IRF},D} - \delta_t$, $\mu_{A,t} = \mu_{\text{IRF},A} - \delta_t$, and $K_t = k_{S_{F,t}} + \frac{1}{\tau_{D,S_{D,t}}}$.

### S5. Description of the computational scheme

The joint probability distribution of our framework is $(\eta_A, \eta_D, \mu_{\text{back},A}, \mu_{\text{back},D}, \{\tau_{A,m_A}, \lambda_{A,m_A}, \overline{\pi}_{A,m_A}, \mu_{ex,A,m_A}\}_{m_A}, \{\tau_{D,m_D}, \lambda_{D,m_D}, \overline{\pi}_{D,m_D}, \mu_{ex,D,m_D}\}_{m_D}, \{k_{F,m_F}, \lambda_{F,m_F}, \overline{\pi}_{F,m_F}\}_{m_F}, \overline{S}_A, \overline{S}_D, \overline{S}_F, \overline{\beta}_A, \overline{\beta}_D, \overline{\beta}_F | \overline{\delta}, \overline{C})$, where the micro-time and detector trajectories (measurements) are gathered in

$$\overline{C} = C_1, C_2, \ldots, C_T \tag{S144}$$

$$\overline{\delta} = \delta_1, \delta_2, \ldots, \delta_T \tag{S145}$$

and the inter-arrival time of the photons, (*e.g.* $\Delta_1 = t_2 - t_1$) are

$$\overline{\Delta} = \Delta_1, \Delta_2, \ldots, \Delta_{T-1}. \tag{S146}$$

Since we only rely on the detected photons based on the pulse excitation, we do not consider the inter-arrival times as part of the likelihood, and only consider them as the experimental timestamps.

So, based on this framework, we develop a specialized Markov chain Monte Carlo (MCMC) scheme that can be used to generate pseudo-random samples [1–6]. This scheme is explained in detail below.

To ease the usage of the algorithm we design an app based on the Matlab app designer shown in Fig. S5. This working implementation complimentary to the source code will allow experts and non-expert in Matlab coders, to use the algorithm without any complications.

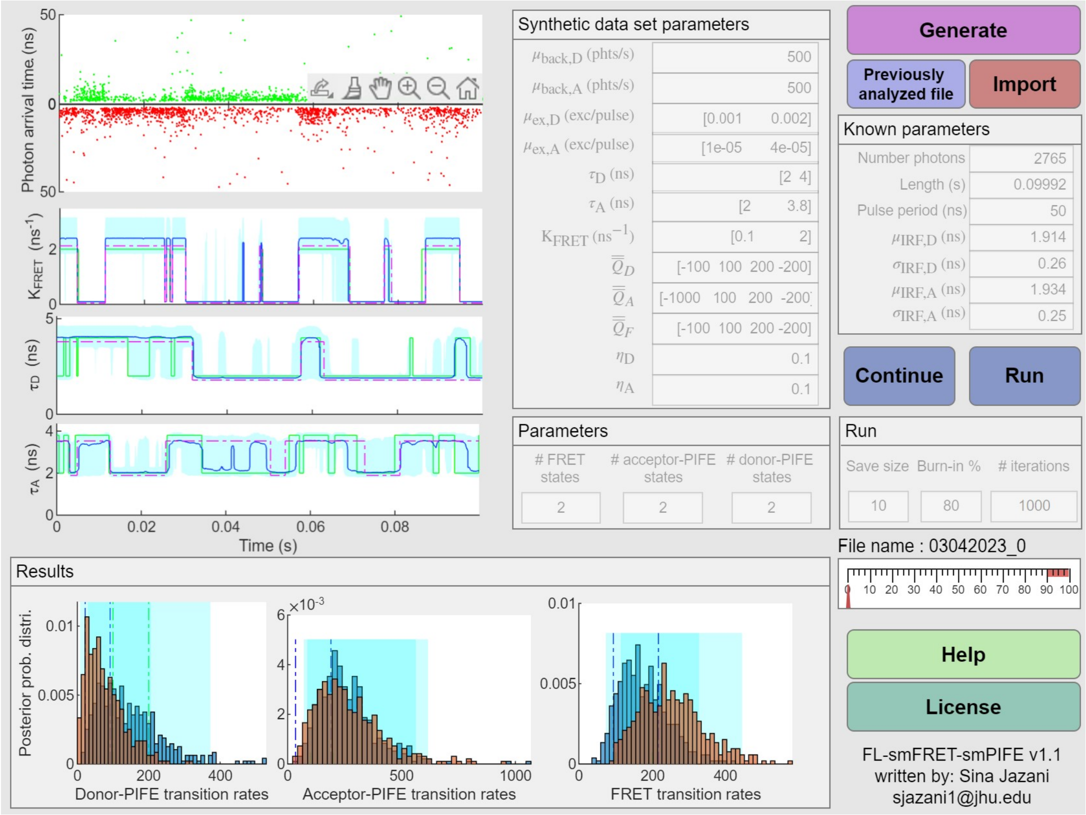


FIG. S5. **The installable application in Matlab as the working implementation of the described algorithm in this study.** This app, along with the source code can be used to analyze single-photon time traces.

#### S5.1. Overview of the sampling updates

Our MCMC utilizes a Gibbs sampling scheme [1, 2, 5]. Accordingly, posterior samples are generated by updating each one of the variables involved sequentially by sampling conditioned on all other variables and measurements $\overline{\delta}$ and $\overline{C}$. Conceptually, the steps involved in the generation of each posterior sample ($\eta_A, \eta_D, \mu_{\text{back},A}, \mu_{\text{back},D}$, $\{\tau_{A,m_A}, \mu_{ex,A,m_A}\}_{m_A}, \{\tau_{D,m_D}, \mu_{ex,D,m_D}\}_{m_D}, \{k_{F,m_F}\}_{m_F}, \overline{\overline{Q}}_A, \overline{\overline{Q}}_D, \overline{\overline{Q}}_F, \overline{S}_A, \overline{S}_D, \overline{S}_F, \overline{\zeta}_A, \overline{\zeta}_D, \overline{\zeta}_F | \overline{\delta}, \overline{C}$) are:

(1) Update the state trajectory of acceptor lifetime (acceptor-PIFE) $\overline{S}_A$

(2) Update the state trajectory of donor lifetime (donor-PIFE) $\overline{S}_D$

(3) Update the state trajectory of energy transfer rates (FRET) $\overline{S}_F$

(4) Update the transition rate matrix of acceptor lifetime trajectory $\overline{\overline{Q}}_A$

(5) Update the transition rate matrix of donor lifetime trajectory $\overline{\overline{Q}}_D$

(6) Update the transition rate matrix of FRET trajectory $\overline{\overline{Q}}_F$

(7) Update the weight on initial state of acceptor PIFE $\overline{\zeta}_A$

(8) Update the weight on initial state of donor PIFE $\overline{\zeta}_D$

(9) Update the weight on initial state of FRET $\overline{\zeta}_F$

(10) Jointly update the acceptor lifetimes of all states $\{\tau_{A,m_A}\}_{m_A}$, donor lifetimes of all states $\{\tau_{D,m_D}\}_{m_D}$, and resonance energy transfer rates of all states $\{k_{F,m_F}\}_{m_F}$

(11) Jointly update the cross-talk ratios of acceptor and donor channels, $\eta_A$ and $\eta_D$

(12) Jointly update the acceptor excitation rates $\{\mu_{ex,A,m_A}\}_{m_A}$, donor excitation rates $\{\mu_{ex,D,m_D}\}_{m_D}$, acceptor channel background photon emission rate $\mu_{\text{back},A}$, and donor channel background photon emission rate $\mu_{\text{back},D}$.

These steps are described in detail below for each individual step. It is worth mentioning that for a better convergence, we can have a mixture of steps as well.

#### S5.2. Update the state trajectory of acceptor lifetimes $\overline{S}_A$

First, we sample the acceptor lifetime state trajectory $\overline{S}_A$ by applying forward filtering and backward sampling (FFBS) scheme [7–9] on the join conditional

$$\begin{aligned}&\overline{S}_A|\eta_A, \eta_D, \mu_{\text{back},A}, \mu_{\text{back},D}, \{\tau_{A,m_A}, \mu_{ex,A,m_A}\}_{m_A}, \{\tau_{D,m_D}, \mu_{ex,D,m_D}\}_{m_D},\\ &\quad \{k_{F,m_F}\}_{m_F}, \overline{\overline{Q}}_A, \overline{\overline{Q}}_D, \overline{\overline{Q}}_F, \overline{S}_D, \overline{S}_F, \overline{\zeta}_A, \overline{\zeta}_D, \overline{\zeta}_F, \overline{\delta}, \overline{C}\end{aligned} \tag{S147}$$

which can be simplify to

$$\overline{S}_A|\eta_A, \eta_D, \mu_{\text{back},A}, \mu_{\text{back},D}, \{\tau_{A,m_A}, \mu_{ex,A,m_A}\}_{m_A}, \{\tau_{D,m_D}, \mu_{ex,D,m_D}\}_{m_D}, \{k_{F,m_F}\}_{m_F}, \overline{\overline{Q}}_A, \overline{S}_D, \overline{S}_F, \overline{\zeta}_F, \overline{\delta}, \overline{C}. \tag{S148}$$

Here, we consider Markovian dynamics on the transition of the acceptor lifetime states with transition probability matrix

$$\overline{\overline{P}}_{A,t} = \exp\left(\overline{\overline{Q}}_A \times \Delta_t\right). \tag{S149}$$

So, to apply the FFBS, first, we need to calculate the filter at each time point. This step is known as the forward filtering. In this step we read the signal from the beginning and trying to filter the possible states. It means that we check the possibility of each event based on the previous one and also compare the probability of it based on the observation. Starting from the initial time which has a dependency on the weights of the states $\overline{\zeta}_A$ as the prior

$$\alpha_1(S_{A,1}) = p(C_1|S_{A,1},\dots)\, p(\delta_1|S_{A,1}, C_1\dots)\, p\left(S_{A,1}|\overline{\zeta}_A\right). \tag{S150}$$

Then, we are marching forward by marginalizing over previous states

$$\alpha_t(S_{A,t}) = p(C_t|S_{A,t},\dots)\, p(\delta_t|S_{A,t}, C_t\dots) \sum_{S_{A,t-1}=1}^{M_A} \overline{\overline{P}}_{A,t}(S_{A,t}|S_{A,t-1},\dots)\, \alpha_{t-1}(S_{A,t-1}), \qquad t = 2,\dots,T. \tag{S151}$$

Now that we compute the filter densities $\alpha_t(S_{A,t})$ in the forward filtering, we are able to sample the states $S_{A,t}$ by using backward sampling. Specifically, given a computed filter, we sample sequentially $S_{A,t}$ according to

$$S_{A,T} \sim \alpha_T(S_{A,T}) \tag{S152}$$

$$S_{A,t} \sim \alpha_t(S_{A,t})\, \overline{\overline{P}}_{A,t}(S_{A,t+1}|S_{A,t}), \qquad t = T-1,\dots,1. \tag{S153}$$

#### S5.3. Update the state trajectory of donor lifetimes $\overline{S}_D$

Next, we sample the donor lifetime state trajectory $\overline{S}_D$ by applying forward filtering and backward sampling (FFBS) scheme [7–9] on the join conditional

$$\begin{aligned}&\overline{S}_D|\eta_A, \eta_D, \mu_{\text{back},A}, \mu_{\text{back},D}, \{\tau_{A,m_A}, \mu_{ex,A,m_A}\}_{m_A}, \{\tau_{D,m_D}, \mu_{ex,D,m_D}\}_{m_D},\\ &\quad \{k_{F,m_F}\}_{m_F}, \overline{\overline{Q}}_A, \overline{\overline{Q}}_D, \overline{\overline{Q}}_F, \overline{S}_A, \overline{S}_F, \overline{\zeta}_A, \overline{\zeta}_D, \overline{\zeta}_F, \overline{\delta}, \overline{C}\end{aligned} \tag{S154}$$

which can be simplify to

$$\overline{S}_D|\eta_A, \eta_D, \mu_{\text{back},A}, \mu_{\text{back},D}, \{\tau_{A,m_A}, \mu_{ex,A,m_A}\}_{m_A}, \{\tau_{D,m_D}, \mu_{ex,D,m_D}\}_{m_D}, \{k_{F,m_F}\}_{m_F}, \overline{\overline{Q}}_D, \overline{S}_A, \overline{\zeta}_F, \overline{\delta}, \overline{C}. \tag{S155}$$

Here, we consider Markovian dynamics on the transition of the donor lifetime states with transition probability matrix

$$\overline{\overline{P}}_{D,t} = \exp\left(\overline{\overline{Q}}_D \times \Delta_t\right). \tag{S156}$$

So, to apply the FFBS, first, we need to calculate the filter at each time point. This step is known as the forward filtering. In this step we read the signal from the beginning and trying to filter the possible states. It means that we check the possibility of each event based on the previous one and also compare the probability of it based on the observation. Starting from the initial time which has a dependency on the weights of the states $overlinezeta_D$ as the prior

$$\alpha_1 \left(S_{D,1}\right) = p\left(C_1 | S_{D,1}, \dots\right) p\left(\delta_1 | S_{D,1}, C_1 \dots\right) p\left(S_{D,1} | \overline{\zeta}_D\right). \tag{S157}$$

Then, we are marching forward by marginalizing over previous states

$$\alpha_t \left(S_{D,t}\right) = p\left(C_t | S_{D,t}, \dots\right) p\left(\delta_t | S_{D,t}, C_t \dots\right) \sum_{S_{D,t-1}=1}^{M_D} \overline{\overline{P}}_{D,t}\left(S_{D,t} | S_{D,t-1}, \dots\right) \alpha_{t-1}\left(S_{D,t-1}\right), \qquad t = 2, \dots, T. \tag{S158}$$

Now that we compute the filter densities $\alpha_t\left(S_{D,t}\right)$ in the forward filtering, we are able to sample the states $S_{D,t}$ by using backward sampling. Specifically, given a computed filter, we sample sequentially $S_{D,t}$ according to

$$S_{D,T} \sim \alpha_T\left(S_{D,T}\right) \tag{S159}$$

$$S_{D,t} \sim \alpha_t\left(S_{D,t}\right) \overline{\overline{P}}_{D,t}\left(S_{D,t+1} | S_{D,t}\right), \qquad t = T-1, \dots, 1. \tag{S160}$$

#### S5.4. Update the state trajectory of energy transfer rates $\overline{S}_F$

Next, we sample the state trajectory of energy transfer rates $\overline{S}_F$ by applying forward filtering and backward sampling (FFBS) scheme [7–9] on the join conditional

$$\begin{aligned} &\overline{S}_F | \eta_A, \eta_D, \mu_{\text{back},A}, \mu_{\text{back},D}, \{\tau_{A,m_A}, \mu_{ex,A,m_A}\}_{m_A}, \{\tau_{D,m_D}, \mu_{ex,D,m_D}\}_{m_D}, \\ &\{k_{F,m_F}\}_{m_F}, \overline{\overline{Q}}_A, \overline{\overline{Q}}_D, \overline{\overline{Q}}_F, \overline{S}_A, \overline{S}_D, \overline{\beta}_A, \overline{\beta}_D, \overline{\beta}_F, \overline{\delta}, \overline{C} \end{aligned} \tag{S161}$$

which can be simplified to

$$\overline{S}_F | \eta_A, \eta_D, \mu_{\text{back},A}, \mu_{\text{back},D}, \{\tau_{A,m_A}, \mu_{ex,A,m_A}\}_{m_A}, \{\tau_{D,m_D}, \mu_{ex,D,m_D}\}_{m_D}, \{k_{F,m_F}\}_{m_F}, \overline{\overline{Q}}_F, \overline{S}_A, \overline{S}_D, \overline{\zeta}_F, \overline{\delta}, \overline{C}. \tag{S162}$$

Here, we consider Markovian dynamics on the transition of the FRET states with transition probability matrix

$$\overline{\overline{P}}_{F,t} = \exp\left(\overline{\overline{Q}}_F \times \Delta_t\right). \tag{S163}$$

So, to apply the FFBS, first, we need to calculate the filter at each time point. This step is known as the forward filtering. In this step, we read the signal from the beginning and try to filter the possible states. It means that we check the possibility of each event based on the previous one and also compare the probability of it based on the observation. Starting from the initial time which has a dependency on the weights of the states $\overline{\zeta}_{Fi}$ as the prior

$$\alpha_1 \left(S_{F,1}\right) = p\left(C_1 | S_{F,1}, \dots\right) p\left(\delta_1 | S_{F,1}, C_1 \dots\right) p\left(S_{F,1} | \overline{\zeta}_F\right). \tag{S164}$$

Then, we are marching forward by marginalizing over previous states

$$\alpha_t \left(S_{F,t}\right) = p\left(C_t | S_{F,t}, \dots\right) p\left(\delta_t | S_{F,t}, C_t \dots\right) \sum_{S_{F,t-1}=1}^{M_F} \overline{\overline{P}}_{F,t}\left(S_{F,t} | S_{F,t-1}, \dots\right) \alpha_{t-1}\left(S_{F,t-1}\right), \qquad t = 2, \dots, T. \tag{S165}$$

Now that we compute the filter densities $\alpha_t\left(S_{F,t}\right)$ in the forward filtering, we are able to sample the states $S_{F,t}$ by using backward sampling. Specifically, given a computed filter, we sample sequentially $S_{F,t}$ according to

$$S_{F,T} \sim \alpha_T\left(S_{F,T}\right) \tag{S166}$$

$$S_{F,t} \sim \alpha_t\left(S_{F,t}\right) \overline{\overline{P}}_{F,t}\left(S_{F,t+1} | S_{F,t}\right), \qquad t = T-1, \dots, 1. \tag{S167}$$

#### S5.5. Update the acceptor-PIFE state transition matrix $\overline{\overline{Q}}_A$

After sampling the state trajectory of the acceptor lifetime we jointly sample all nondiagonal elements of the transition matrix $\overline{\overline{Q}}_A$ through the join conditional $p(\overline{\overline{Q}}_A|\eta_A, \eta_D, \mu_{\text{back},A}, \mu_{\text{back},D}, \{\tau_{A,m_A},\ \mu_{ex,A,m_A}\}_{m_A}, \{\tau_{D,m_D},\ \mu_{ex,D,m_D}\}_{m_D},\ \{k_{F,m_F}\}_{m_F}, \overline{\overline{Q}}_D, \overline{\overline{Q}}_F, \overline{S}_A, \overline{S}_D, \overline{S}_F,\ \overline{\zeta}_A,\ \overline{\zeta}_D,\ \overline{\zeta}_F,\ \overline{\delta}, \overline{C})$ which can be simplify to $p\left(\overline{\overline{Q}}_A|\overline{S}_A\right)$. Here, the $p\left(\overline{\overline{Q}}_A|\overline{S}_A\right)$ can be rewritten as

$$p\left(\overline{\overline{Q}}_A|\overline{S}_A\right) = p\left(\{\Lambda_{m_A}, \overline{\pi}^A_{m_A}\}_{m_A}|\overline{S}_A\right). \tag{S168}$$

So, for escape rates we have

$$\begin{aligned} p\left(\Lambda_{m'_A}|\overline{S}_A, \{\Lambda_{m_A}\}_{m_A\neq m'_A}, \{\overline{\pi}^A_{m_A}\}_{m_A}\right) &\propto p\left(\overline{S}_A|\{\Lambda_{m_A}, \overline{\pi}^A_{m_A}\}_{m_A}\right) p\left(\Lambda_{m'_A}\right) \\ &= \left[\prod_{k=1}^{K} \textbf{Exp}\left(\Delta_{A,k}; \Lambda_{m_A}\right)\right] \textbf{Gamma}\left(\Lambda_{m'_A}; \alpha_{\lambda,A}, \beta_{\lambda,A}\right), m'_A = 1,\ldots,M_A \end{aligned} \tag{S169}$$

where, $\Delta_{A,k}$ is the kth dwell time at state $m_A$, and $K$ is the total number of dwell times at state $m'_A$. So, we have

$$\Lambda_{m'_A} \sim \textbf{Gamma}\left(\alpha', \beta'\right), m'_A = 1,\ldots,M_A \tag{S170}$$

where, $\alpha' = \alpha_{\lambda,A} + \sum_{t=1}^{T-1} \mathbb{I}\left(S_{A,t} = m'_A\right) \mathbb{I}\left(S_{A,t} \neq m'_A\right)$ and $\beta' = \frac{1}{\frac{1}{\beta_{\lambda,A}} + \sum_{k=1}^{K} \Delta_{A,k}}$.

Then, for the weights we have

$$\begin{aligned} p\left(\overline{\pi}^A_{m'_A}|\overline{S}_A, \{\Lambda_{m_A}\}_{m_A}, \{\overline{\pi}^A_{m_A}\}_{m_A\neq m'_A}\right) &\propto p\left(\overline{S}_A|\{\Lambda_{m_A}, \overline{\pi}^A_{m_A}\}_{m_A}\right) p\left(\overline{\pi}^A_{m_A}\right) \\ &= \textbf{Cat}\left(\left[\sum_{t=1}^{T-1} \mathbb{I}\left(S_{A,t} = m'_A\right) \mathbb{I}\left(S_{A,t+1} = 1\right), \ldots, \sum_{t=1}^{T-1} \mathbb{I}\left(S_{A,t} = m'_A\right) \mathbb{I}\left(S_{A,t+1} = M_A\right)\right]\right) \\ &\times \textbf{Dir}\left(\overline{\pi}^A_{m'_A}; \alpha_{\pi,A}, \overline{\beta}_{\pi,A}\right), m'_A = 1,\ldots,M_A. \end{aligned} \tag{S171}$$

So, we have

$$\begin{aligned} \overline{\pi}^A_{m'_A} \sim \textbf{Dir}&\left(\left[\frac{\alpha_{\pi,A}}{M_A - 1} + \sum_{t=1}^{T-1} \mathbb{I}\left(S_{A,t} = m'_A\right) \mathbb{I}\left(S_{A,t+1} = 1\right), \ldots, \frac{\alpha_{\pi,A}}{M_A - 1} + \sum_{t=1}^{T-1} \mathbb{I}\left(S_{A,t} = m'_A\right) \mathbb{I}\left(S_{A,t+1} = M_A\right)\right]\right) \\ & m'_A = 1,\ldots,M_A. \end{aligned} \tag{S172}$$

#### S5.6. Update the transition matrix of donor lifetime trajectory $\overline{\overline{Q}}_D$

After sampling the state trajectory of the donor lifetime we jointly sample all nondiagonal elements of the transition matrix $\overline{\overline{Q}}_D$ through the join conditional $p(\overline{\overline{Q}}_D|\eta_A, \eta_D, \mu_{\text{back},A}, \mu_{\text{back},D}, \{\tau_{A,m_A},\ \mu_{ex,A,m_A}\}_{m_A}, \{\tau_{D,m_D},\ \mu_{ex,D,m_D}\}_{m_D}, \{k_{F,m_F}\}_{m_F}, \overline{\overline{Q}}_D, \overline{\overline{Q}}_F, \overline{S}_A, \overline{S}_D, \overline{S}_F,\ \overline{\zeta}_A,\ \overline{\zeta}_D,\ \overline{\zeta}_F,\ \overline{\delta}, \overline{C})$ which can be simplify to $p\left(\overline{\overline{Q}}_A|\overline{S}_A\right)$. Here, the $p\left(\overline{\overline{Q}}_D|\overline{S}_D\right)$ can be rewritten as

$$p\left(\overline{\overline{Q}}_D|\overline{S}_D\right) = p\left(\{\Lambda_{m_D}, \overline{\pi}^D_{m_D}\}_{m_D}|\overline{S}_D\right). \tag{S173}$$

So, for escape rates we have

$$\begin{aligned} p\left(\Lambda_{m'_D}|\overline{S}_D, \{\Lambda_{m_D}\}_{m_D\neq m'_D}, \{\overline{\pi}^D_{m_D}\}_{m_D}\right) &\propto p\left(\overline{S}_D|\{\Lambda_{m_D}, \overline{\pi}^D_{m_D}\}_{m_D}\right) p\left(\Lambda_{m'_D}\right) \\ &= \left[\prod_{k=1}^{K} \textbf{Exp}\left(\Delta_{D,k}; \Lambda_{m_D}\right)\right] \textbf{Gamma}\left(\Lambda_{m'_D}; \alpha_{\lambda,D}, \beta_{\lambda,D}\right), m'_D = 1,\ldots,M_D \end{aligned} \tag{S174}$$

where, $\Delta_{D,k}$ is the kth dwell time at state $m_D$, and $K$ is the total number of dwell times at state $m'_D$. So, we have

$$\Lambda_{m'_D} \sim \mathbf{Gamma}\left(\alpha', \beta'\right), m'_D = 1, \ldots, M_D \tag{S175}$$

where, $\alpha' = \alpha_{\lambda,D}$ and $\beta' = \frac{1}{\frac{1}{\beta_{\lambda,D}} + \sum_{k=1}^{K} \Delta_{D,k}}$.

Then, for the weights we have

$$\begin{aligned} p\left(\overline{\pi}^D_{m'_D}|\overline{S}_D, \{\Lambda_{m_D}\}_{m_D}, \{\overline{\pi}^D_{m_D}\}_{m_D \neq m'_D}\right) &\propto p\left(\overline{S}_D|\{\Lambda_{m_D}, \overline{\pi}^D_{m_D}\}_{m_D}\right) p\left(\overline{\pi}^D_{m_D}\right) \\ &= \mathbf{Cat}\left(\left[\sum_{t=1}^{T-1} \mathbb{I}\left(S_{D,t} = m'_D\right) \mathbb{I}\left(S_{D,t+1} = 1\right), \ldots, \sum_{t=1}^{T-1} \mathbb{I}\left(S_{D,t} = m'_D\right) \mathbb{I}\left(S_{A,t+1} = M_D\right)\right]\right) \\ &\times \mathbf{Dir}\left(\overline{\pi}^D_{m'_D}; \alpha_{\pi,D}, \overline{\beta}_{\pi,D}\right), m'_D = 1, \ldots, M_D. \end{aligned} \tag{S176}$$

So, we have

$$\begin{aligned} \overline{\pi}^D_{m'_D} \sim \mathbf{Dir}&\left(\left[\frac{\alpha_{\pi,D}}{M_D - 1} + \sum_{t=1}^{T-1} \mathbb{I}\left(S_{D,t} = m'_D\right) \mathbb{I}\left(S_{D,t+1} = 1\right), \ldots, \frac{\alpha_{\pi,D}}{M_D - 1} + \sum_{t=1}^{T-1} \mathbb{I}\left(S_{D,t} = m'_D\right) \mathbb{I}\left(S_{D,t+1} = M_D\right)\right]\right) \\ &m'_D = 1, \ldots, M_D. \end{aligned} \tag{S177}$$

#### S5.7. Update the transition matrix of FRET trajectory $\overline{\overline{Q}}_F$

After sampling the state trajectory of the FRET states we jointly sample all nondiagonal elements of the transition matrix $\overline{\overline{Q}}_F$ through the join conditional $p(\overline{\overline{Q}}_F|\eta_A, \eta_D, \mu_{\text{back},A}, \mu_{\text{back},D}, \{\tau_{A,m_A}, \ \mu_{ex,A,m_A}\}_{m_A}, \{\tau_{D,m_D}, \ \mu_{ex,D,m_D}\}_{m_D}, \{k_{F,m_F}\}_{m_F}, \overline{\overline{Q}}_D, \overline{\overline{Q}}_A, \overline{S}_A, \overline{S}_D, \overline{S}_F, \ \overline{\zeta}_A, \ \overline{\zeta}_D, \ \overline{\zeta}_F, \ \overline{\delta}, \overline{C})$ which can be simplify to $p\left(\overline{\overline{Q}}_F|\overline{S}_F\right)$. Here, the $p\left(\overline{\overline{Q}}_F|\overline{S}_F\right)$ can be rewritten as

$$p\left(\overline{\overline{Q}}_F|\overline{S}_F\right) = p\left(\{\Lambda_{m_F}, \overline{\pi}^F_{m_F}\}_{m_F}|\overline{S}_F\right). \tag{S178}$$

So, for escape rates we have

$$\begin{aligned} p\left(\Lambda_{m'_F}|\overline{S}_F, \{\Lambda_{m_F}\}_{m_F \neq m'_F}, \{\overline{\pi}^F_{m_F}\}_{m_F}\right) &\propto p\left(\overline{S}_F|\{\Lambda_{m_F}, \overline{\pi}^F_{m_F}\}_{m_F}\right) p\left(\Lambda_{m'_F}\right) \\ &= \left[\prod_{k=1}^{K} \mathbf{Exp}\left(\Delta_{F,k}; \Lambda_{m_F}\right)\right] \mathbf{Gamma}\left(\Lambda_{m'_F}; \alpha_{\lambda,F}, \beta_{\lambda,F}\right), m'_F = 1, \ldots, M_F \end{aligned} \tag{S179}$$

where, $\Delta_{F,k}$ is the kth dwell time at state $m_A$, and $K$ is the total number of dwell times at state $m'_F$. So, we have

$$\Lambda_{m'_F} \sim \mathbf{Gamma}\left(\alpha', \beta'\right), m'_F = 1, \ldots, M_F \tag{S180}$$

where, $\alpha' = \alpha_{\lambda,F}$ and $\beta' = \frac{1}{\frac{1}{\beta_{\lambda,F}} + \sum_{k=1}^{K} \Delta_{F,k}}$.

Then, for the weights we have

$$\begin{aligned} p\left(\overline{\pi}^F_{m'_F}|\overline{S}_F, \{\Lambda_{m_F}\}_{m_F}, \{\overline{\pi}^F_{m_F}\}_{m_F \neq m'_F}\right) &\propto p\left(\overline{S}_F|\{\Lambda_{m_F}, \overline{\pi}^F_{m_F}\}_{m_F}\right) p\left(\overline{\pi}^F_{m_F}\right) \\ &= \mathbf{Cat}\left(\left[\sum_{t=1}^{T-1} \mathbb{I}\left(S_{F,t} = m'_F\right) \mathbb{I}\left(S_{F,t+1} = 1\right), \ldots, \sum_{t=1}^{T-1} \mathbb{I}\left(S_{F,t} = m'_F\right) \mathbb{I}\left(S_{F,t+1} = M_F\right)\right]\right) \\ &\times \mathbf{Dir}\left(\overline{\pi}^F_{m'_F}; \alpha_{\pi,F}, \overline{\beta}_{\pi,F}\right), m'_F = 1, \ldots, M_F. \end{aligned} \tag{S181}$$

So, we have

$$\begin{aligned} \overline{\pi}^F_{m'_F} \sim \mathbf{Dir}&\left(\left[\frac{\alpha_{\pi,F}}{M_F - 1} + \sum_{t=1}^{T-1} \mathbb{I}\left(S_{F,t} = m'_F\right) \mathbb{I}\left(S_{F,t+1} = 1\right), \ldots, \frac{\alpha_{\pi,F}}{M_F - 1} + \sum_{t=1}^{T-1} \mathbb{I}\left(S_{F,t} = m'_F\right) \mathbb{I}\left(S_{F,t+1} = M_F\right)\right]\right) \\ &m'_F = 1, \ldots, M_F. \end{aligned} \tag{S182}$$

#### S5.8. Update the weight on the initial state of acceptor PIFE $\overline{\zeta}_A$

We sample such weights $\overline{\zeta}_A$ from the join conditional $p(\overline{\zeta}_A|\eta_A, \eta_D, \mu_{\text{back},A},\mu_{\text{back},D}, \{\tau_{A,m_A}, \mu_{ex,A}\}_{m_A}, \{\tau_{D,m_D}, \mu_{ex,D,m_D}\}_{m_D}, \{K_{F,m_F}\}_{m_F}, \overline{\overline{Q}}_A, \overline{\overline{Q}}_D, \overline{\overline{Q}}_F, \overline{S}_A, \overline{S}_D, \overline{S}_F, \overline{\zeta}_D, \overline{\zeta}_F, \overline{\delta}, \overline{C})$ which can be simplify to

$$p\left(\overline{\zeta}_A|S_{A,1}\right) \propto p\left(S_{A,1}|\overline{\zeta}_A\right) p\left(\overline{\zeta}_A\right). \tag{S183}$$

So, we can directly sample $\overline{\zeta}_A$

$$\overline{\zeta}_A \sim \mathbf{Dir}\left(\alpha_{A,\zeta}\overline{\beta}_{A,hyper} + \sum_{m_A=1}^{M_A} \mathbb{I}\left(S_{A,1} = m_A\right)\right). \tag{S184}$$

#### S5.9. Update the weight on initial state of donor PIFE $\overline{\zeta}_D$

The same as the acceptor case, we sample such weights $\overline{\zeta}_A$ from the join conditional $p(\overline{\zeta}_D|\eta_A, \eta_D, \mu_{\text{back},A},\mu_{\text{back},D}, \{\tau_{A,m_A},\ \mu_{ex,A}\}_{m_A}, \{\tau_{D,m_D},\ \mu_{ex,D,m_D}\}_{m_D},\ \{K_{F,m_F}\}_{m_F},\overline{\overline{Q}}_A,\ \overline{\overline{Q}}_D,\ \overline{\overline{Q}}_F,\ \overline{S}_A,\ \overline{S}_D,\ \overline{S}_F, \overline{\zeta}_A, \overline{\zeta}_F, \overline{\delta}, \overline{C})$ which can be simplify to

$$p\left(\overline{\zeta}_D|S_{D,1}\right) \propto p\left(S_{D,1}|\overline{\zeta}_D\right) p\left(\overline{\zeta}_D\right). \tag{S185}$$

So, we can directly sample $\overline{\zeta}_D$

$$\overline{\zeta}_D \sim \mathbf{Dir}\left(\alpha_{D,\zeta}\overline{\zeta}_{D,hyper} + \sum_{m_D=1}^{M_D} \mathbb{I}\left(S_{D,1} = m_D\right)\right). \tag{S186}$$

#### S5.10. Update the weight on initial state of FRET $\overline{\zeta}_F$

Same as donor and acceptor cases, we sample such weights $\overline{\zeta}_F$ from the join conditional $p(\overline{\beta}_F|\eta_A, \eta_D, \mu_{\text{back},A},\mu_{\text{back},D}, \{\tau_{A,m_A},\ \ \mu_{ex,A}\}_{m_A}, \{\tau_{D,m_D},\ \ \mu_{ex,D,m_D}\}_{m_D},\ \ \{k_{F,m_F}\}_{m_F},\ \overline{\overline{Q}}_A,\ \overline{\overline{Q}}_D,\ \overline{\overline{Q}}_F,\ \overline{S}_A,\ \overline{S}_D, \overline{S}_F, \overline{\zeta}_A, \overline{\zeta}_D, \overline{\delta}, \overline{C})$ which can be simplify to

$$p\left(\overline{\zeta}_F|S_{F,1}\right) \propto p\left(S_{F,1}|\overline{\zeta}_F\right) p\left(\overline{\zeta}_F\right). \tag{S187}$$

So, we can directly sample $\overline{\zeta}_F$

$$\overline{\zeta}_F \sim \mathbf{Dir}\left(\alpha_{F,\zeta}\overline{\beta}_{F,hyper} + \sum_{m_F=1}^{M_F} \mathbb{I}\left(S_{F,1} = m_F\right)\right). \tag{S188}$$

#### S5.11. Jointly update the acceptor lifetimes of all states $\{\tau_{A,m_A}\}_{m_A}$, donor lifetimes of all states $\{\tau_{D,m_D}\}_{m_D}$, and resonance energy transfer rates of all states $\{k_{F,m_F}\}_{m_F}$

Next, we sample the acceptor lifetimes of all states $\{\tau_{A,m_A}\}_{m_A}$, donor lifetimes of all states $\{\tau_{D,m_D}\}_{m_D}$, and resonance energy transfer rates of all states $\{k_{F,m_F}\}_{m_F}$ from their conditional distribution $p(\{\tau_{A,m_A}\}_{m_A}, \{\tau_{D,m_D}\}_{m_D}, \{k_{F,m_F}\}_{m_F}|\eta_A, \eta_D, \mu_{\text{back},A},\mu_{\text{back},D},\ \mu_{ex,A}\}_{m_A},\ \mu_{ex,D,m_D}\}_{m_D}, \{\lambda_{F,m_F}\}_{m_F},\ \overline{\overline{Q}}_A,\ \overline{\overline{Q}}_D,\ \overline{\overline{Q}}_F, \overline{S}_A, \overline{S}_D, \overline{S}_F, \overline{\zeta}_A, \overline{\zeta}_D, \overline{\zeta}_F, \overline{\delta}, \overline{C})$ which can be simplify to

$$p\left(\{\tau_{A,m_A}\}_{m_A}, \{\tau_{D,m_D}\}_{m_D}, \{k_{F,m_F}\}_{m_F}|\eta_A, \eta_D, \mu_{\text{back},A}, \mu_{\text{back},D}, \{\mu_{ex,A,m_A}\}_{m_A}, \{\mu_{ex,D,m_D}\}_{m_D}, \overline{S}_A, \overline{S}_D, \overline{S}_F, \overline{\delta}, \overline{C}\right) \sim \left[\prod_{t=1}^{T} p\left(\delta_t|\eta_A, \eta_D, \dots\right) \times p\left(C_t|\eta_A, \eta_D, \dots\right)\right] \times \left[\prod_{m_D=1}^{M_D} p\left(\tau_{D,m_D}\right)\right] \times \left[\prod_{m_A=1}^{M_A} p\left(\tau_{A,m_A}\right)\right] \times \left[\prod_{m_F=1}^{M_F} p\left(k_{F,m_F}\right)\right] \tag{S189}$$

Since, this conditional distribution has a complex form, we cannot directly sample from it. So, instead, we sample through the Metropolis Hastings algorithm and the proposals distributions in this case are

$$\tau^{\text{prop}}_{A,m_A} \sim \textbf{Gamma}\left(\alpha^{\text{prop}}_{A}, \frac{\tau^{\text{old}}_{A,m_A}}{\alpha^{\text{prop}}_{A}}\right), \qquad m_A = 1, \ldots, M_A \tag{S190}$$

$$\tau^{\text{prop}}_{D,m_D} \sim \textbf{Gamma}\left(\alpha^{\text{prop}}_{D}, \frac{\tau^{\text{old}}_{D,m_D}}{\alpha^{\text{prop}}_{D}}\right), \qquad m_D = 1, \ldots, M_D \tag{S191}$$

$$K^{\text{prop}}_{F,m_F} \sim \textbf{Gamma}\left(\alpha^{\text{prop}}_{F}, \frac{K^{\text{old}}_{F,m_F}}{\alpha^{\text{prop}}_{F}}\right) \qquad m_F = 1, \ldots, M_F \tag{S192}$$

where, $\{\tau^{\text{prop}}_{A,m_A}\}_{m_A}$, $\{\tau^{\text{prop}}_{D,m_D}\}_{m_D}$, and $\{k^{\text{prop}}_{F,m_F}\}_{m_F}$ are the proposed acceptor lifetimes, donor lifetimes, and resonance energy transfer rates, $\{\tau^{\text{old}}_{A,m_A}\}_{m_A}$, $\{\tau^{\text{old}}_{D,m_D}\}_{m_D}$, and $\{k^{\text{old}}_{F,m_F}\}_{m_F}$, are the previously sampled acceptor lifetimes, donor lifetimes, and resonance energy transfer rates, and $\alpha^{\text{prop}}_{A}$, $\alpha^{\text{prop}}_{D}$, and $\alpha^{\text{prop}}_{F}$ are the proposal distribution parameters.

Now, we need to calculate the acceptance ratio

$$
\begin{aligned}
r_{DFA} = &\left[\prod_{t=1}^{T} \frac{p\left(\delta_t | \eta_A, \eta_D, \mu_{\text{back},A}, \mu_{\text{back},D}, \{\tau^{\text{prop}}_{A,m_A}, \mu_{ex,A,m_A}\}_{m_A}, \{\tau^{\text{prop}}_{D,m_D}, \mu_{ex,D,m_D}\}_{m_D}, \{k^{\text{prop}}_{F,m_F}\}_{m_F}, S_{A,t}, S_{D,t}, S_{F,t}, C_t\right)}{p\left(\delta_t | \eta_A, \eta_D, \mu_{\text{back},A}, \mu_{\text{back},D}, \{\tau^{\text{old}}_{A,m_A}, \mu_{ex,A,m_A}\}_{m_A}, \{\tau^{\text{old}}_{D,m_D}, \mu_{ex,D,m_D}\}_{m_D}, \{k^{\text{old}}_{F,m_F}\}_{m_F}, S_{A,t}, S_{D,t}, S_{F,t}, C_t\right)}\right. \\
&\left.\times \frac{p\left(C_t | \eta_A, \eta_D, \mu_{\text{back},A}, \mu_{\text{back},D}, \{\mu_{ex,A,m_A}\}_{m_A}, \{\tau^{\text{prop}}_{D,m_D}, \mu_{ex,D,m_D}\}_{m_D}, \{k^{\text{prop}}_{F,m_F}\}_{m_F}, S_{A,t}, S_{D,t}, S_{F,t}\right)}{p\left(C_t | \eta_A, \eta_D, \mu_{\text{back},A}, \mu_{\text{back},D}, \{\mu_{ex,A,m_A}\}_{m_A}, \{\tau^{\text{old}}_{D,m_D}, \mu_{ex,D,m_D}\}_{m_D}, \{k^{\text{old}}_{F,m_F}\}_{m_F}, S_{A,t}, S_{D,t}, S_{F,t}\right)}\right] \\
&\times \left[\prod_{m_A=1}^{M_A} \frac{p\left(\tau^{\text{prop}}_{A,m_A}\right)}{p\left(\tau^{\text{old}}_{A,m_A}\right)} \frac{p\left(\tau^{\text{old}}_{A,m_A} | \tau^{\text{prop}}_{A,m_A}\right)}{p\left(\tau^{\text{prop}}_{A,m_A} | \tau^{\text{old}}_{A,m_A}\right)}\right] \times \left[\prod_{m_D=1}^{M_D} \frac{p\left(\tau^{\text{prop}}_{D,m_D}\right)}{p\left(\tau^{\text{old}}_{D,m_D}\right)} \frac{p\left(\tau^{\text{old}}_{D,m_D} | \tau^{\text{prop}}_{D,m_D}\right)}{p\left(\tau^{\text{prop}}_{D,m_D} | \tau^{\text{old}}_{D,m_D}\right)}\right] \times \left[\prod_{m_F=1}^{M_F} \frac{p\left(k^{\text{prop}}_{F,m_F}\right)}{p\left(k^{\text{old}}_{F,m_F}\right)} \frac{p\left(k^{\text{old}}_{F,m_F} | k^{\text{prop}}_{F,m_F}\right)}{p\left(k^{\text{prop}}_{F,m_F} | k^{\text{old}}_{F,m_F}\right)}\right] \\
= &\left[\prod_{t=1}^{T} \frac{p\left(\delta_t | \eta_A, \eta_D, \mu_{\text{back},A}, \mu_{\text{back},D}, \{\tau^{\text{prop}}_{A,m_A}, \mu_{ex,A,m_A}\}_{m_A}, \{\tau^{\text{prop}}_{D,m_D}, \mu_{ex,D,m_D}\}_{m_D}, \{k^{\text{prop}}_{F,m_F}\}_{m_F}, S_{A,t}, S_{D,t}, S_{F,t}, C_t\right)}{p\left(\delta_t | \eta_A, \eta_D, \mu_{\text{back},A}, \mu_{\text{back},D}, \{\tau^{\text{old}}_{A,m_A}, \mu_{ex,A,m_A}\}_{m_A}, \{\tau^{\text{old}}_{D,m_D}, \mu_{ex,D,m_D}\}_{m_D}, \{k^{\text{old}}_{F,m_F}\}_{m_F}, S_{A,t}, S_{D,t}, S_{F,t}, C_t\right)}\right. \\
&\left.\times \frac{p\left(C_t | \eta_A, \eta_D, \mu_{\text{back},A}, \mu_{\text{back},D}, \{\mu_{ex,A,m_A}\}_{m_A}, \{\tau^{\text{prop}}_{D,m_D}, \mu_{ex,D,m_D}\}_{m_D}, \{k^{\text{prop}}_{F,m_F}\}_{m_F}, S_{A,t}, S_{D,t}, S_{F,t}\right)}{p\left(C_t | \eta_A, \eta_D, \mu_{\text{back},A}, \mu_{\text{back},D}, \{\mu_{ex,A,m_A}\}_{m_A}, \{\tau^{\text{old}}_{D,m_D}, \mu_{ex,D,m_D}\}_{m_D}, \{k^{\text{old}}_{F,m_F}\}_{m_F}, S_{A,t}, S_{D,t}, S_{F,t}\right)}\right] \\
&\times \left[\prod_{m_A=1}^{M_A} \frac{\textbf{Gamma}\left(\tau^{\text{prop}}_{a,m_A}; \alpha_A, \beta_A\right)}{\textbf{Gamma}\left(\tau^{\text{old}}_{A,m_A}; \alpha_A, \beta_A\right)} \frac{\textbf{Gamma}\left(\tau^{\text{old}}_{A,m_A}; \alpha^{\text{prop}}_A, \frac{\tau^{\text{prop}}_{A,m_A}}{\alpha^{\text{prop}}_A}\right)}{\textbf{Gamma}\left(\tau^{\text{prop}}_{A,m_A}; \alpha^{\text{prop}}_A, \frac{\tau^{\text{old}}_{A,m_A}}{\alpha^{\text{prop}}_A}\right)}\right] \\
&\times \left[\prod_{m_D=1}^{M_D} \frac{\textbf{Gamma}\left(\tau^{\text{prop}}_{D,m_D}; \alpha_D, \beta_D\right)}{\textbf{Gamma}\left(\tau^{\text{old}}_{D,m_D}; \alpha_D, \beta_D\right)} \frac{\textbf{Gamma}\left(\tau^{\text{old}}_{D,m_D}; \alpha^{\text{prop}}_D, \frac{\tau^{\text{prop}}_{D,m_D}}{\alpha^{\text{prop}}_D}\right)}{\textbf{Gamma}\left(\tau^{\text{prop}}_{D,m_D}; \alpha^{\text{prop}}_D, \frac{\tau^{\text{old}}_{D,m_D}}{\alpha^{\text{prop}}_D}\right)}\right] \\
&\times \left[\prod_{m_F=1}^{M_F} \frac{\textbf{Gamma}\left(k^{\text{prop}}_{F,m_F}; \alpha_F, \beta_F\right)}{\textbf{Gamma}\left(k^{\text{old}}_{F,m_F}; \alpha_F, \beta_F\right)} \frac{\textbf{Gamma}\left(k^{\text{old}}_{F,m_F}; k^{\text{prop}}_{F,m_F}, \frac{k^{\text{prop}}_{F,m_F}}{\alpha^{\text{prop}}_F}\right)}{\textbf{Gamma}\left(k^{\text{prop}}_{F,m_F}; \alpha^{\text{prop}}_F, \frac{F^{\text{old}}_{m_F}}{\alpha^{\text{prop}}_F}\right)}\right] \\
= &\left[\prod_{t=1}^{T} \frac{p\left(\delta_t | \eta_A, \eta_D, \mu_{\text{back},A}, \mu_{\text{back},D}, \{\tau^{\text{prop}}_{A,m_A}, \mu_{ex,A,m_A}\}_{m_A}, \{\tau^{\text{prop}}_{D,m_D}, \mu_{ex,D,m_D}\}_{m_D}, \{k^{\text{prop}}_{F,m_F}\}_{m_F}, S_{A,t}, S_{D,t}, S_{F,t}, C_t\right)}{p\left(\delta_t | \eta_A, \eta_D, \mu_{\text{back},A}, \mu_{\text{back},D}, \{\tau^{\text{old}}_{A,m_A}, \mu_{ex,A,m_A}\}_{m_A}, \{\tau^{\text{old}}_{D,m_D}, \mu_{ex,D,m_D}\}_{m_D}, \{k^{\text{old}}_{F,m_F}\}_{m_F}, S_{A,t}, S_{D,t}, S_{F,t}, C_t\right)}\right. \\
&\left.\times \frac{p\left(C_t | \eta_A, \eta_D, \mu_{\text{back},A}, \mu_{\text{back},D}, \{\mu_{ex,A,m_A}\}_{m_A}, \{\tau^{\text{prop}}_{D,m_D}, \mu_{ex,D,m_D}\}_{m_D}, \{k^{\text{prop}}_{F,m_F}\}_{m_F}, S_{A,t}, S_{D,t}, S_{F,t}\right)}{p\left(C_t | \eta_A, \eta_D, \mu_{\text{back},A}, \mu_{\text{back},D}, \{\mu_{ex,A,m_A}\}_{m_A}, \{\tau^{\text{old}}_{D,m_D}, \mu_{ex,D,m_D}\}_{m_D}, \{k^{\text{old}}_{F,m_F}\}_{m_F}, S_{A,t}, S_{D,t}, S_{F,t}\right)}\right] \\
&\times \left[\prod_{m_A=1}^{M_A} \left(\frac{\tau^{\text{old}}_{A,m_A}}{\tau^{\text{prop}}_{A,m_A}}\right)^{2\alpha^{\text{prop}}_A - \alpha_A} \exp\left(\alpha^{\text{prop}}_A \left(\frac{\tau^{\text{prop}}_{A,m_A}}{\tau^{\text{old}}_{A,m_A}} - \frac{\tau^{\text{old}}_{A,m_A}}{\tau^{\text{prop}}_{A,m_A}}\right)\right) \exp\left(\frac{\tau^{\text{old}}_{A,m_a} - \tau^{\text{prop}}_{A,m_A}}{\beta_A}\right)\right] \\
&\times \left[\prod_{m_D=1}^{M_D} \left(\frac{\tau^{\text{old}}_{D,m_D}}{\tau^{\text{prop}}_{D,m_D}}\right)^{2\alpha^{\text{prop}}_D - \alpha_D} \exp\left(\alpha^{\text{prop}}_D \left(\frac{\tau^{\text{prop}}_{D,m_D}}{\tau^{\text{old}}_{D,m_D}} - \frac{\tau^{\text{old}}_{D,m_D}}{\tau^{\text{prop}}_{D,m_D}}\right)\right) \exp\left(\frac{\tau^{\text{old}}_{D,m_D} - \tau^{\text{prop}}_{D,m_D}}{\beta_D}\right)\right] \\
&\times \left[\prod_{m_F=1}^{M_F} \left(\frac{k^{\text{old}}_{F,m_F}}{k^{\text{prop}}_{F,m_F}}\right)^{2\alpha^{\text{prop}}_F - \alpha_F} \exp\left(\alpha^{\text{prop}}_F \left(\frac{k^{\text{prop}}_{F,m_F}}{k^{\text{old}}_{F,m_F}} - \frac{k^{\text{old}}_{F,m_F}}{k^{\text{prop}}_{F,m_F}}\right)\right) \exp\left(\frac{k^{\text{old}}_{F,m_F} - k^{\text{prop}}_{F,m_F}}{\beta_F}\right)\right]
\end{aligned}
\tag{S193}
$$

#### S5.12. Jointly update the cross-talk ratios of acceptor and donor channels, $\eta_A$ and $\eta_D$

In the case that the values of the cross-talk are not measured by pre-calibrations, we sample cross-talk ratios of donor and acceptor channels, $\eta_D$ and $\eta_A$, from their conditional distribution $p(\eta_A, \eta_D|\ \mu_{\text{back},A},\ \mu_{\text{back},B},\ \{\tau_{A,m_A},\ \mu_{ex,A,m_A}\}_{m_A},\ \{\tau_{D,m_D}, \mu_{ex,D,m_D}\}_{m_D},\ \{k_{F,m_F}\}_{m_F}\ \overline{\overline{Q}}_A,\ \overline{\overline{Q}}_D,\ \overline{\overline{Q}}_F,\ \overline{S}_A,\ \overline{S}_D,\ \overline{S}_F,\ \overline{\zeta}_A,\ \overline{\zeta}_D,\ \overline{\zeta}_F,\ \overline{\delta}, \overline{C})$ which can be simplify to

$$
\begin{aligned}
&p\left(\eta_A, \eta_D|\mu_{\text{back},A}, \mu_{\text{back},D}, \{\tau_{A,m_A}, \mu_{ex,A,m_A}\}_{m_A}, \{\tau_{D,m_D}, \mu_{ex,D,m_D}\}_{m_D}, \{k_{F,m_F}\}_{m_F}, \overline{S}_A, \overline{S}_D, \overline{S}_F, \overline{\delta}, \overline{C}\right) \sim \\
&\left[\prod_{t=1}^{T} p\left(\delta_t|\eta_A, \eta_D, \mu_{\text{back},A}, \mu_{\text{back},D}, \{\tau_{A,m_A}, \mu_{ex,A,m_A}\}_{m_A}, \{\tau_{D,m_D}, \mu_{ex,D,m_D}\}_{m_D}, \{k_{F,m_F}\}_{m_F}, S_{A,t}, S_{D,t}, S_{F,t}, C_t\right)\right. \\
&\quad \left. \times\, p\left(C_t|\eta_A, \eta_D, \mu_{\text{back},A}, \mu_{\text{back},D}, \{\mu_{ex,A,m_A}\}_{m_A}, \{\mu_{ex,D,m_D}\}_{m_D}, \{k_{F,m_F}\}_{m_F}, S_{A,t}, S_{D,t}, S_{F,t}\right)\right] \\
&\quad p\left(\eta_A\right) p\left(\eta_D\right).
\end{aligned}
\tag{S194}
$$

Since, direct sampling from this conditional distribution is not possible, we sample the $\eta_D$ and $\eta_A$ through the Metropolis algorithm. The proposals are

$$
\eta_A^{\text{prop}} \sim \textbf{Beta}\left(\alpha_{\eta,A}^{\text{prop}}, \alpha_{\eta,A}^{\text{prop}}\left(\frac{1}{\eta_D^{\text{old}}} - 1\right)\right) \tag{S195}
$$

$$
\eta_D^{\text{prop}} \sim \textbf{Beta}\left(\alpha_{\eta,D}^{\text{prop}}, \alpha_{\eta,D}^{\text{prop}}\left(\frac{1}{\eta_D^{\text{old}}} - 1\right)\right) \tag{S196}
$$

where, $\eta_A^{\text{prop}}$ and $\eta_D^{\text{prop}}$ are the proposed acceptor and donor channel detection efficiencies, $\eta_A^{\text{old}}$ and $\eta_D^{\text{old}}$ are the previous acceptor and donor channel detection efficiencies, and $\alpha_{\eta,A}^{\text{prop}}$ and $\alpha_{\eta,D}^{\text{prop}}$ are the proposal distribution parameters.

$$
\begin{aligned}
r_\eta = &\left[\prod_{t=1}^{T} \frac{p\left(\delta_t|\eta_A^{\text{prop}}, \eta_D^{\text{prop}}, \mu_{\text{back},A}, \mu_{\text{back},D}, \{\tau_{A,m_A}, \mu_{ex,A,m_A}\}_{m_A}, \{\tau_{D,m_D}, \mu_{ex,D,m_D}\}_{m_D}, \{k_{F,m_F}\}_{m_F}, S_{A,t}, S_{D,t}, S_{F,t}, C_t\right)}{p\left(\delta_t|\eta_A^{\text{old}}, \eta_D^{\text{old}}, \mu_{\text{back},A}, \mu_{\text{back},D}, \{\tau_{A,m_A}, \mu_{ex,A,m_A}\}_{m_A}, \{\tau_{D,m_D}, \mu_{ex,D,m_D}\}_{m_D}, \{k_{F,m_F}\}_{m_F}, S_{A,t}, S_{D,t}, S_{F,t}, C_t\right)}\right. \\
&\left. \times \frac{p\left(C_t|\eta_A^{\text{prop}}, \eta_D^{\text{prop}}, \mu_{\text{back},A}, \mu_{\text{back},D}, \{\mu_{ex,A,m_A}\}_{m_A}, \{\tau_{D,m_D}, \mu_{ex,D,m_D}\}_{m_D}, \{k_{F,m_F}\}_{m_F}, S_{A,t}, S_{D,t}, S_{F,t}\right)}{p\left(C_t|\eta_A^{\text{old}}, \eta_D^{\text{old}}, \mu_{\text{back},A}, \mu_{\text{back},D}, \{\mu_{ex,A,m_A}\}_{m_A}, \{\tau_{D,m_D}, \mu_{ex,D,m_D}\}_{m_D}, \{k_{F,m_F}\}_{m_F}, S_{A,t}, S_{D,t}, S_{F,t}\right)}\right] \\
\times &\frac{p\left(\eta_A^{\text{prop}}\right)}{p\left(\eta_A^{\text{old}}\right)} \frac{p\left(\eta_D^{\text{prop}}\right)}{p\left(\eta_D^{\text{old}}\right)} \frac{p\left(\eta_A^{\text{old}}|\eta_A^{\text{prop}}\right)}{p\left(\eta_A^{\text{prop}}|\eta_A^{\text{old}}\right)} \frac{p\left(\eta_D^{\text{old}}|\eta_D^{\text{prop}}\right)}{p\left(\eta_D^{\text{prop}}|\eta_D^{\text{old}}\right)} \\
= &\left[\prod_{t=1}^{T} \frac{p\left(\delta_t|\eta_A^{\text{prop}}, \eta_D^{\text{prop}}, \mu_{\text{back},A}, \mu_{\text{back},D}, \{\tau_{A,m_A}, \mu_{ex,A,m_A}\}_{m_A}, \{\tau_{D,m_D}, \mu_{ex,D,m_D}\}_{m_D}, \{k_{F,m_F}\}_{m_F}, S_{A,t}, S_{D,t}, S_{F,t}, C_t\right)}{p\left(\delta_t|\eta_A^{\text{old}}, \eta_D^{\text{old}}, \mu_{\text{back},A}, \mu_{\text{back},D}, \{\tau_{A,m_A}, \mu_{ex,A,m_A}\}_{m_A}, \{\tau_{D,m_D}, \mu_{ex,D,m_D}\}_{m_D}, \{k_{F,m_F}\}_{m_F}, S_{A,t}, S_{D,t}, S_{F,t}, C_t\right)}\right. \\
&\left. \times \frac{p\left(C_t|\eta_A^{\text{prop}}, \eta_D^{\text{prop}}, \mu_{\text{back},A}, \mu_{\text{back},D}, \{\mu_{ex,A,m_A}\}_{m_A}, \{\tau_{D,m_D}, \mu_{ex,D,m_D}\}_{m_D}, \{k_{F,m_F}\}_{m_F}, S_{A,t}, S_{D,t}, S_{F,t}\right)}{p\left(C_t|\eta_A^{\text{old}}, \eta_D^{\text{old}}, \mu_{\text{back},A}, \mu_{\text{back},D}, \{\mu_{ex,A,m_A}\}_{m_A}, \{\tau_{D,m_D}, \mu_{ex,D,m_D}\}_{m_D}, \{k_{F,m_F}\}_{m_F}, S_{A,t}, S_{D,t}, S_{F,t}\right)}\right] \\
&\times \frac{\textbf{Beta}\left(\eta_A^{\text{prop}}; \alpha_{\eta,A}, \beta_{\eta,A}\right)}{\textbf{Beta}\left(\eta_A^{\text{old}}; \alpha_{\eta,A}, \beta_{\eta,A}\right)} \frac{\textbf{Beta}\left(\eta_A^{\text{old}}; \alpha_{\eta,A}^{\text{prop}}, \alpha_{\eta,A}^{\text{prop}}\left(\frac{1}{\eta_A^{\text{prop}}} - 1\right)\right)}{\textbf{Beta}\left(\eta_A^{\text{prop}}; \alpha_{\eta,A}^{\text{prop}}, \alpha_{\eta,A}^{\text{prop}}\left(\frac{1}{\eta_A^{\text{old}}} - 1\right)\right)} \\
&\times \frac{\textbf{Beta}\left(\eta_D^{\text{prop}}; \alpha_{\eta,D}, \beta_{\eta,D}\right)}{\textbf{Beta}\left(\eta_D^{\text{old}}; \alpha_{\eta,D}, \beta_{\eta,D}\right)} \frac{\textbf{Beta}\left(\eta_D^{\text{old}}; \alpha_{\eta,D}^{\text{prop}}, \alpha_{\eta,D}^{\text{prop}}\left(\frac{1}{\eta_D^{\text{prop}}} - 1\right)\right)}{\textbf{Beta}\left(\eta_D^{\text{prop}}; \alpha_{\eta,D}^{\text{prop}}, \alpha_{\eta,D}^{\text{prop}}\left(\frac{1}{\eta_D^{\text{old}}} - 1\right)\right)}
\end{aligned}
\tag{S197}
$$

$$
\begin{aligned}
&= \left[ \prod_{t=1}^{T} \frac{p\left(\delta_t|\eta_A^{\text{prop}}, \eta_D^{\text{prop}}, \mu_{\text{back},A}, \mu_{\text{back},D}, \{\tau_{A,m_A}, \mu_{ex,A,m_A}\}_{m_A}, \{\tau_{D,m_D}, \mu_{ex,D,m_D}\}_{m_D}, \{k_{F,m_F}\}_{m_F}, S_{A,t}, S_{D,t}, S_{F,t}, C_t\right)}{p\left(\delta_t|\eta_A^{\text{old}}, \eta_D^{\text{old}}, \mu_{\text{back},A}, \mu_{\text{back},D}, \{\tau_{A,m_A}, \mu_{ex,A,m_A}\}_{m_A}, \{\tau_{D,m_D}, \mu_{ex,D,m_D}\}_{m_D}, \{k_{F,m_F}\}_{m_F}, S_{A,t}, S_{D,t}, S_{F,t}, C_t\right)} \right. \\
&\quad \left. \times \frac{p\left(C_t|\eta_A^{\text{prop}}, \eta_D^{\text{prop}}, \mu_{\text{back},A}, \mu_{\text{back},D}, \{\mu_{ex,A,m_A}\}_{m_A}, \{\tau_{D,m_D}, \mu_{ex,D,m_D}\}_{m_D}, \{k_{F,m_F}\}_{m_F}, S_{A,t}, S_{D,t}, S_{F,t}\right)}{p\left(C_t|\eta_A^{\text{old}}, \eta_D^{\text{old}}, \mu_{\text{back},A}, \mu_{\text{back},D}, \{\mu_{ex,A,m_A}\}_{m_A}, \{\tau_{D,m_D}, \mu_{ex,D,m_D}\}_{m_D}, \{k_{F,m_F}\}_{m_F}, S_{A,t}, S_{D,t}, S_{F,t}\right)} \right] \\
&\quad \times \frac{\Gamma\left(\frac{\alpha^{\text{prop}}_{\eta,A}}{\eta_A^{\text{prop}}}\right)\Gamma\left(\alpha^{\text{prop}}_{\eta,A}\left(\frac{1}{\eta_A^{\text{old}}}-1\right)\right)}{\Gamma\left(\frac{\alpha^{\text{prop}}_{\eta,A}}{\eta_A^{\text{old}}}\right)\Gamma\left(\alpha^{\text{prop}}_{\eta,A}\left(\frac{1}{\eta_A^{\text{prop}}}-1\right)\right)} \left(\frac{\eta_A^{\text{old}}}{\eta_A^{\text{prop}}}\right)^{\alpha^{\text{prop}}_{\eta,A}-\alpha_{\eta,A}} \frac{\left(1-\eta_A^{\text{old}}\right)^{\alpha^{\text{prop}}_{\eta,A}\left(\frac{1}{\eta_A^{\text{prop}}}-1\right)-\beta_{\eta,A}}}{\left(1-\eta_A^{\text{prop}}\right)^{\alpha^{\text{prop}}_{\eta,A}\left(\frac{1}{\eta_A^{\text{old}}}-1\right)-\beta_{\eta,A}}} \\
&\quad \times \frac{\Gamma\left(\frac{\alpha^{\text{prop}}_{\eta,D}}{\eta_D^{\text{prop}}}\right)\Gamma\left(\alpha^{\text{prop}}_{\eta,D}\left(\frac{1}{\eta_D^{\text{old}}}-1\right)\right)}{\Gamma\left(\frac{\alpha^{\text{prop}}_{\eta,D}}{\eta_D^{\text{old}}}\right)\Gamma\left(\alpha^{\text{prop}}_{\eta,D}\left(\frac{1}{\eta_D^{\text{prop}}}-1\right)\right)} \left(\frac{\eta_D^{\text{old}}}{\eta_D^{\text{prop}}}\right)^{\alpha^{\text{prop}}_{\eta,D}-\alpha_{\eta,D}} \frac{\left(1-\eta_D^{\text{old}}\right)^{\alpha^{\text{prop}}_{\eta,D}\left(\frac{1}{\eta_D^{\text{prop}}}-1\right)-\beta_{\eta,D}}}{\left(1-\eta_D^{\text{prop}}\right)^{\alpha^{\text{prop}}_{\eta,D}\left(\frac{1}{\eta_D^{\text{old}}}-1\right)-\beta_{\eta,D}}}
\end{aligned}
\tag{S198}
$$

#### S5.13. Jointly update the acceptor excitation rates $\{\mu_{ex,A,m_A}\}_{m_A}$, donor excitation rates $\{\mu_{ex,D,m_D}\}_{m_D}$, acceptor channel background photon emission rate $\mu_{\textbf{back},A}$, and donor channel background photon emission rate $\mu_{\textbf{back},D}$

Next, we sample the acceptor excitation rates $\{\mu_{ex,A,m_A}\}_{m_A}$, donor excitation states $\{\mu_{ex,D,m_D}\}_{m_D}$, acceptor channel background $\mu_{\text{back},A}$, and donor channel background $\mu_{\text{back},D}$ from their conditional distribution $p(\{\mu_{ex,A,m_A}\}_{m_A}, \{\mu_{ex,D,m_D}\}_{m_D}, \mu_{\text{back},A}, \mu_{\text{back},D}|\eta_A, \eta_D, \{\tau_{A,m_A}\}_{m_A}, \{\tau_{D,m_D}\}_{m_D}, \{k_{F,m_F}\}_{m_F}, \overline{\overline{Q}}_A, \overline{\overline{Q}}_D, \overline{\overline{Q}}_F, \overline{S}_A, \overline{S}_D, \overline{S}_F, \overline{\zeta}_A, \overline{\zeta}_D, \overline{\zeta}_F, \overline{\delta}, \overline{C})$ which can be simplify to

$$
\begin{aligned}
&p\left(\{\mu_{ex,A,m_A}\}_{m_A}, \{\mu_{ex,D,m_D}\}_{m_D}, \mu_{\text{back},A}, \mu_{\text{back},D}|\eta_A, \eta_D, \{\tau_{A,m_A}\}_{m_A}, \{\tau_{D,m_D}\}_{m_D}, \{k_{F,m_F}\}_{m_F}, \dots, \overline{\delta}, \overline{C}\right) \sim \\
&\left[\prod_{t=1}^{T} p\left(\delta_t|\eta_A, \eta_D, \{\tau_{A,m_A}, \mu_{ex,A,m_A}\}_{m_A}, \dots\right) \times p\left(C_t|\eta_A, \eta_D, \{\mu_{ex,A,m_A}\}_{m_A}, \dots\right)\right] \\
&\times \left[\prod_{m_A=1}^{M_A} p\left(\mu_{ex,A,m_A}\right)\right]\left[\prod_{m_D=1}^{M_D} p\left(\mu_{ex,D,m_D}\right)\right] p\left(\mu_{\text{back},A}\right) p\left(\mu_{\text{back},D}\right).
\end{aligned}
\tag{S199}
$$

Since, this conditional distribution has a complex form, we cannot directly sample from it. So, instead, we sample through the Metropolis Hastings algorithm and the proposals distributions in this case are

$$
\mu^{\text{prop}}_{ex,D,m_D} \sim \textbf{Gamma}\left(\alpha^{\text{prop}}_{ex,D}, \frac{\mu^{\text{old}}_{ex,D,m_D}}{\alpha^{\text{prop}}_{ex,D}}\right), \qquad m_D = 1, \dots, M_D \tag{S200}
$$

$$
\mu^{\text{prop}}_{ex,A,m_A} \sim \textbf{Gamma}\left(\alpha^{\text{prop}}_{ex,A}, \frac{\mu^{\text{old}}_{ex,A,m_A}}{\alpha^{\text{prop}}_{ex,A}}\right), \qquad m_A = 1, \dots, M_A \tag{S201}
$$

$$
\mu^{\text{prop}}_{\text{back},D} \sim \textbf{Gamma}\left(\alpha^{\text{prop}}_{\text{back},D}, \frac{\mu^{\text{old}}_{\text{back},D}}{\alpha^{\text{prop}}_{\text{back},D}}\right) \tag{S202}
$$

$$
\mu^{\text{prop}}_{\text{back},A} \sim \textbf{Gamma}\left(\alpha^{\text{prop}}_{\text{back},A}, \frac{\mu^{\text{old}}_{\text{back},A}}{\alpha^{\text{prop}}_{\text{back},A}}\right) \tag{S203}
$$

where, $\{\mu^{\text{prop}}_{ex,A,m_A}\}_{m_A}$, $\{\mu^{\text{prop}}_{ex,D,m_D}\}_{m_D}$, and $\mu^{\text{prop}}_{\text{back},A}$, and $\mu^{\text{prop}}_{\text{back},D}$ are the proposed acceptor excitation rates, donor excitation rate, acceptor channel background , and donor channel background, and $\{\mu^{\text{old}}_{ex,D,m_D}\}_{m_D}$, $\{\mu^{\text{old}}_{ex,A,m_A}\}_{m_A}$, and $\mu^{\text{old}}_{\text{back},A}$ and $\mu^{\text{old}}_{\text{back},D}$ are the previously sampled values, and $\alpha^{\text{prop}}_{ex,A}$, $\alpha^{\text{prop}}_{ex,D}$, $\alpha^{\text{prop}}_{\text{back},D}$ and $\alpha^{\text{prop}}_{\text{back},D}$ are the proposal distribution parameters. Now, we need to calculate the acceptance ratio

$$
\begin{aligned}
r_{AD_{ex}B} &= \Bigg[ \prod_{t=1}^{T} \frac{p\left(\delta_t | \eta_A, \eta_D, \mu_{\text{back},A}^{\text{prop}}, \mu_{\text{back},D}^{\text{prop}}, \{\tau_{A,m_A}, \mu_{ex,A,m_A}^{\text{prop}}\}_{m_A}, \{\tau_{D,m_D}, \mu_{ex,D,m_D}^{\text{prop}}\}_{m_D}, \{K_{F,m_F}\}_{m_F}, S_{A,t}, S_{D,t}, S_{F,t}, C_t\right)}{p\left(\delta_t | \eta_A, \eta_D, \mu_{\text{back},A}^{\text{old}}, \mu_{\text{back},D}^{\text{old}}, \{\tau_{A,m_A}, \mu_{ex,A,m_A}^{\text{old}}\}_{m_A}, \{\tau_{D,m_D}, \mu_{ex,D,m_D}^{\text{old}}\}_{m_D}, \{K_{F,m_F}\}_{m_F}, , S_{A,t}, S_{D,t}, S_{F,t}, C_t\right)} \\
&\quad \times \frac{p\left(C_t | \eta_A, \eta_D, \mu_{\text{back},A}^{\text{prop}}, \mu_{\text{back},D}^{\text{prop}}, \{\mu_{ex,A,m_A}^{\text{prop}}\}_{m_A}, \{\tau_{D,m_D}, \mu_{ex,D,m_D}^{\text{prop}}\}_{m_D}, \{K_{F,m_F}\}_{m_F}, S_{A,t}, S_{D,t}, S_{F,t}\right)}{p\left(C_t | \eta_A, \eta_D, \mu_{\text{back},A}^{\text{old}}, \mu_{\text{back},D}^{\text{old}}, \{\mu_{ex,A,m_A}^{\text{old}}\}_{m_A}, \{\tau_{D,m_D}, \mu_{ex,D,m_D}^{\text{old}}\}_{m_D}, \{K_{F,m_F}\}_{m_F}, S_{A,t}, S_{D,t}, S_{F,t}\right)} \Bigg] \\
&\times \left[ \prod_{m_A=1}^{M_A} \frac{p\left(\mu_{ex,A,m_A}^{\text{prop}}\right)}{p\left(\mu_{ex,A,m_A}^{\text{old}}\right)} \frac{p\left(\mu_{ex,A,m_A}^{\text{old}} | \mu_{ex,A,m_A}^{\text{prop}}\right)}{p\left(\mu_{ex,A,m_A}^{\text{prop}} | \mu_{ex,A,m_A}^{\text{old}}\right)} \right] \times \left[ \prod_{m_D=1}^{M_D} \frac{p\left(\mu_{ex,D,m_D}^{\text{prop}}\right)}{p\left(\mu_{ex,D,m_D}^{\text{old}}\right)} \frac{p\left(\mu_{ex,D,m_D}^{\text{old}} | \mu_{ex,D,m_D}^{\text{prop}}\right)}{p\left(\mu_{ex,D,m_D}^{\text{prop}} | \mu_{ex,D,m_D}^{\text{old}}\right)} \right] \\
&\quad \times \frac{p\left(\mu_{\text{back},A}^{\text{prop}}\right)}{p\left(\mu_{\text{back},A}^{\text{old}}\right)} \frac{p\left(\mu_{\text{back},A}^{\text{old}} | \mu_{\text{back},A}^{\text{prop}}\right)}{p\left(\mu_{\text{back},A}^{\text{prop}} | \mu_{\text{back},A}^{\text{old}}\right)} \frac{p\left(\mu_{\text{back},D}^{\text{prop}}\right)}{p\left(\mu_{\text{back},D}^{\text{old}}\right)} \frac{p\left(\mu_{\text{back},D}^{\text{old}} | \mu_{\text{back},D}^{\text{prop}}\right)}{p\left(\mu_{\text{back},D}^{\text{prop}} | \mu_{\text{back},D}^{\text{old}}\right)} \\
&= \Bigg[ \prod_{t=1}^{T} \frac{p\left(\delta_t | \eta_A, \eta_D, \mu_{\text{back},D}^{\text{prop}}, \mu_{\text{back},A}^{\text{prop}}, \{\tau_{A,m_A}, \mu_{ex,A,m_A}^{\text{prop}}\}_{m_D}, \{\tau_{D,m_D}, \mu_{ex,D,m_D}^{\text{prop}}\}_{m_D}, \{K_{F,m_F}\}_{m_F}, S_{A,t}, S_{D,t}, S_{F,t}, C_t\right)}{p\left(\delta_t | \eta_A, \eta_D, \mu_{\text{back},A}^{\text{old}}, \mu_{\text{back},D}^{\text{old}}, \{\tau_{A,m_A}, \mu_{ex,A,m_A}^{\text{old}}\}_{m_A}, \{\tau_{D,m_D}, \mu_{ex,D,m_D}^{\text{old}}\}_{m_D}, \{K_{F,m_F}\}_{m_F}, , S_{A,t}, S_{D,t}, S_{F,t}, C_t\right)} \\
&\quad \times \frac{p\left(C_t | \eta_A, \eta_D, \mu_{\text{back},A}^{\text{prop}}, \mu_{\text{back},D}^{\text{prop}}, \{\mu_{ex,A,m_A}^{\text{prop}}\}_{m_A}, \{\tau_{D,m_D}, \mu_{ex,D,m_D}^{\text{prop}}\}_{m_D}, \{K_{F,m_F}\}_{m_F}, S_{A,t}, S_{D,t}, S_{F,t}\right)}{p\left(C_t | \eta_A, \eta_D, \mu_{\text{back},A}^{\text{old}}, \mu_{\text{back},D}^{\text{old}}, \{\mu_{ex,A,m_A}^{\text{old}}\}_{m_A}, \{\tau_{D,m_D}, \mu_{ex,D,m_D}^{\text{old}}\}_{m_D}, \{K_{F,m_F}\}_{m_F}, S_{A,t}, S_{D,t}, S_{F,t}\right)} \Bigg] \\
&\quad \times \left[ \prod_{m_A=1}^{M_A} \frac{\textbf{Gamma}\left(\mu_{ex,A,m_A}^{\text{prop}}; \alpha_{ex,A}, \beta_{ex,A}\right)}{\textbf{Gamma}\left(\mu_{ex,A,m_A}^{\text{old}}; \alpha_{ex,A}, \beta_{ex,A}\right)} \frac{\textbf{Gamma}\left(\mu_{ex,A,m_A}^{\text{old}}; \alpha_{ex,A}^{\text{prop}}, \frac{\mu_{ex,A,m_A}^{\text{prop}}}{\alpha_{ex,A}^{\text{prop}}}\right)}{\textbf{Gamma}\left(\mu_{ex,A,m_A}^{\text{prop}}; \alpha_{ex,A}^{\text{prop}}, \frac{\mu_{ex,A,m_A}^{\text{old}}}{\alpha_{ex,A}^{\text{prop}}}\right)} \right] \\
&\quad \times \left[ \prod_{m_D=1}^{M_D} \frac{\textbf{Gamma}\left(\mu_{ex,D,m_D}^{\text{prop}}; \alpha_{ex,D}, \beta_{ex,D}\right)}{\textbf{Gamma}\left(\mu_{ex,D,m_D}^{\text{old}}; \alpha_{ex,D}, \beta_{ex,D}\right)} \frac{\textbf{Gamma}\left(\mu_{ex,D,m_D}^{\text{old}}; \alpha_{ex,D}^{\text{prop}}, \frac{\mu_{ex,D,m_D}^{\text{prop}}}{\alpha_{ex,D}^{\text{prop}}}\right)}{\textbf{Gamma}\left(\mu_{ex,D,m_D}^{\text{prop}}; \alpha_{ex,D}^{\text{prop}}, \frac{\mu_{ex,D,m_D}^{\text{old}}}{\alpha_{ex,D}^{\text{prop}}}\right)} \right] \\
&\quad \times \frac{\textbf{Gamma}\left(\mu_{\text{back},A}^{\text{prop}}; \alpha_{\text{back},A}, \beta_{\text{back},A}\right)}{\textbf{Gamma}\left(\mu_{\text{back},A}^{\text{old}}; \alpha_{\text{back},A}, \beta_{\text{back},A}\right)} \frac{\textbf{Gamma}\left(\mu_{\text{back},A}^{\text{old}}; \alpha_{\text{back},A}^{\text{prop}}, \frac{\mu_{\text{back},A}^{\text{prop}}}{\alpha_{\text{back},A}^{\text{prop}}}\right)}{\textbf{Gamma}\left(\mu_{\text{back},A}^{\text{prop}}; \alpha_{\text{back},A}^{\text{prop}}, \frac{\mu_{\text{back},A}^{\text{old}}}{\alpha_{\text{back},A}^{\text{prop}}}\right)} \\
&\quad \times \frac{\textbf{Gamma}\left(\mu_{\text{back},D}^{\text{prop}}; \alpha_{\text{back},D}, \beta_{\text{back},D}\right)}{\textbf{Gamma}\left(\mu_{\text{back},D}^{\text{old}}; \alpha_{\text{back},D}, \beta_{\text{back},D}\right)} \frac{\textbf{Gamma}\left(\mu_{\text{back},D}^{\text{old}}; \alpha_{\text{back},D}^{\text{prop}}, \frac{\mu_{\text{back},D}^{\text{prop}}}{\alpha_{\text{back},D}^{\text{prop}}}\right)}{\textbf{Gamma}\left(\mu_{\text{back},D}^{\text{prop}}; \alpha_{\text{back},D}^{\text{prop}}, \frac{\mu_{\text{back},D}^{\text{old}}}{\alpha_{\text{back},D}^{\text{prop}}}\right)} \\
&= \Bigg[ \prod_{t=1}^{T} \frac{p\left(\delta_t | \eta_A, \eta_D, \mu_{\text{back},A}^{\text{prop}}, \mu_{\text{back},D}^{\text{prop}}, \{\tau_{A,m_A}, \mu_{ex,A,m_A}^{\text{prop}}\}_{m_A}, \{\tau_{D,m_D}, \mu_{ex,D,m_D}^{\text{prop}}\}_{m_D}, \{K_{F,m_F}\}_{m_F}, S_{A,t}, S_{D,t}, S_{F,t}, C_t\right)}{p\left(\delta_t | \eta_A, \eta_D, \mu_{\text{back},A}^{\text{old}}, \mu_{\text{back},D}^{\text{old}}, \{\tau_{A,m_A}, \mu_{ex,A,m_A}^{\text{old}}\}_{m_A}, \{\tau_{D,m_D}, \mu_{ex,D,m_D}^{\text{old}}\}_{m_D}, \{K_{F,m_F}\}_{m_F}, , S_{A,t}, S_{D,t}, S_{F,t}, C_t\right)} \\
&\quad \times \frac{p\left(C_t | \eta_A, \eta_D, \mu_{\text{back},A}^{\text{prop}}, \mu_{\text{back},D}^{\text{prop}}, \{\mu_{ex,A,m_A}^{\text{prop}}\}_{m_A}, \{\tau_{D,m_D}, \mu_{ex,D,m_D}^{\text{prop}}\}_{m_D}, \{K_{F,m_F}\}_{m_F}, S_{A,t}, S_{D,t}, S_{F,t}\right)}{p\left(C_t | \eta_A, \eta_D, \mu_{\text{back},A}^{\text{old}}, \mu_{\text{back},D}^{\text{old}}, \{\mu_{ex,A,m_A}^{\text{old}}\}_{m_A}, \{\tau_{D,m_D}, \mu_{ex,D,m_D}^{\text{old}}\}_{m_D}, \{K_{F,m_F}\}_{m_F}, S_{A,t}, S_{D,t}, S_{F,t}\right)} \Bigg] \\
&\quad \times \left[ \prod_{m_A=1}^{M_A} \left(\frac{\mu_{ex,A,m_A}^{\text{old}}}{\mu_{ex,A,m_A}^{\text{prop}}}\right)^{2\alpha_{ex,A}^{\text{prop}} - \alpha_{ex,A}} \exp\left(\alpha_{ex,A}^{\text{prop}} \left(\frac{\mu_{ex,A,m_A}^{\text{prop}}}{\mu_{ex,A,m_A}^{\text{old}}} - \frac{\mu_{ex,A,m_A}^{\text{old}}}{\mu_{ex,A,m_A}^{\text{prop}}}\right)\right) \exp\left(\frac{\mu_{ex,A,m_A}^{\text{old}} - \mu_{ex,A,m_A}^{\text{prop}}}{\beta_{ex,A}}\right) \right] \\
&\quad \times \left[ \prod_{m_D=1}^{M_D} \left(\frac{\mu_{ex,D,m_D}^{\text{old}}}{\mu_{ex,D,m_D}^{\text{prop}}}\right)^{2\alpha_{ex,D}^{\text{prop}} - \alpha_{ex,D}} \exp\left(\alpha_{ex,D}^{\text{prop}} \left(\frac{\mu_{ex,D,m_D}^{\text{prop}}}{\mu_{ex,D,m_D}^{\text{old}}} - \frac{\mu_{ex,D,m_D}^{\text{old}}}{\mu_{ex,D,m_D}^{\text{prop}}}\right)\right) \exp\left(\frac{\mu_{ex,D,m_D}^{\text{old}} - \mu_{ex,D,m_D}^{\text{prop}}}{\beta_{ex,D}}\right) \right] \\
&\quad \times \left(\frac{\mu_{\text{back},A}^{\text{old}}}{\mu_{\text{back},A}^{\text{prop}}}\right)^{2\alpha_{\text{back},A}^{\text{prop}} - \alpha_{\text{back},A}} \exp\left(\alpha_{\text{back},A}^{\text{prop}} \left(\frac{\mu_{\text{back},A}^{\text{prop}}}{\mu_{\text{back},A}^{\text{old}}} - \frac{\mu_{\text{back},A}^{\text{old}}}{\mu_{\text{back},A}^{\text{prop}}}\right)\right) \exp\left(\frac{\mu_{\text{back},A}^{\text{old}} - \mu_{\text{back},A}^{\text{prop}}}{\beta_{\text{back},A}}\right) \\
&\quad \times \left(\frac{\mu_{\text{back},D}^{\text{old}}}{\mu_{\text{back},D}^{\text{prop}}}\right)^{2\alpha_{\text{back},D}^{\text{prop}} - \alpha_{\text{back},D}} \exp\left(\alpha_{\text{back},D}^{\text{prop}} \left(\frac{\mu_{\text{back},D}^{\text{prop}}}{\mu_{\text{back},D}^{\text{old}}} - \frac{\mu_{\text{back},D}^{\text{old}}}{\mu_{\text{back},D}^{\text{prop}}}\right)\right) \exp\left(\frac{\mu_{\text{back},D}^{\text{old}} - \mu_{\text{back},D}^{\text{prop}}}{\beta_{\text{back},D}}\right)
\end{aligned}
\tag{S204}
$$

#### S5.14. Maximum of posteriori (MAP) estimate

To be able to report the most probable values of our parameters of interest, we calculate the joint posterior of all parameters and report the values of the sampled parameters which coincide with the maximum posteriori (MAP) estimation as our output. So, to be able to calculate the MAP estimate we calculate the joint posterior of all of random variables

$$\mathbb{P}\left(\begin{matrix}\eta_A, \eta_D, \mu_{\text{back},A}, \mu_{\text{back},D}, \{\tau_{A,m_A}, \mu_{ex,m_A}\}_{m_A}, \{\tau_{D,m_D}, \mu_{ex,m_D}\}_{m_D} \\ , \{k_{F,m_F}\}_{m_F}, \overline{\overline{Q}}_A, \overline{\overline{Q}}_D, \overline{\overline{Q}}_F, \overline{S}_A, \overline{S}_D, \overline{S}_F, \overline{\zeta}_A, \overline{\zeta}_D, \overline{\zeta}_F\end{matrix}\middle| \overline{\delta}, \overline{C}\right). \tag{S205}$$

Also, to avoid any numerical problem, we consider the logarithm and simplify version of the above posterior

$$\begin{aligned}
&\log\mathbb{P}\left(\begin{matrix}\eta_A, \eta_D, \mu_{\text{back},A}, \mu_{\text{back},D}, \{\tau_{A,m_A}, \mu_{ex,m_A}\}_{m_A}, \{\tau_{D,m_D}, \mu_{ex,m_D}\}_{m_D} \\ , \{k_{F,m_F}\}_{m_F}, \overline{\overline{Q}}_A, \overline{\overline{Q}}_D, \overline{\overline{Q}}_F, \overline{S}_A, \overline{S}_D, \overline{S}_F, \overline{\zeta}_A, \overline{\zeta}_D, \overline{\zeta}_F\end{matrix}\middle| \overline{\delta}, \overline{C}\right) \\
&= \Bigg[\sum_{t=1}^{T} \log p\left(\delta_t | \eta_A, \eta_D, \mu_{\text{back},A}, \mu_{\text{back},D}, \{\tau_{A,m_A}, \mu_{ex,A,m_A}\}_{m_A}, \{\tau_{D,m_D}, \mu_{ex,D,m_D}\}_{m_D}, \{k_{F,m_F}\}_{m_F}, S_{A,t}, S_{D,t}, S_{F,t}, C_t\right) \\
&\quad + \log p\left(C_t | \eta_A, \eta_D, \mu_{\text{back},A}, \mu_{\text{back},D}, \{\mu_{ex,A,m_A}\}_{m_A}, \{\tau_{D,m_D}, \mu_{ex,D,m_D}\}_{m_D}, \{k_{F,m_F}\}_{m_F}, S_{A,t}, S_{D,t}, S_{F,t}\right)\Bigg] \\
&\quad + \log p\left(S_{A,1} | \overline{\zeta}_A\right) + \left[\sum_{t=2}^{T} \log \overline{\overline{P}}_{A,t}\left(S_{A,t} | S_{A,t-1}, \{\lambda_{A,m_A}\right)\right] \\
&\quad + \log p\left(S_{D,1} | \overline{\zeta}_D\right) + \left[\sum_{t=2}^{T} \log \overline{\overline{P}}_{D,t}\left(S_{D,t} | S_{D,t-1}, \{\lambda_{D,m_D}\right)\right] \\
&\quad + \log p\left(S_{F,1} | \overline{\zeta}_F\right) + \left[\sum_{t=2}^{T} \log \overline{\overline{P}}_{F,t}\left(S_{F,t} | S_{F,t-1}, \{\lambda_{F,m_F}\right)\right] \\
&\quad + \left[\sum_{m_A=1}^{M_A} \log p\left(\tau_{A,m_A}\right) + \log p\left(\mu_{ex,A,m_A}\right)\right] + \left[\sum_{m_D=1}^{M_D} \log p\left(\tau_{D,m_D}\right) + \log p\left(\mu_{ex,D,m_D}\right)\right] + \left[\sum_{m_F=1}^{M_F} \log p\left(K_{F,m_F}\right)\right] \\
&\quad + \left[\sum_{m_A=1}^{M_A(M_A-1)} \log p\left(\lambda_{A,m_A}\right)\right] + \left[\sum_{m_D=1}^{M_D(M_D-1)} \log p\left(\lambda_{D,m_D}\right)\right] + \left[\sum_{m_F=1}^{M_F(M_F-1)} \log p\left(\lambda_{F,m_F}\right)\right] \\
&\quad + \log p\left(\overline{\zeta}_A\right) + \log p\left(\overline{\zeta}_D\right) + \log p\left(\overline{\zeta}_F\right) \\
&\quad + \log p\left(\eta_A\right) + \log p\left(\eta_D\right) + \log p\left(\mu_{\text{back},A}\right) + \log p\left(\mu_{\text{back},D}\right)
\end{aligned} \tag{S206}$$

### S6. Summary of notation, abbreviations, parameters and other options

TABLE S3. Summary of notation.

| Description | Variable | Units |
|---|---|---|
| Micro time traces | $\overline{\delta}$ | ns |
| Channel detection | $\overline{C}$ | - |
| Inter-pulse time | $\Delta_p$ | ns |
| Acceptor and donor channels cross-talk | $\eta_A$ , $\eta_D$ | - , - |
| Number of acceptor lifetime (acceptor-PIFE) states | $M_A$ | - |
| Number of donor lifetime (donor-PIFE) states | $M_D$ | - |
| Number of energy transfer rate (FRET) states | $M_F$ | - |
| Acceptor and donor background photon emission rates | $\mu_{\text{back},A}$ , $\mu_{\text{back},D}$ | photons $s^{-1}$ |
| Acceptor and donor excitation rates per pulse and state | $\{\mu_{ex,A,m_A}\}_{m_A}$, $\{\mu_{ex,D,m_D}\}_{m_D}$ | - , - |
| Acceptor lifetimes | $\{\tau_{A,m_A}\}_{m_A}$ | ns |
| donor lifetimes | $\{\tau_{D,m_D}\}_{m_D}$ | ns |
| resonance energy transfer rates | $\{k_{F,m_F}\}_{m_F}$ | $ns^{-1}$ |
| Acceptor-PIFE state trajectory | $\overline{S}_A$ | - |
| DonorPIFE state trajectory | $\overline{S}_D$ | - |
| FRET state trajectory | $\overline{S}_F$ | - |
| Transition matrix of acceptor lifetime trajectory | $\overline{\overline{Q}}_A$ | $s^{-1}$ |
| Transition matrix of donor lifetime trajectory | $\overline{\overline{Q}}_D$ | $s^{-1}$ |
| Transition matrix of FRET lifetime trajectory | $\overline{\overline{Q}}_F$ | $s^{-1}$ |
| Weights on the initial acceptor-PIFE state | $\overline{\zeta}_A$ | - |
| Weights on the initial donor-PIFE state | $\overline{\zeta}_D$ | - |
| Weights on the initial FRET state | $\overline{\zeta}_F$ | - |
| $\alpha$ and $\beta$ parameter of the acceptor excitation rate per pulse's prior | $\alpha_{\mu_A}$ , $\beta_{\mu_A}$ | - , - |
| $\alpha$ and $\beta$ parameter of the donor excitation rate per pulse's prior | $\alpha_{\mu_D}$ , $\beta_{\mu_D}$ | - , - |
| $\alpha$ and $\beta$ parameters of the acceptor background emission rate's prior | $\alpha_{\text{back},A}$ , $\beta_{\text{back},A}$ | - , photons $s^{-1}$ |
| $\alpha$ and $\beta$ parameters of the donor background emission rate's prior | $\alpha_{\text{back},D}$ , $\beta_{\text{back},D}$ | -, photons $s^{-1}$ |
| $\alpha$ and $\beta$ parameters of the acceptor lifetimes' prior | $\alpha_A$ , $\beta_A$ | - , ns |
| $\alpha$ and $\beta$ parameters of the donor lifetimes' prior | $\alpha_D$ , $\beta_D$ | - , ns |
| $\alpha$ and $\beta$ parameters of the energy transfer rates' prior | $\alpha_F$ , $\beta_F$ | - , $ns^{-1}$ |
| $\alpha$ and $\beta$ parameters of the acceptor-PIFE state escape rates' prior | $\alpha_{\lambda,A}$ , $\beta_{\lambda,A}$ | - , $s^{-1}$ |
| $\alpha$ and $\beta$ parameters of the donor-PIFE state escape rates' prior | $\alpha_{\lambda,D}$ , $\beta_{\lambda,D}$ | - , $s^{-1}$ |
| $\alpha$ and $\beta$ parameters of the FRET state escape rates' prior | $\alpha_{\lambda,F}$ , $\beta_{\lambda,F}$ | - , $s^{-1}$ |
| $\alpha$ and $\beta$ parameters of the weights on acceptor states' prior | $\alpha_{A,\zeta_A}$ , $\overline{\beta}_{A,hyper}$ | - , - |
| $\alpha$ and $\beta$ parameter of the weights on donor states' prior | $\alpha_{D,\zeta_D}$ , $\overline{\beta}_{D,hyper}$ | - , - |
| $\alpha$ and $\beta$ parameter of the weights on FRET states' prior | $\alpha_{F,\zeta_F}$ , $\overline{\beta}_{F,hyper}$ | - , - |
| $\alpha$ and $\beta$ parameter of acceptor channel cross-talk's prior | $\alpha_{\eta,A}$ , $\beta_{\eta,A}$ | - , - |
| $\alpha$ and $\beta$ parameter of donor channel cross-talk's prior | $\alpha_{\eta,D}$ , $\beta_{\eta,D}$ | - , - |

TABLE S4. List of abbreviations.

| Phrase | Abbreviation |
|---|---|
| Metropolis Hastings | MH |
| Markov Chain Monte Carlo | MCMC |
| Hidden Markov Model | HMM |
| Infinite Hidden Markov Model | iHMM |
| Forward Filtering Backward Sampling | FFBS |
| Förster Resonance Energy Transfer | FRET |
| Protein Induced Fluorescence Enhancement | PIFE |
| Instrument Response Function | IRF |
| Gaussian Process | GP |
| Dirichlet process | DP |
| Griffiths, Engen and McCloskey | GEM |
| Time correlated single photon counting | TCSPC |

TABLE S5. Probability distributions used and their densities. Here, the corresponding random variables are denoted by $x$.

| Distribution | Notation | Probability density function | Mean value | Variance/Covariance |
|---|---|---|---|---|
| Normal | $\mathbf{Normal}(\mu, \sigma^2)$ | $\frac{1}{\sqrt{2\pi\sigma^2}} e^{-\frac{(x-\mu)^2}{2\sigma^2}}$ | $\mu$ | $\sigma^2$ |
| Exponential | $\mathbf{Exp}(\mu)$ | $\mu e^{-\mu x}$ | $\mu^{-1}$ | $\mu^{-2}$ |
| Gamma | $\mathbf{Gamma}(\alpha, \beta)$ | $\frac{1}{\Gamma(\alpha)\beta^\alpha} x^{\alpha-1} e^{-\frac{x}{\beta}}$ | $\alpha\beta$ | $\alpha\beta^2$ |
| Beta | $\mathbf{Beta}(\alpha, \beta)$ | $\frac{\Gamma(\alpha+\beta)}{\Gamma(\alpha)\Gamma(\beta)} x^{\alpha-1}(1-x)^{\beta-1}$ | $\frac{\alpha}{\alpha+\beta}$ | $\frac{\alpha\beta}{(\alpha+\beta)^2(\alpha+\beta+1)}$ |
| Dirichlet | $\mathbf{Dir}(\alpha, \overline{\beta})$ | $\frac{\Gamma\left(\sum_i \alpha\beta_i\right)}{\prod_i \Gamma(\alpha\beta_i)} \prod_i x^{\alpha\beta_i - 1}$ | $\frac{\beta_i}{\sum_i \beta_i}$ | $\frac{\left(\frac{\beta_i}{\sum_i \beta_i}\right)\left(1-\frac{\beta_i}{\sum_i \beta_i}\right)}{\sum_i \alpha\beta_i + 1}$ |

TABLE S6. Parameter values used in the generation of the synthetic traces. Choices are listed according to figures.

| | $\begin{bmatrix}\eta_A\\ \eta_D\end{bmatrix}$ | $\begin{bmatrix}\mu_{\mathrm{back},A}\\ \mu_{\mathrm{back},D}\end{bmatrix}$ | $\begin{bmatrix}M_A\\ M_D\\ M_F\end{bmatrix}$ | $\{\tau_{A,m_A}\}_{m_A}$ | $\{\tau_{A,m_A}\}_{m_A}$ | $\{k_{F,m_F}\}_{m_F}$ | $\{\mu_{ex,A,m_A}\}_{m_A}$ | $\{\mu_{ex,D,m_D}\}_{m_D}$ |
|---|---|---|---|---|---|---|---|---|
| Units | - | phs $s^{-1}$ | - | ns | ns | $ns^{-1}$ | - | - |
| Fig. 3(a1) | $\begin{bmatrix}0.1\\0.1\end{bmatrix}$ | $\begin{bmatrix}500\\500\end{bmatrix}$ | $\begin{bmatrix}1\\1\\2\end{bmatrix}$ | 3.8 | 4 | $\begin{bmatrix}0.1\\1\end{bmatrix}$ | $10^{-5}$ | $2\times10^{-3}$ |
| Fig. 3(a2) | $\begin{bmatrix}0.1\\0.1\end{bmatrix}$ | $\begin{bmatrix}500\\500\end{bmatrix}$ | $\begin{bmatrix}1\\1\\2\end{bmatrix}$ | 3.8 | 4 | $\begin{bmatrix}0.1\\1\end{bmatrix}$ | $10^{-5}$ | $2\times10^{-3}$ |
| Fig. 3(a3) | $\begin{bmatrix}0.1\\0.1\end{bmatrix}$ | $\begin{bmatrix}500\\500\end{bmatrix}$ | $\begin{bmatrix}1\\1\\2\end{bmatrix}$ | 3.8 | 4 | $\begin{bmatrix}0.1\\1\end{bmatrix}$ | $10^{-5}$ | $2\times10^{-3}$ |
| Fig. 4(a1) | $\begin{bmatrix}0.1\\0.1\end{bmatrix}$ | $\begin{bmatrix}500\\500\end{bmatrix}$ | $\begin{bmatrix}2\\2\\2\end{bmatrix}$ | $\begin{bmatrix}0.1\\1\end{bmatrix}$ | $\begin{bmatrix}2\\4\end{bmatrix}$ | $\begin{bmatrix}0.1\\0.5\end{bmatrix}$ | $\begin{bmatrix}1\\2\end{bmatrix}\times10^{-5}$ | $\begin{bmatrix}1.5\\3\end{bmatrix}\times10^{-3}$ |
| Fig. 4(a2) | $\begin{bmatrix}0.1\\0.1\end{bmatrix}$ | $\begin{bmatrix}500\\500\end{bmatrix}$ | $\begin{bmatrix}3\\3\\3\end{bmatrix}$ | $\begin{bmatrix}0.1\\1\end{bmatrix}$ | $\begin{bmatrix}2\\4\end{bmatrix}$ | $\begin{bmatrix}0.5\\3\end{bmatrix}$ | $\begin{bmatrix}1\\2\\3\end{bmatrix}\times10^{-5}$ | $\begin{bmatrix}1.5\\3\\4.5\end{bmatrix}\times10^{-3}$ |
| Fig. 4(a3) | $\begin{bmatrix}0.1\\0.1\end{bmatrix}$ | $\begin{bmatrix}500\\500\end{bmatrix}$ | $\begin{bmatrix}4\\4\\4\end{bmatrix}$ | $\begin{bmatrix}1.1\\2.2\\3.3\\4.4\end{bmatrix}$ | $\begin{bmatrix}1\\2\\3\\4\end{bmatrix}$ | $\begin{bmatrix}0.1\\0.5\\1\\3\end{bmatrix}$ | $\begin{bmatrix}1\\2\\3\\4\end{bmatrix}\times10^{-5}$ | $\begin{bmatrix}2\\4\\6\\8\end{bmatrix}\times10^{-3}$ |
| Fig. S1 | $\begin{bmatrix}0.05\\0.25\end{bmatrix}$ | $\begin{bmatrix}500\\500\end{bmatrix}$ | $\begin{bmatrix}1\\1\\2\end{bmatrix}$ | 3.8 | 2.5 | $\begin{bmatrix}0.2\\5\end{bmatrix}$ | $10^{-5}$ | $3\times10^{-3}$ |

TABLE S7. Parameter values used in the generation of the synthetic traces. Choices are listed according to figures.

| | $\overline{\overline{Q}}_A$ | $\overline{\overline{Q}}_D$ | $\overline{\overline{Q}}_F$ | $T_p$ | T |
|---|---|---|---|---|---|
| Units | $s^{-1}$ | $s^{-1}$ | $s^{-1}$ | ns | - |
| Fig. 3(a1) | - | - | $\begin{bmatrix} -10 & 10 \\ 20 & -20 \end{bmatrix}$ | 50 | $10^4$ |
| Fig. 3(a2) | - | - | $\begin{bmatrix} -100 & 100 \\ 2000 & -200 \end{bmatrix}$ | 50 | $10^4$ |
| Fig. 3(a3) | - | - | $\begin{bmatrix} -1000 & 1000 \\ 2000 & -2000 \end{bmatrix}$ | 50 | $10^4$ |
| Fig. 4(a1) | $\begin{bmatrix} -10 & 50 \\ 10 & -50 \end{bmatrix}$ | $\begin{bmatrix} -20 & 70 \\ 20 & -70 \end{bmatrix}$ | $\begin{bmatrix} -80 & 30 \\ 80 & -30 \end{bmatrix}$ | 50 | $2\times10^7$ |
| Fig. 4(a2) | $\begin{bmatrix} -40 & 50 & 60 \\ 10 & -80 & 20 \\ 30 & 30 & -80 \end{bmatrix}$ | $\begin{bmatrix} -60 & 70 & 10 \\ 10 & -80 & 50 \\ 50 & 10 & -60 \end{bmatrix}$ | $\begin{bmatrix} -120 & 130 & 60 \\ 80 & -80 & 50 \\ 40 & 50 & -110 \end{bmatrix}$ | 50 | $2\times10^7$ |
| Fig. 4(a3) | $\begin{bmatrix} -90 & 50 & 60 & 10 \\ 10 & -90 & 20 & 50 \\ 30 & 30 & -110 & 20 \\ 50 & 10 & 30 & -80 \end{bmatrix}$ | $\begin{bmatrix} -120 & 70 & 10 & 20 \\ 10 & -110 & 50 & 60 \\ 50 & 10 & -90 & 20 \\ 60 & 30 & 30 & -100 \end{bmatrix}$ | $\begin{bmatrix} -140 & 30 & 60 & 80 \\ 80 & -100 & 50 & 20 \\ 40 & 50 & -140 & 10 \\ 20 & 20 & 30 & -110 \end{bmatrix}$ | 50 | $2\times10^7$ |
| Fig. S1 | - | - | $\begin{bmatrix} -250 & 250 \\ 450 & -450 \end{bmatrix}$ | 50 | $1.65\times10^7$ |

TABLE S8. Parameter values used in the analyses of the traces. Choices are listed according to figures.

| | $\alpha_F$ | $\beta_F$ | $\alpha_D$ | $\beta_D$ | $\alpha_A$ | $\beta_A$ | $\alpha_{\lambda,F}$ | $\beta_{\lambda,F}$ | $\alpha_{\lambda,D}$ | $\beta_{\lambda,D}$ | $\alpha_{\lambda,A}$ | $\beta_{\lambda,A}$ | $\alpha_{\eta,D}$ | $\beta_{\eta,D}$ | $\alpha_{\eta,A}$ | $\beta_{\eta,A}$ |
|---|---|---|---|---|---|---|---|---|---|---|---|---|---|---|---|---|
| Units | - | $ns^{-1}$ | - | ns | - | ns | - | s | - | s | - | s | - | - | - | - |
| Fig. 3(a1) | 1 | 10 | 1 | 10 | 1 | 10 | 1 | $10^3$ | 1 | $10^3$ | 1 | $10^3$ | 1 | 50 | 1 | 50 |
| Fig. 3(a2) | 1 | 10 | 1 | 10 | 1 | 10 | 1 | $10^3$ | 1 | $10^3$ | 1 | $10^3$ | 1 | 50 | 1 | 50 |
| Fig. 3(a3) | 1 | 10 | 1 | 10 | 1 | 10 | 1 | $10^3$ | 1 | $10^3$ | 1 | $10^3$ | 1 | 50 | 1 | 50 |
| Fig. 4(a1) | 1 | 10 | 1 | 10 | 1 | 10 | 1 | $10^3$ | 1 | $10^3$ | 1 | $10^3$ | 1 | 50 | 1 | 50 |
| Fig. 4(a2) | 1 | 10 | 1 | 10 | 1 | 10 | 1 | $10^3$ | 1 | $10^3$ | 1 | $10^3$ | 1 | 50 | 1 | 50 |
| Fig. 4(a3) | 1 | 10 | 1 | 10 | 1 | 10 | 1 | $10^3$ | 1 | $10^3$ | 1 | $10^3$ | 1 | 50 | 1 | 50 |
| Fig. S1($5\times10^2$) | 1 | 10 | 1 | 10 | 1 | 10 | 1 | $10^3$ | 1 | $10^3$ | 1 | $10^3$ | 1 | 50 | 1 | 50 |
| Fig. S1($10^3$) | 1 | 10 | 1 | 10 | 1 | 10 | 1 | $10^3$ | 1 | $10^3$ | 1 | $10^3$ | 1 | 50 | 1 | 50 |
| Fig. S1($5\times10^3$) | 1 | 10 | 1 | 10 | 1 | 10 | 1 | $10^3$ | 1 | $10^3$ | 1 | $10^3$ | 1 | 50 | 1 | 50 |
| Fig. S1($10^4$) | 1 | 10 | 1 | 10 | 1 | 10 | 1 | $10^3$ | 1 | $10^3$ | 1 | $10^3$ | 1 | 50 | 1 | 50 |
| Fig. S1($5\times10^4$) | 1 | 10 | 1 | 10 | 1 | 10 | 1 | $10^3$ | 1 | $10^3$ | 1 | $10^3$ | 1 | 50 | 1 | 50 |
| Fig. S2($5\times10^2$) | 1 | 10 | 1 | 10 | 1 | 10 | 1 | $10^3$ | 1 | $10^3$ | 1 | $10^3$ | 1 | 50 | 1 | 50 |
| Fig. S2($10^3$) | 1 | 10 | 1 | 10 | 1 | 10 | 1 | $10^3$ | 1 | $10^3$ | 1 | $10^3$ | 1 | 50 | 1 | 50 |
| Fig. S2($5\times10^3$) | 1 | 10 | 1 | 10 | 1 | 10 | 1 | $10^3$ | 1 | $10^3$ | 1 | $10^3$ | 1 | 50 | 1 | 50 |
| Fig. S2($10^4$) | 1 | 10 | 1 | 10 | 1 | 10 | 1 | $10^3$ | 1 | $10^3$ | 1 | $10^3$ | 1 | 50 | 1 | 50 |
| Fig. S2($5\times10^4$) | 1 | 10 | 1 | 10 | 1 | 10 | 1 | $10^3$ | 1 | $10^3$ | 1 | $10^3$ | 1 | 50 | 1 | 50 |

TABLE S9. Parameter values used in the analyses of the traces. Choices are listed according to figures.

| | $\alpha_{F,\zeta_F}$ | $\overline{\beta}_{F,hyper}$ | $\alpha_{D,\zeta_D}$ | $\overline{\beta}_{D,hyper}$ | $\alpha_{A,\zeta_A}$ | $\overline{\beta}_{A,hyper}$ | $\alpha_{\mu_D}$ | $\beta_{\mu_D}$ | $\alpha_{\mu_A}$ | $\beta_{\mu_A}$ | $\alpha_{\mu_{\text{back},D}}$ | $\beta_{\mu_{\text{back},D}}$ | $\alpha_{\mu_{\text{back},A}}$ | $\beta_{\mu_{\text{back},A}}$ |
|---|---|---|---|---|---|---|---|---|---|---|---|---|---|---|
| Units | - | - | - | - | - | - | - | - | - | - | - | phts $s^{-1}$ | - | phts $s^{-1}$ |
| Fig. 3(a1) | 1 | $\begin{bmatrix} 0.5 \\ 0.5 \end{bmatrix}$ | 1 | 1 | 1 | 1 | 1 | $10^{-2}$ | 1 | $10^{-3}$ | 1 | $10^{-3}$ | 1 | $10^{-3}$ |
| Fig. 3(a2) | 1 | $\begin{bmatrix} 0.5 \\ 0.5 \end{bmatrix}$ | 1 | 1 | 1 | 1 | 1 | $10^{-2}$ | 1 | $10^{-3}$ | 1 | $10^{-3}$ | 1 | $10^{-3}$ |
| Fig. 3(a3) | 1 | $\begin{bmatrix} 0.5 \\ 0.5 \end{bmatrix}$ | 1 | 1 | 1 | 1 | 1 | $10^{-2}$ | 1 | $10^{-3}$ | 1 | $10^{-3}$ | 1 | $10^{-3}$ |
| Fig. 4(a1) | 1 | $\begin{bmatrix} 0.5 \\ 0.5 \end{bmatrix}$ | 1 | $\begin{bmatrix} 0.5 \\ 0.5 \end{bmatrix}$ | 1 | $\begin{bmatrix} 0.5 \\ 0.5 \end{bmatrix}$ | 1 | $10^{-2}$ | 1 | $10^{-3}$ | 1 | $10^{-3}$ | 1 | $10^{-3}$ |
| Fig. 4(a2) | 1 | $\begin{bmatrix} 0.33 \\ 0.33 \\ 0.33 \end{bmatrix}$ | 1 | $\begin{bmatrix} 0.33 \\ 0.33 \\ 0.33 \end{bmatrix}$ | 1 | $\begin{bmatrix} 0.33 \\ 0.33 \\ 0.33 \end{bmatrix}$ | 1 | $10^{-2}$ | 1 | $10^{-3}$ | 1 | $10^{-3}$ | 1 | $10^{-3}$ |
| Fig. 4(a3) | 1 | $\begin{bmatrix} 0.25 \\ 0.25 \\ 0.25 \\ 0.25 \end{bmatrix}$ | 1 | $\begin{bmatrix} 0.25 \\ 0.25 \\ 0.25 \\ 0.25 \end{bmatrix}$ | 1 | $\begin{bmatrix} 0.25 \\ 0.25 \\ 0.25 \\ 0.25 \end{bmatrix}$ | 1 | $10^{-2}$ | 1 | $10^{-3}$ | 1 | $10^{-3}$ | 1 | $10^{-3}$ |
| Fig. S2($5\times10^2$) | 1 | $\begin{bmatrix} 0.5 \\ 0.5 \end{bmatrix}$ | 1 | 1 | 1 | 1 | 1 | $10^{-2}$ | 1 | $10^{-3}$ | 1 | $10^{-3}$ | 1 | $10^{-3}$ |
| Fig. S2($10^3$) | 1 | $\begin{bmatrix} 0.5 \\ 0.5 \end{bmatrix}$ | 1 | 1 | 1 | 1 | 1 | $10^{-2}$ | 1 | $10^{-3}$ | 1 | $10^{-3}$ | 1 | $10^{-3}$ |
| Fig. S2($5\times10^3$) | 1 | $\begin{bmatrix} 0.5 \\ 0.5 \end{bmatrix}$ | 1 | 1 | 1 | 1 | 1 | $10^{-2}$ | 1 | $10^{-3}$ | 1 | $10^{-3}$ | 1 | $10^{-3}$ |
| Fig. S2($10^4$) | 1 | $\begin{bmatrix} 0.5 \\ 0.5 \end{bmatrix}$ | 1 | 1 | 1 | 1 | 1 | $10^{-2}$ | 1 | $10^{-3}$ | 1 | $10^{-3}$ | 1 | $10^{-3}$ |
| Fig. S2($5\times10^4$) | 1 | $\begin{bmatrix} 0.5 \\ 0.5 \end{bmatrix}$ | 1 | 1 | 1 | 1 | 1 | $10^{-2}$ | 1 | $10^{-3}$ | 1 | $10^{-3}$ | 1 | $10^{-3}$ |

### Supplementary References


[1] Robert, C. P., Casella, G. & Casella, G. *Introducing monte carlo methods with r*, vol. 18 (Springer, 2010).

[2] Gelman, A., Hwang, J. & Vehtari, A. Understanding predictive information criteria for bayesian models. *Statistics and computing* **24**, 997–1016 (2014).

[3] Lee, A., Tsekouras, K., Calderon, C., Bustamante, C. & Pressé, S. Unraveling the thousand word picture: an introduction to super-resolution data analysis. *Chemical reviews* **117**, 7276–7330 (2017).

[4] Tavakoli, M., Taylor, J. N., Li, C.-B., Komatsuzaki, T. & Pressé, S. Single molecule data analysis: An introduction. *arXiv preprint arXiv:1606.00403* (2016).

[5] Von Toussaint, U. Bayesian inference in physics. *Reviews of Modern Physics* **83**, 943 (2011).

[6] Liu, H. & Motoda, H. *Computational methods of feature selection* (CRC Press, 2007).

[7] Cappé, O., Moulines, E. & Rydén, T. Inference in hidden markov models. In *Proceedings of EUSFLAT Conference*, 14–16 (2009).

[8] Sgouralis, I. & Pressé, S. An introduction to infinite hmms for single-molecule data analysis. *Biophysical Journal* **112**, 2021–2029 (2017).

[9] Rydén, T. Em versus markov chain monte carlo for estimation of hidden markov models: A computational perspective. *Bayesian Analysis* **3**, 659–688 (2008).
